# Comparative Modeling of CDK9 Inhibitors to Explore Selectivity and Structure-Activity Relationships

**DOI:** 10.1101/2020.06.08.138602

**Authors:** Palani Kirubakaran, George Morton, Pingfeng Zhang, Hanghang Zhang, John Gordon, Magid Abou-Gharbia, Jean-Pierre J. Issa, Jinhua Wu, Wayne Childers, John Karanicolas

**Affiliations:** Program in Molecular Therapeutics, Fox Chase Cancer Center, Philadelphia, PA 19111-2497; Moulder Center for Drug Discovery Research, Temple University School of Pharmacy, Philadelphia, PA 19140; Fels Institute for Cancer Research, Temple University School of Medicine, Philadelphia, PA 19140; Coriell Institute for Medical Research, Camden, NJ 08103

## Abstract

Cyclin-dependent kinase 9 (CDK9) plays a key role in transcription elongation, and more recently it was also identified as the molecular target of a series of diaminothiazole compounds that reverse epigenetic silencing in a phenotypic assay. To better understand the structural basis underlying these compounds’ activity and selectivity, we developed a comparative modeling approach that we describe herein. Briefly, this approach draws upon the strong structural conservation across the active conformation of all protein kinases, and their shared pattern of interactions with Type I inhibitors. Because of this, we hypothesized that the large collection of inhibitor/kinase structures available in the Protein Data Bank (PDB) would enable accurate modeling of this diaminothiazole series in complex with each CDK family member. We apply this new comparative modeling pipeline to build each of these structural models, and then demonstrate that these models provide retrospective rationale for the structure-activity relationships that ultimately guided optimization to the lead diaminothiazole compound MC180295 (**14e**). We then solved the crystal structure of the **14e**/CDK9 complex, and found the resulting structure to be in excellent agreement with our corresponding comparative model. Finally, inspired by these models, we demonstrate how structural changes to **14e** can be used to rationally tune this compound’s selectivity profile. With the emergence of CDK9 as a promising target for therapeutic intervention in cancer, we anticipate that comparative modeling can provide a valuable tool to guide optimization of potency and selectivity of new inhibitors targeting this kinase.

## Introduction

Cyclin-dependent kinases (CDKs) belong to the Ser/Thr subfamily of protein kinases, and include a number of emerging therapeutic targets for many cancers [1]. CDKs are activated upon binding of a cyclin partner protein, leading to the name originally ascribed to this family [2]. The human genome encodes 20 CDK kinases [3]: these are often divided into regulators of the cell cycle (CDKs 1-7 and 14-18 and regulators of transcription (CDKs 8-13 and 19-20), though certain CDKs exhibit both activities [4]. Beyond these two primary functions, a number of additional diverse roles for CDKs have also emerged in metabolism, differentiation, stem cell self-renewal, and epigenetic regulation [5]. The catalytic domain of CDKs are extremely structurally conserved, particularly in the ATP-binding site, which presents a challenge for development of selective inhibitors [6, 7]. Despite this hurdle, three compounds targeting CDK4/6 are currently approved for treatment of advanced breast cancer [8] and many more CDK inhibitors are in various stages of clinical trials [1].

The transcriptional regulator CDK9 has been implicated as a potential target for therapeutic intervention in cancer, because it mediates transcription of certain anti-apoptotic proteins and can be necessary for survival of transformed cells [9]. CDK9 can be activated by four distinct cyclins (T1, T2a, T2b, and K) to form one of four positive transcription elongation factor (P-TEFb) complexes that regulate RNA polymerase II [10]. CDK9 is found to be overexpressed in a variety of cancers including pancreatic [11], ovarian [12], cervical [13], esophageal [14] and osteosarcoma [15], with increased CDK9 correlated to poorer patient prognosis [11,15,12]. From *in vitro* studies and animal models, CDK9 has been implicated as a potential target for therapeutic intervention due to its role in MYC stabilization [16], chemo- and radio-resistance [17–19], and maintenance of cancer cell stemness [20, 21]. While the lack of selectivity in most current CDK9 inhibitors has led to some trepidation regarding their use in the clinic [22], nonetheless a number of clinical candidates targeting CDK9 are being advanced in clinical trials addressing blood cancers, melanoma, glioblastoma, breast cancer, and other advanced solid tumors (clinicaltrials.gov).

While a subset of tumors can arise and progress through a very specific and prescribed set of oncogenic mutations, most notably as elucidated in colorectal cancer [23], other cancers can be completely devoid of established driver gene mutations [24]. Aberrant signaling that underlies oncogenesis in these cases may be driven by epigenetic alterations, such as silencing tumor suppressor genes, rather than by genetic alterations [25]. Epigenetic reprogramming can also lead to drug resistance by shifting the transcriptional state of the cancer cells to avoid reliance on the signaling pathway targeted by a drug [26]; the same epigenetic effect can also be manifest through immune avoidance in response to cancer immunotherapy [27]. Based on this rationale, we envisioned that pharmacologically reversing cancer-associated epigenetic alterations could serve as the basis for new cancer therapeutics.

To this end, we recently carried out a phenotypic screen in search of compounds that would reverse epigenetic silencing of a GFP reporter gene. Four structurally similar aminothiazole-containing hits were obtained (**Figure 1, 1-4**). The hits were characterized by a 3,4-dimethoxyphenyl ketone or homologous benzodioxan group on one side of the scaffold, and aromatic or aliphatic amino moieties on the other side. Upon optimization of the resulting hit compounds and deconvolution of the therapeutic mechanism, we discovered that their molecular target was CDK9 [28]. Our most promising lead compound, MC180295 (**Figure 1, 14e**), proved both potent and selective for CDK9, and showed exciting anti-tumoral activity coupled with immunosensitization [28]. With respect to selectivity, the preference of **14e** for CDK9 over CDK2 and CDK7 was especially notable: due to their structural and functional similarities, most CDK9 inhibitors also inhibit CDK2 and CDK7 [29, 30]. In our first study, we explored selectivity of **14e** by developing a comparative modeling approach to build a model of the **14e**/CDK9 complex, and then using this model to propose a structural basis for the observed selectivity [28].

**Figure 1:**
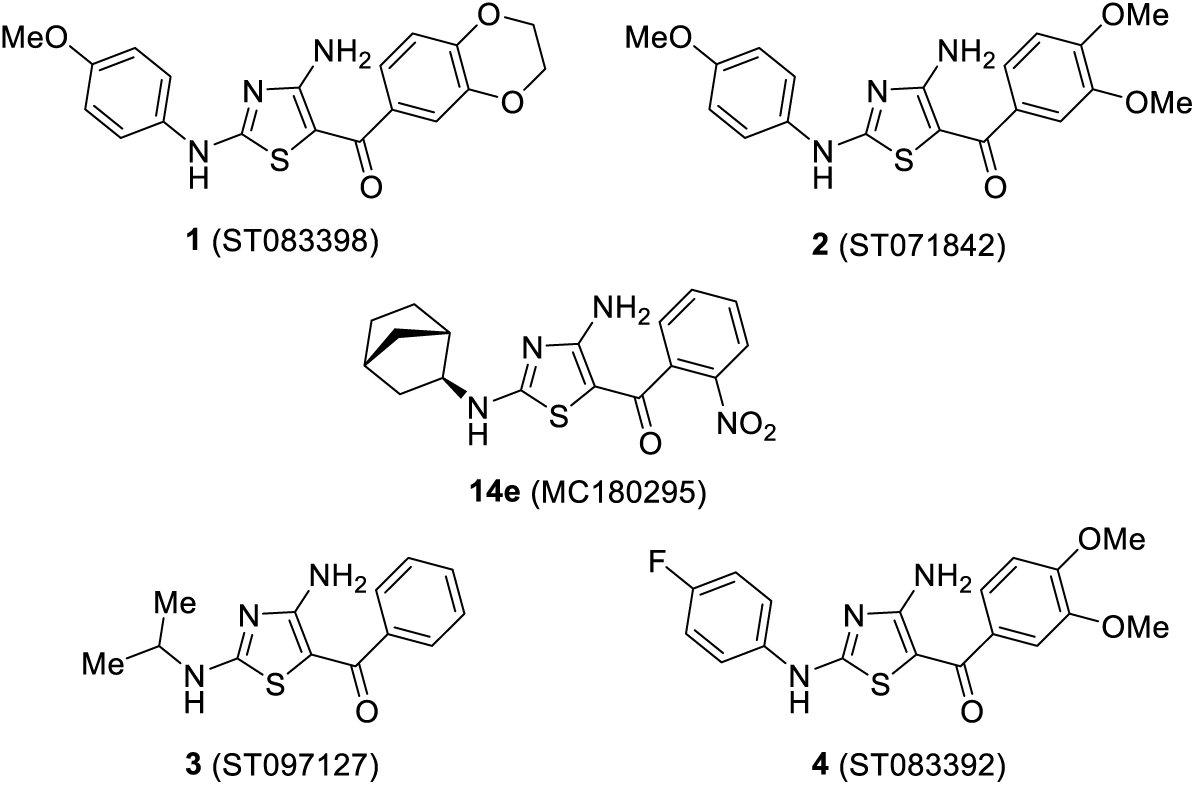
Aminothiazole-containing hits obtained from phenotypic high throughput screen (**1-4**), which ultimately led to our development of CDK9 inhibitor MC180295 (**14e**).

In the present study we expand our understanding of the structure-activity relationship (SAR) of the series and use these data to critically evaluate our comparative modeling approach. In particular, we present the SAR of 77 compounds related to **14e**, and retrospectively assess the extent to which comparative modeling could have driven optimization. We also characterize **14e** selectivity further, in light of a recent study demonstrating the surprising lack of selectivity in compounds ostensibly selective for other CDKs [6], and a report summarizing the broad selectivity profiles of CDK9 inhibitors that have advanced to clinical trials [22]. Finally, to definitively evaluate the accuracy of our comparative modeling for this application, we solve the crystal structure of **14e** in complex with CDK9.

## Results and Discussion

Our previous study culminated with the identification and characterization of **14e** (**Figure 1**) as our lead compound. In the course of our earlier study, we developed a preliminary comparative modeling approach in order to build a tentative model of this compound in complex with CDK9 [28]. In the present study, to understand the SAR around **14e** and to further explore this compound’s selectivity, we report a more sophisticated comparative modeling approach and we use it to build models of additional compounds in complex with multiple CDK kinases. Among other enhancements, we have expanded the space of available templates from the PDB that are used in our comparative approach: whereas previously we used only structures of inhibitors bound to CDK kinases, we now use structures of inhibitors bound to *any* protein kinase found in its active conformation.

The model of **14e** in complex with CDK9 from our updated protocol (**Figure 2**) is largely consistent with the model resulting from our earlier approach [28]. Interestingly, the structural template that led to our lowest-energy model (and thus our final prediction of the complex) comes from a crystal structure of CDK9 in complex with an inhibitor that is at first glance quite different from **14e** in terms of chemical structure [31] (**Figure 2c**). The power of our comparative modeling approach, described below in more detail, is that templates can be identified on the basis of three-dimensional inhibitor similarity. By finding templates that engage their cognate kinase’s ATP-binding site using similar interactions, instead of simply searching for shared core chemical structures, we vastly broaden the chemical space of inhibitors that can be modeled using the (fixed) space of available templates in the Protein Data Bank (PDB).

**Figure 2:**
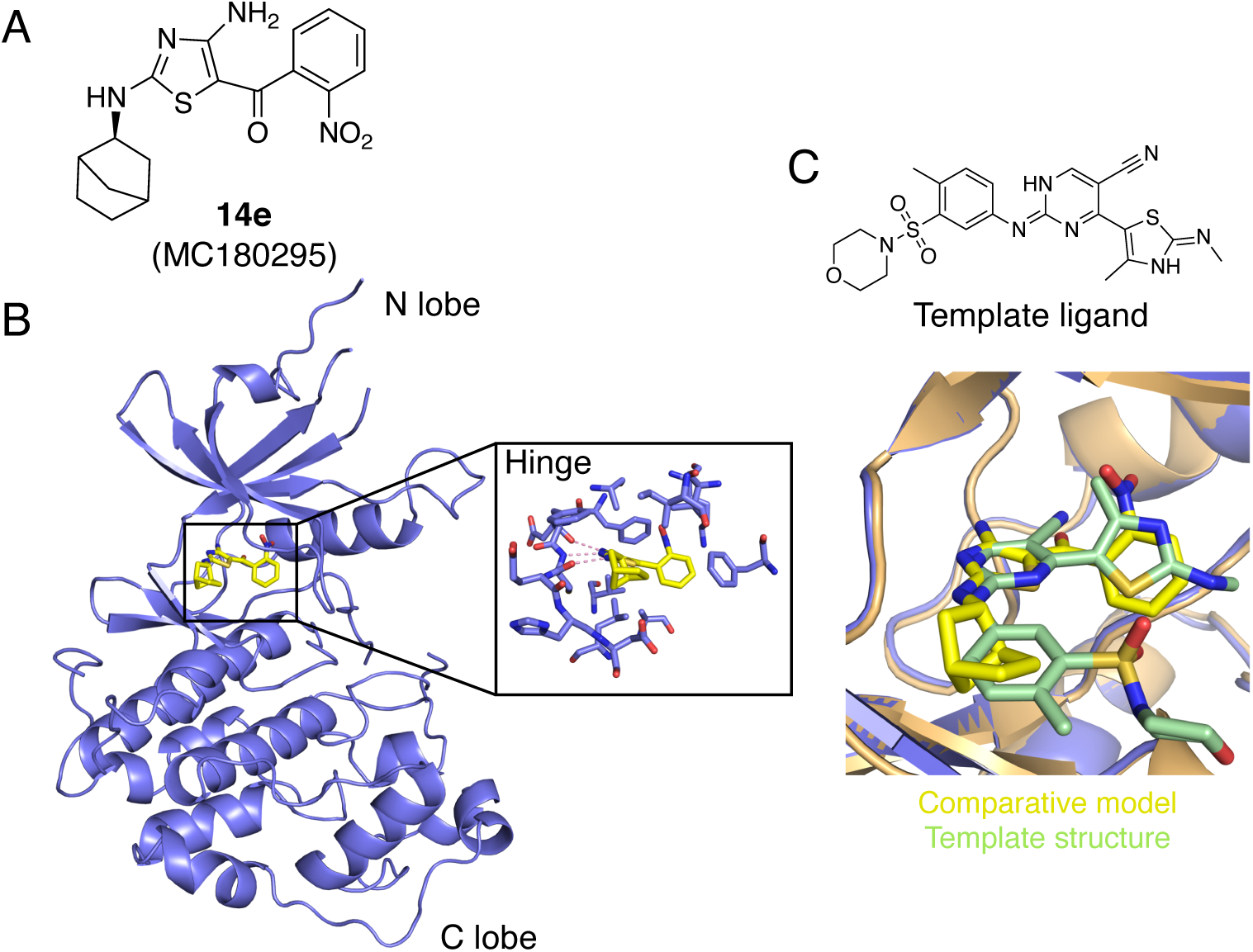
**Optimized CDK9 inhibitor MC180295. (A)** The chemical structure of MC180295. **(B)** Comparative model of the MC180295/CDK9 complex. **(C)** This comparative model was built from the crystal structure of a different inhibitor in complex with CDK9. Despite their dramatically different chemical scaffolds, the ligand in the template structure (PDB ID 4BCH) leads to a model in which MC180295 engages CDK9 through very similar interactions in the binding site.

### Overview of the comparative modeling pipeline

At present, the PDB includes 494 publicly-available structures of CDK kinase catalytic domains. Modern studies typically build models of inhibitor-bound structures via docking: sampling all possible locations and orientations for the inhibitor, then selecting the best-scoring binding mode from among these many possibilities [32–40]. In contrast, and by analogy to protein structure prediction, we anticipated that the PDB’s continuously growing body of structural information could serve as the basis for building more accurate models via comparative modeling. With respect to modeling Type I kinase inhibitors in particular, it has long been recognized that – by virtue of mimicking the interactions of ATP in a highly conserved binding site – the interactions used by many chemically-diverse scaffolds can be strikingly similar to one another [41, 42]. This underscores the applicability of comparative modeling for predicting the structures of new Type I inhibitors in complex with their cognate kinases. To better explore the SAR and selectivity of **14e**, we therefore developed a pipeline for modeling the structures of arbitrary inhibitors in complex with CDK kinases. We have also benchmarked this approach beyond the CDK family, but will report those results in the context of a separate study. Our approach is summarized below, and complete detail is provided in the *Experimental Section*.

In preparation for this comparative modeling approach, we began by compiling from the PDB all kinase domains in the active conformation [43], and then restricted this set to those 1732 entries that contain a ligand in the ATP-binding site: these complexes were aligned into a shared reference frame, to serve as templates for comparative modeling. From here, our computational pipeline for modeling a given query inhibitor (e.g., **14e**) bound to a given kinase (e.g., CDK9) comprises three steps (**Figure 3**). First, low-energy conformations (“conformers”) of the query inhibitor are built and individually aligned onto the structures of the template ligands in the PDB on the basis of shape and shared chemical features: this provides a set of initial guesses as to the query inhibitor’s position and orientation in the binding site, and those 100 with the closest matches between query and template are retained. Second, the protein templates are screened on the basis of sequence similarity relative to the query kinase and three-dimensional structural similarity of the bound inhibitor, and those 10 with closest similarity are retained. The protein structures of these closest matches are then updated to reflect the sequence of the query kinase. Together, these two steps yield a set of 1000 crude models of the desired inhibitor/kinase pairing: the third stage of our pipeline entails refinement to a final model. Rather than carry out refinement from all 1000 crude models, we instead started from the single closest protein template, and carried out energy minimization from each of the top 100 inhibitor poses. The 10 top-scoring poses from this step were paired with each of the top 10 protein templates, and these 100 models were subjected to energy minimization. The top-scoring model from this set was taken to be our final predicted structure of the desired complex.

**Figure 3:**
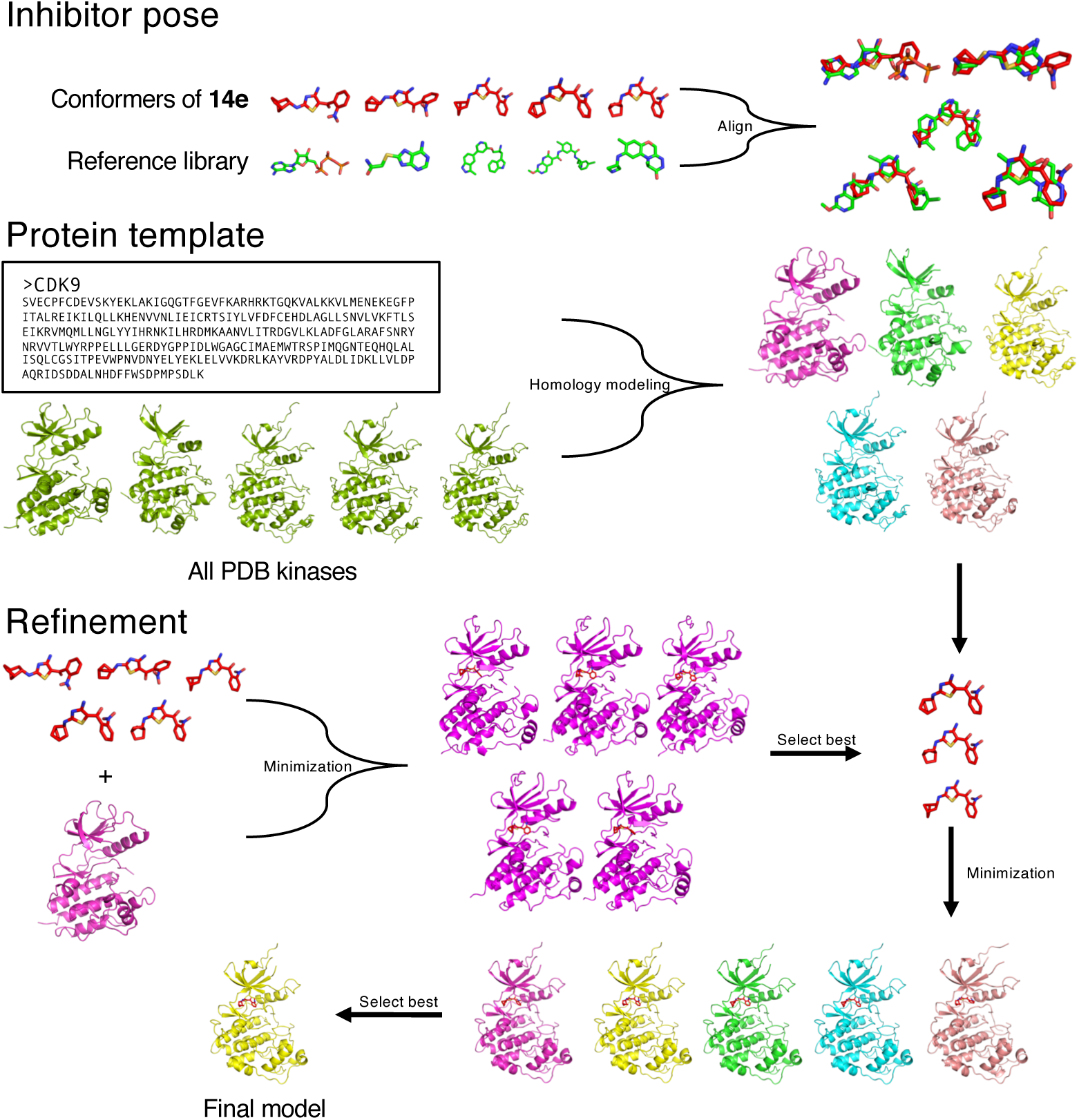
**Pipeline for building comparative models of inhibitor/kinase complexes.** The approach makes use of a template library of active kinase structures that have been pre-aligned on the basis of the protein component. 1) From the chemical structure of the query inhibitor, low-energy conformers are generated and aligned onto the set of template inhibitors from the PDB, to yield crude predictions of the query inhibitor’s pose. 2) A protein sequence alignment coupled with ligand similarity scores is used to identify suitable templates for modeling the query protein. 3) Refinement is carried out in two stages. The top-scoring 100 crude inhibitor poses are combined with the single closest protein template and minimized, then the top-scoring 10 refined inhibitor poses are combined with each of 10 protein templates and minimized. The top-scoring prediction from this refinement (on the basis of interaction energy) is taken as the final predicted model of the complex.

By focusing the search space to interactions similar those observed in the PDB, this pipeline is considerably faster than traditional high-resolution docking while also allowing for complete flexibility of both the ligand and the protein during refinement. Because of these two advantages, this pipeline can be used to explore both SAR (by modeling multiple **14e** analogs with CDK9) and selectivity (by modeling **14e** with multiple CDK kinases). We will demonstrate our application of this pipeline for both of these tasks below, but will first we summarize our campaign that led to selective CDK9 inhibitor **14e**.

### Structure activity relationships around 14e: chemistry

Prior to deconvoluting the mechanistic target of **14e** as CDK9, we executed an SAR campaign around screening hits **1–4** and assessed their activity phenotypically in the YB5 colon cancer cell line which contains a GFP reporter gene whose expression is under epigenetic control [28]. The SAR campaign involved exploring three regions of the core scaffold (**Figure 4**), namely, 1) the aryl/alkyl amine appended to position 2 of the thiazole, 2) the 4-amino group on the thiazole, and 3) the arylmethanone moiety in the 5-position of the thiazole. Analogs were synthesized by building the core aminothiazole through a Thorpe-Ziegler cyclization [44, 45]. Four general methodologies (Methods A–D) were initially employed for compounds **8a-m** and **12a-an**, obtained as reported in **Schemes 1-3**. Further exploration led to compounds **13a-l**, **14a-p**, and **17a-g**, obtained as reported in **Schemes 4-8**.

**Figure 4:**
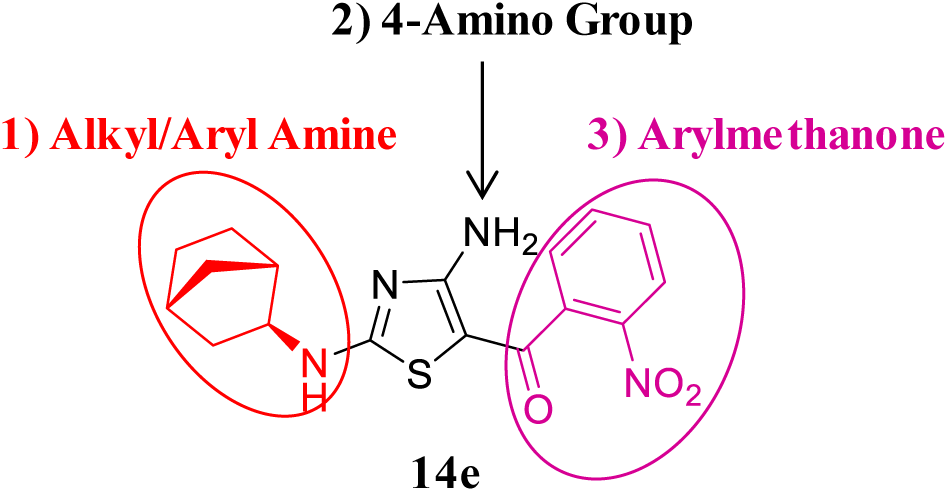
Regions of **14e** scaffold explored in the SAR campaign.

To expand the SAR around hits **2** and **4**, compounds **8a-m** were prepared using Method A (**Scheme 1**), which involved treating the appropriate isothiacyanate (**5a-m**) with cyanamide (**6**) in the presence of potassium t-butoxide, followed by reaction with (3,4-dimethoxyphenyl)(2-(methylthio)thiazol-5-yl)methanone (**7**) and subsequent cyclization to the aminothiazole (**Scheme 1**). All of the isothiocyanates used in the synthesis of **8a-m** were obtained from commercial suppliers with the exception of 8-isothiocyanato-1,4-dioxaspiro[4,5]decane (**5m**) which was synthesized using a published procedure [46]. Yields ranged from 6% to 41% over the two steps of the one-pot reaction sequence.

**Scheme 1.**
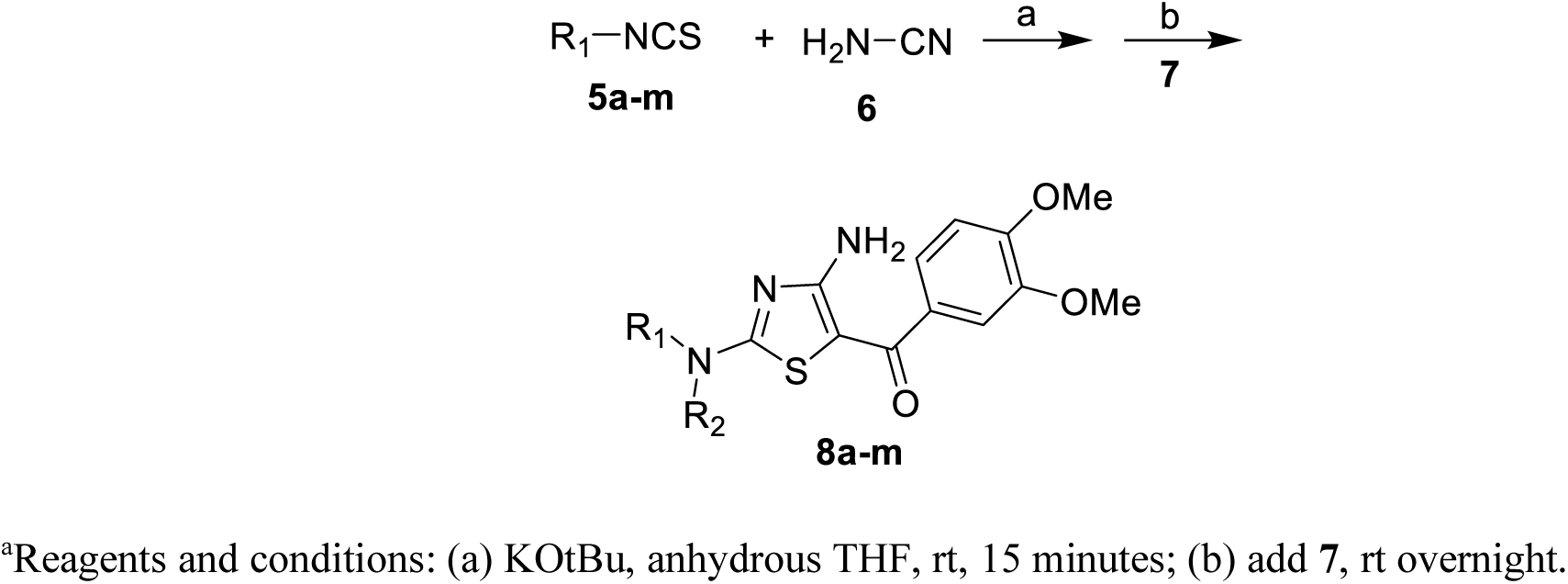
Synthesis of Compounds 8a-m. Method A.^a^

To obtain a better understanding of the SAR around hit **1**, we prepared benzodioxan analogs **12a- an**. Three synthetic methods were employed. Compounds **12a-f, h-w, z** were synthesized using Method B (Scheme 2), which differed from Method A only in the use of 2-bromo-1-(2,3-dihydro-1,4-benzodioxin- 6-yl)ethan-1-one (**10**) in place of (3,4-dimethoxyphenyl)(2-(methylthio)thiazol-5-yl)methanone (**7**). Yields ranged from 5% to 49% over the two steps and final compounds were isolated as yellow solids. For derivatives **12e, g, aa, ac-an**, the cyclization reaction was carried out in ethanol at 100°C overnight in a sealed pressure vessel in the absence of any base (Method C, Scheme 2). Yields were in the range of 10% to 72%. Method D was used to synthesize the tertiary amine-derived compounds **12x-y, ab** (Scheme 3). The appropriate cyclic secondary amine was treated with dimethyl N-cyanodithioimidocarbonate in anhydrous DMF at 70°C followed by sodium sulfide at the same temperature [47]. The one pot reaction was then cooled to 50°C and intermediate **10** was added, followed by potassium carbonate. Yields ranged from 32% to 63%.

**Scheme 2.**
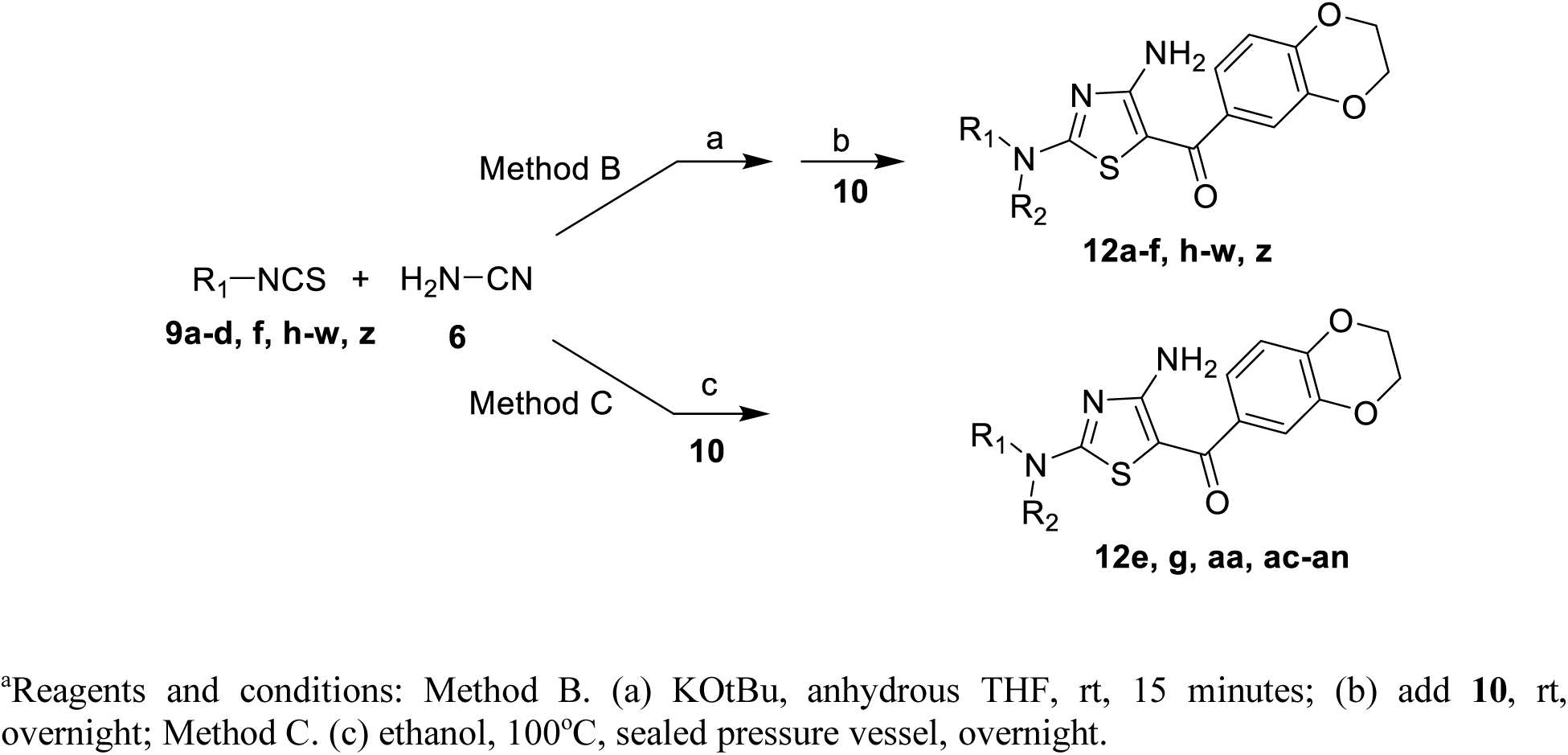
Synthesis of compounds 12a-w, z, aa, ac-an. Methods B and C.^a^

**Scheme 3.**
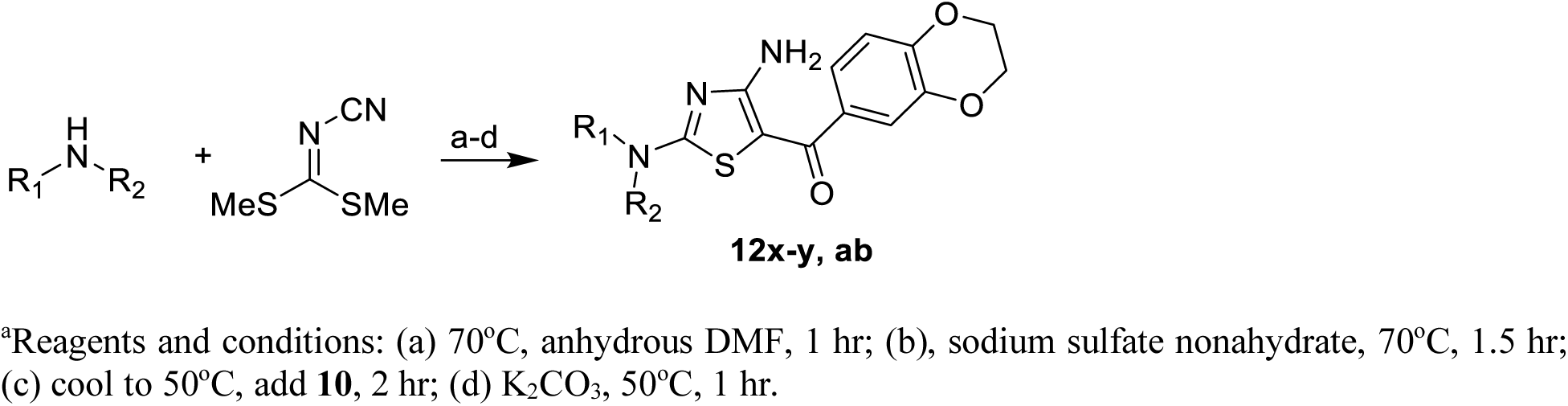
Synthesis of Compounds 12x-y, ab. Method D.^a^

To explore the effect of altering the arylmethanone moiety we synthesized thiomethyl-aminothiazole intermediates **13a-l** (Scheme 4). Compounds **13a-h** and **13j** were prepared in one step from commercially available bromoethanones in a method analogous to that for the preparation of **10**. The required bromoethanone starting materials for the synthesis of **13i, k-l** were obtained in 2-5 steps and then converted to the desired thiomethyl-aminothiazoles in one step using a procedure analogous to that for the preparation of **10**. Intermediates **13a-l** were then converted to final target compounds **14a-g** and **14i-p** using Method C with yields of the final step ranging from 14% to 99%. The piperazine-carboxamide **14h** required a different synthetic route. It was obtained in 5 steps and 6% overall yield from intermediate **13d** (Scheme 5).

**Scheme 4.**
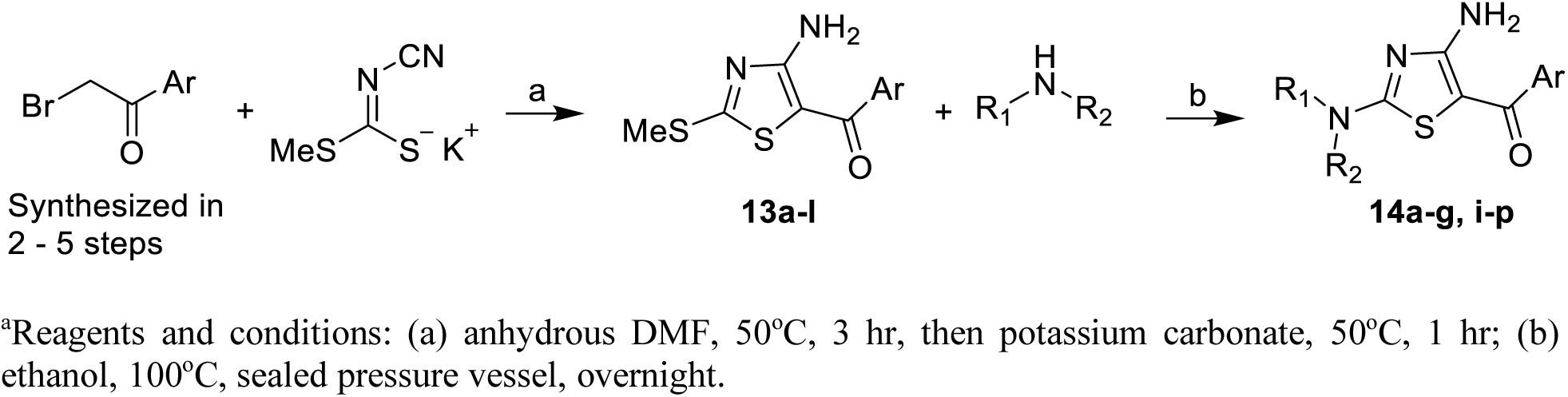
Synthesis of Intermediates 13a-l and Compounds 14a-g, i-p.^a^

**Scheme 5.**
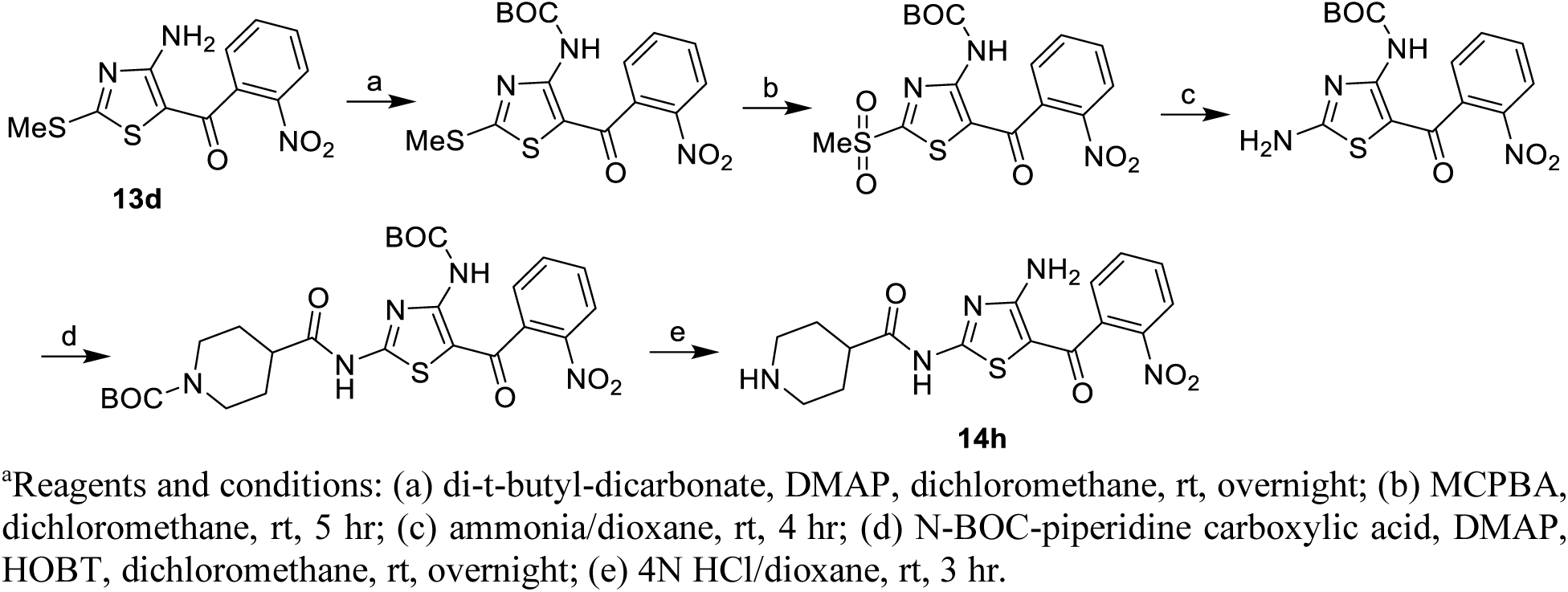
Synthesis of Compound 14h.^a^

To test the necessity of the 3-amino group on the thiazole ring we prepared acylated analogs and analogs lacking the amine functionality. Acylated derivatives **17a-d** were synthesized from **10** through a 3-step procedure (Scheme 6) that involved acylation with the appropriate acylating agent, oxidation of the thiomethyl group with 3-chloroperbenzoic acid to generate a more reactive sulfonylmethyl leaving group, and displacement of the sulfonylmethyl moiety with the required amine. Analogs **17e** and **17f**, which lacked the amine group on the thiazole ring, were synthesized in two steps (Scheme 7). 2-Thiomethylthiazole was reacted with the appropriate benzonitrile in the presence of n-butyl lithium to give an imine [48], which was hydrolyzed with aqueous acetic acid to provide the required thiazolylmethanones. These intermediates were then treated with (exo)-2-aminonorbornane to give the desired final products.

**Scheme 6.**
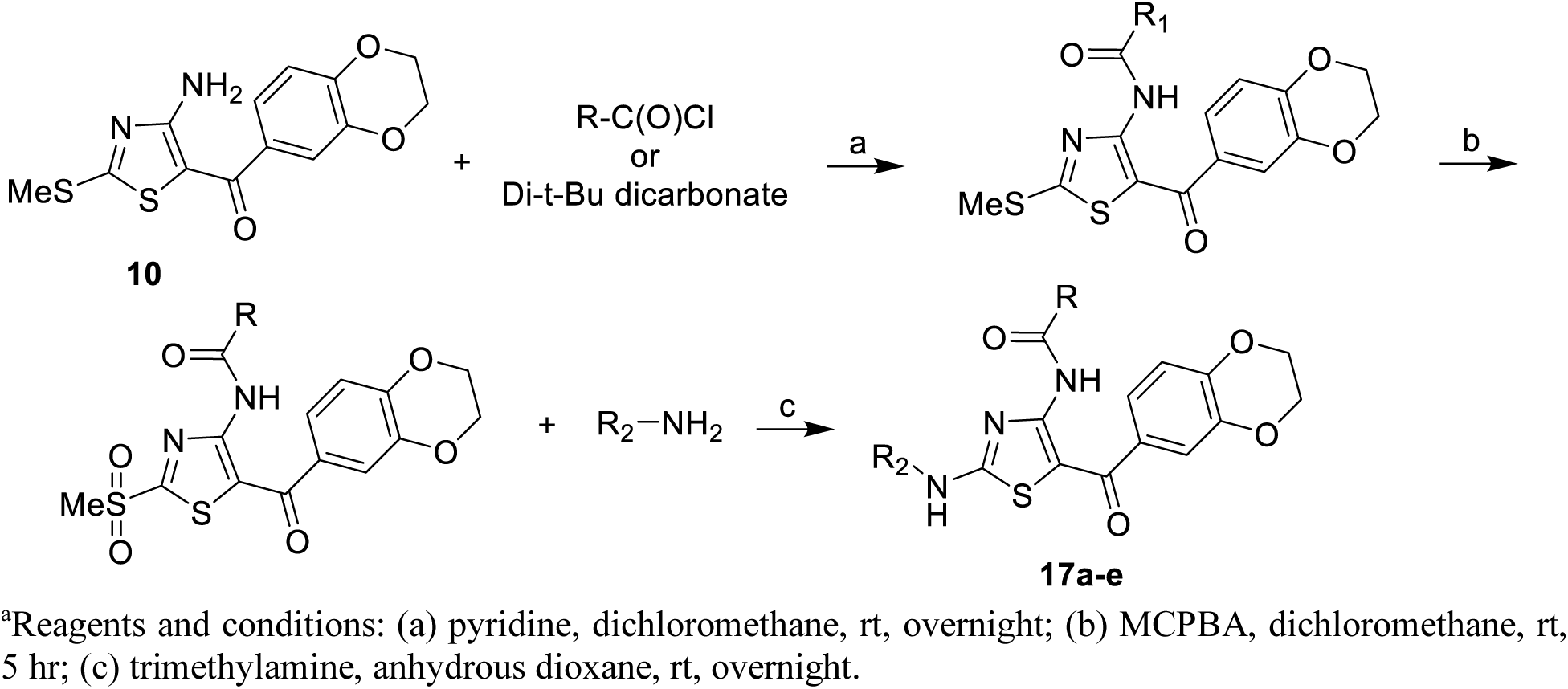
Synthesis of Compounds 17a-e.^a^

**Scheme 7.**
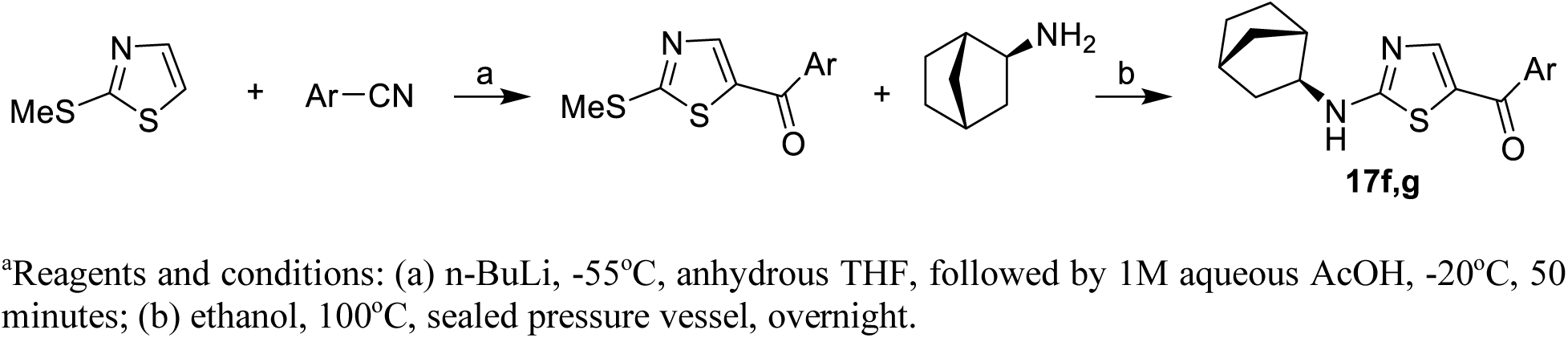
Synthesis of Compounds 17f and 17g.^a^

Compound **17e** combined a biotin group with an active analog (**12h**). Our intention was to use this as a reagent for building an affinity capture assay [49] at a time in the project when the mechanistic target of the series was not known. It was prepared in 16% yield by treating compound **12h** directly with the acid chloride of biotin, which was generated by treating biotin with thionyl chloride (Scheme 8) [50].

**Scheme 8.**
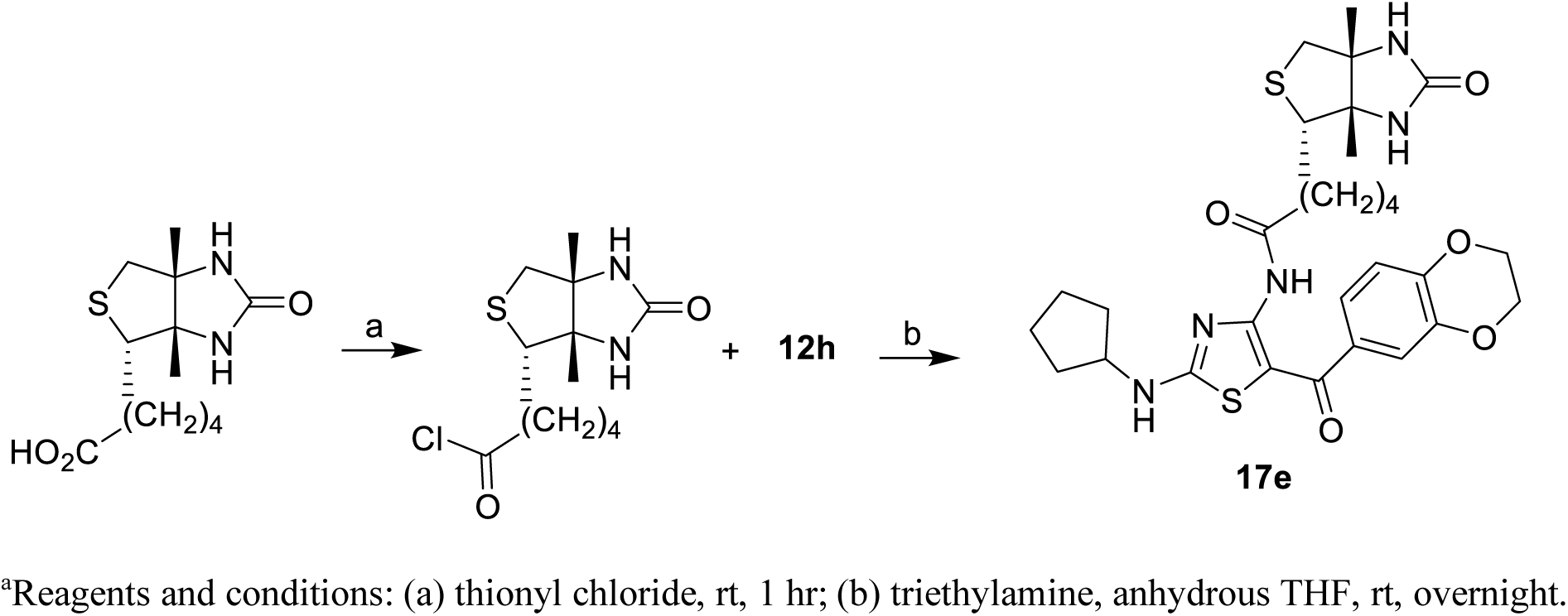
Synthesis of Compound 17e.^a^

### Structure activity relationships around 14e: cellular activity

As noted earlier, **14e** was identified and optimized through the use of a cellular assay probing for reversal of epigenetic silencing of a GFP reporter gene. Given that the molecular target of **14e** was not known at the outset of our study, as a positive control for our assay we used the histone deacetylase (HDAC) inhibitor depsipeptide (the natural product romidepsin), a marketed cancer drug [51]. Correspondingly, throughout our initial screening and subsequent optimization leading to **14e**, cellular activity was determined as the GFP fluorescence induced by the compound of interest, relative to the GFP fluorescence induced by depsipeptide (after 24-hour treatment).

Cellular activity for each compound was measured at several concentrations (ranging from 500 nM to 50 µM), and in each case compared to depsipeptide at 40 nM. Our optimization spanned 77 compounds (**Tables 1-4**), and was focused on developing compound that induced GFP fluorescence even as we reduced the compound concentration. Diversity on the thiazole core was incorporated at three positions (**Figure 4**).

**Table 1.**
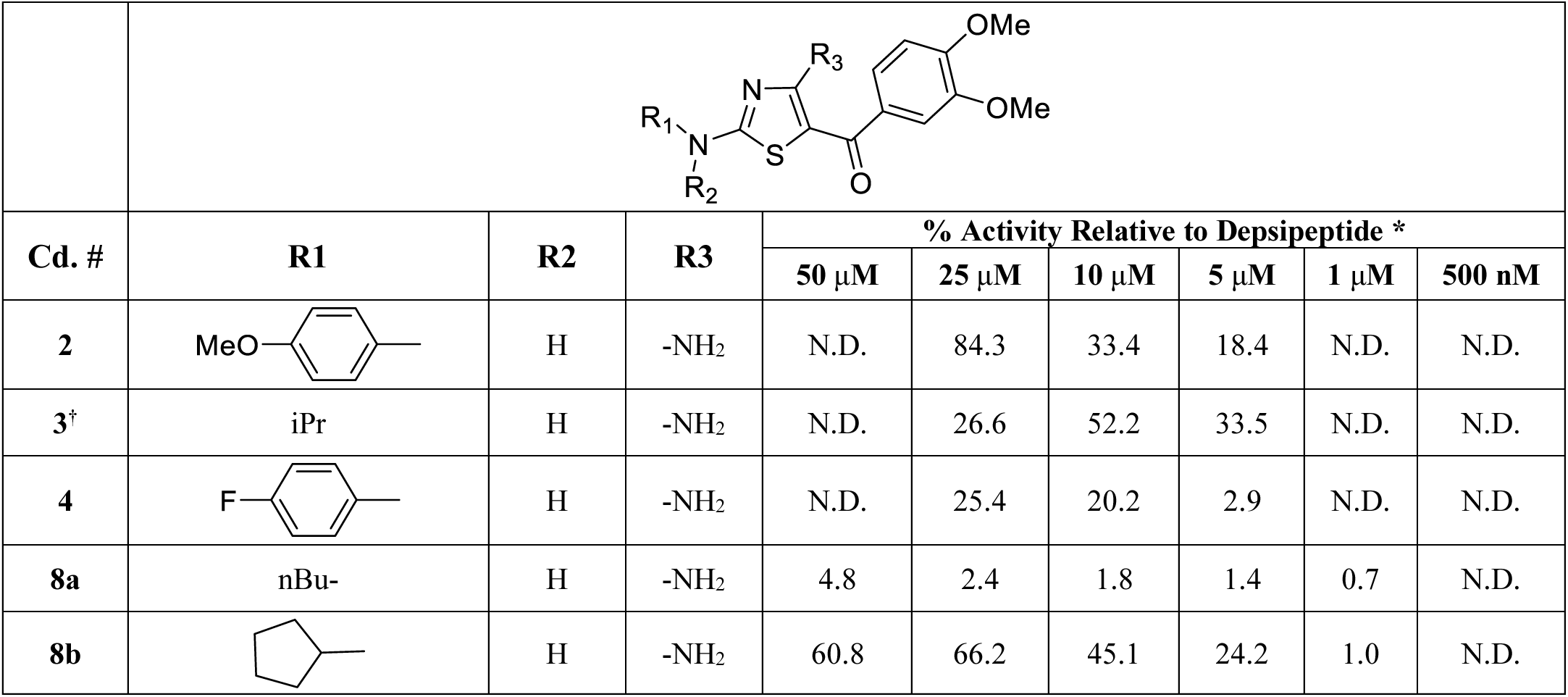
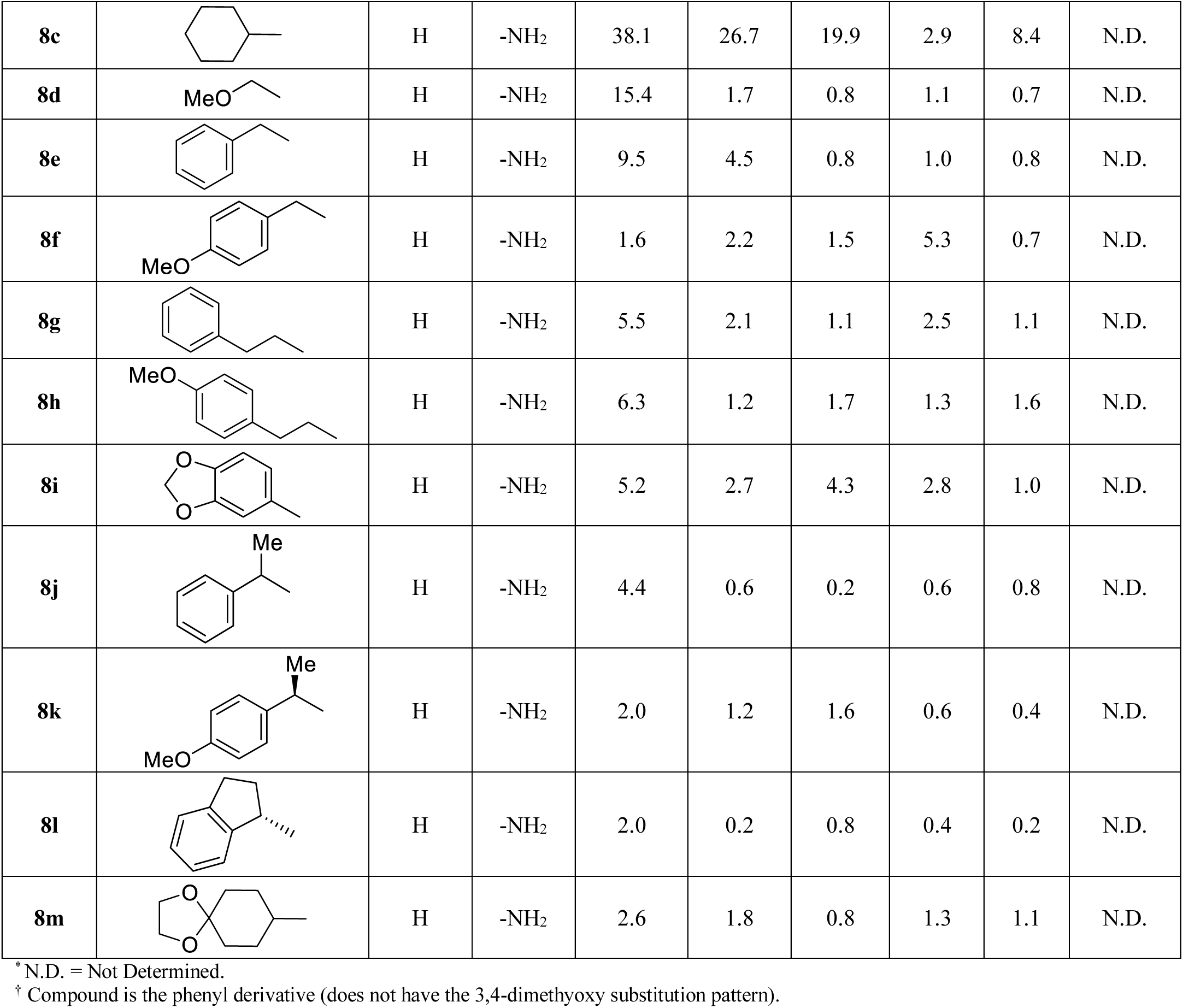
Reversal of epigenetic silencing by compounds 2-4 and 8a-m, as probed via a GFP reporter gene in YB5 cells. Activity at each compound concentration is reported relative to the activity of depsipeptide at 40 nM.

| Cd. # | R1 | R2 | R3 |  |  |  |  |  |  |
| --- | --- | --- | --- | --- | --- | --- | --- | --- | --- |
|  |  |  |  | % Activity Relative to Depsipeptide * |  |  |  |  |  |
| | | | | 50 μM | 25 μM | 10 μM | 5 μM | 1 μM | 500 nM |
| **2** |  | H | -NH<sub>2</sub> | N.D. | 84.3 | 33.4 | 18.4 | N.D. | N.D. |
| **3<sup>†</sup>** | iPr | H | -NH<sub>2</sub> | N.D. | 26.6 | 52.2 | 33.5 | N.D. | N.D. |
| **4** |  | H | -NH<sub>2</sub> | N.D. | 25.4 | 20.2 | 2.9 | N.D. | N.D. |
| **8a** | nBu- | H | -NH<sub>2</sub> | 4.8 | 2.4 | 1.8 | 1.4 | 0.7 | N.D. |
| **8b** |  | H | -NH<sub>2</sub> | 60.8 | 66.2 | 45.1 | 24.2 | 1.0 | N.D. |
| **8c** |  | H | -NH<sub>2</sub> | 38.1 | 26.7 | 19.9 | 2.9 | 8.4 | N.D. |
| **8d** |  | H | -NH<sub>2</sub> | 15.4 | 1.7 | 0.8 | 1.1 | 0.7 | N.D. |
| **8e** |  | H | -NH<sub>2</sub> | 9.5 | 4.5 | 0.8 | 1.0 | 0.8 | N.D. |
| **8f** |  | H | -NH<sub>2</sub> | 1.6 | 2.2 | 1.5 | 5.3 | 0.7 | N.D. |
| **8g** |  | H | -NH<sub>2</sub> | 5.5 | 2.1 | 1.1 | 2.5 | 1.1 | N.D. |
| **8h** |  | H | -NH<sub>2</sub> | 6.3 | 1.2 | 1.7 | 1.3 | 1.6 | N.D. |
| **8i** |  | H | -NH<sub>2</sub> | 5.2 | 2.7 | 4.3 | 2.8 | 1.0 | N.D. |
| **8j** |  | H | -NH<sub>2</sub> | 4.4 | 0.6 | 0.2 | 0.6 | 0.8 | N.D. |
| **8k** |  | H | -NH<sub>2</sub> | 2.0 | 1.2 | 1.6 | 0.6 | 0.4 | N.D. |
| **8l** |  | H | -NH<sub>2</sub> | 2.0 | 0.2 | 0.8 | 0.4 | 0.2 | N.D. |
| **8m** |  | H | -NH<sub>2</sub> | 2.6 | 1.8 | 0.8 | 1.3 | 1.1 | N.D. |

\* N.D. = Not Determined.
† Compound is the phenyl derivative (does not have the 3,4-dimethoxy substitution pattern).

We first explored the 2-amino substituent in the 3,4-dimethoxyphenylmethanone series (**8a-m**, **Table 1**). Initial observations from the screening hits included the fact that the 4-methoxyphenylamine analog **2** was more potent and effective than either the 4-fluorophenyl derivative **4** or the isopropyl compound **3**. Increasing the length of the alkyl group from isopropyl (**3**) to n-butyl (**8a**) resulted in a decrease in potency, with **8a** demonstrating less activity than either **3** or depsipeptide across the range of concentrations tested. Positioning a methoxy group on the end of an alkyl chain in the position that would roughly correspond to the methoxy moiety in the para position of **2** gave an analog (**8d**) that showed only 15% of the depsipeptide effect at the highest concentration tested (50 µM). Replacing the isopropyl group of **3** with a cyclopentyl group (**8b**) resulted in a derivative that was essentially equipotent with **3**. The analogous cyclohexyl derivative (**8c**) was not as potent as **8b** and adding oxygen functionality to the 4-position of the cyclohexyl ring (**8m**) also reduced potency compared to **3** and **8b**. Within the structural changes examined, the 4-methoxyphenyl group remained the most potent analog (**2**). Removal of the methoxy group (**8e, 8g**) decreased activity, as did lengthening the distance between the amine and the methoxy group via methylene insertion (**8f, 8h**). Addition of branching (**8j, 8k**) did not restore activity to the benzyl group, nor did conformational restraint (**8l**). Addition of a 3-oxygen functionality to give a methylenedioxy (**8i**) also decreased potency dramatically.

Because screening hit **1** was more potent than **2-4**, more effort was devoted to examining structural variation within the dihydrobenzodioxanylmethanone series (**12a-an**, **Table 2**). Many of the structural changes that reduced activity in the 3,4-dimethoxyphenylmethanone showed similar results in the dihydrobenzodioxanylmethanone scaffold. Interestingly, the isopropyl group, which provided some activity when combined with a phenylmethanone moiety (**3**) was not as well tolerated in the dihydrobenzodioxanylmethanone scaffold (**12e**). Most alkyl analogs (**12a-f, g**) demonstrated greatly reduced potency, although the isobutyl analog (**12g**) did show some activity at 50 µM. As in the 3,4-dimethoxyphenylmethanone series, the cyclopentyl group (**12h**) gave an analog that was equipotent with depsipeptide at higher concentrations. The analogous cyclohexyl (**12i**) and cycloheptyl (**12ag**) derivatives were less potent. Likewise, significantly less potent analogs resulted from incorporating the 5-amino functionality into a cyclic tertiary amine (**12x, 12y**). Placing the oxygen atom in a fixed position relative to the amine provided some analogs that showed some activity (**12ab, 12ac**), with the 3-methoxycyclopentyl derivative being essentially equipotent with the best 3,4-diemthoxyphenylmethanones **3** and **8b**. However, attempts to enhance hydrogen bonding through the introduction of other polar groups within this framework proved unsuccessful (**12aa, 12ae, 12af**).

**Table 2.**
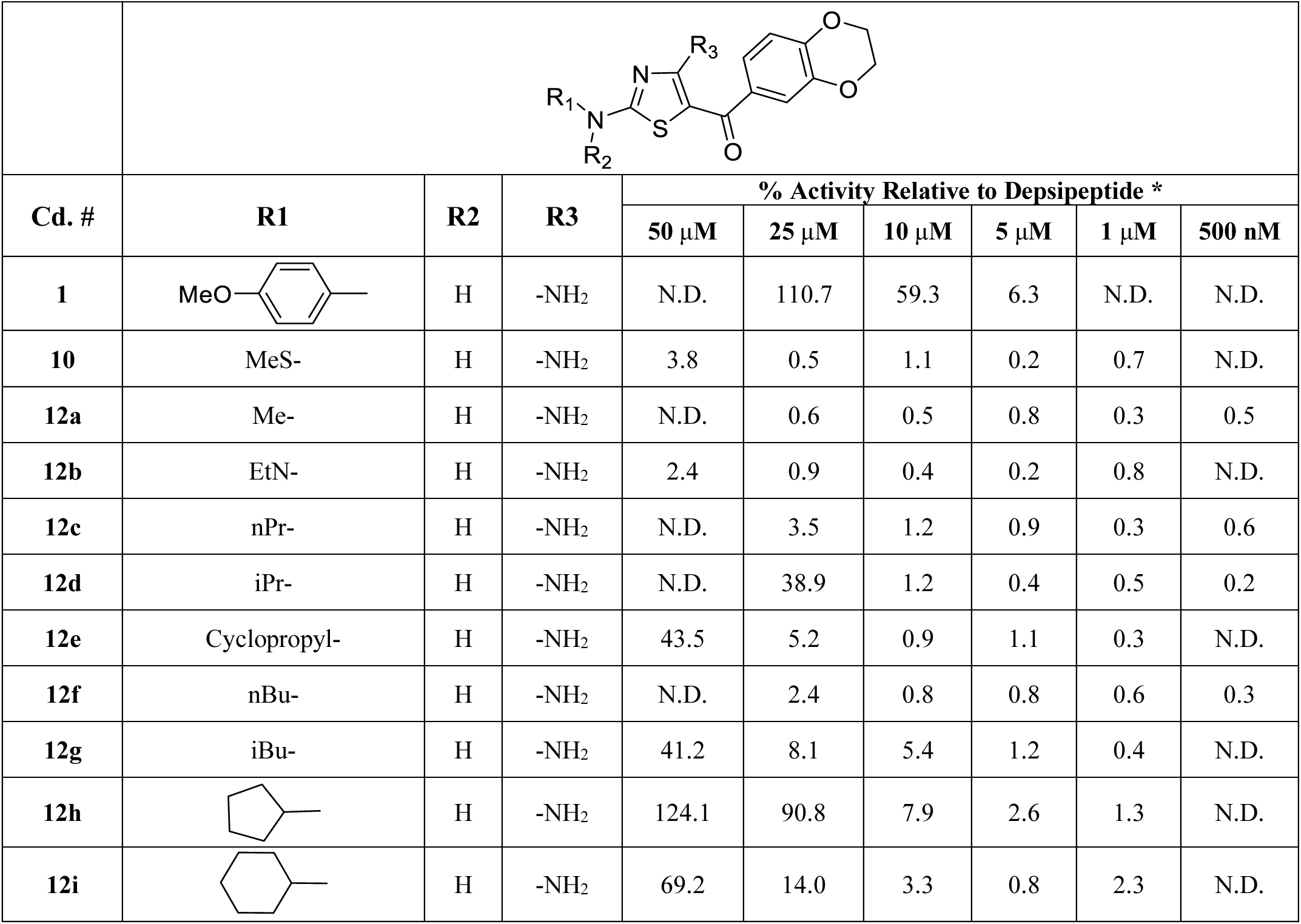
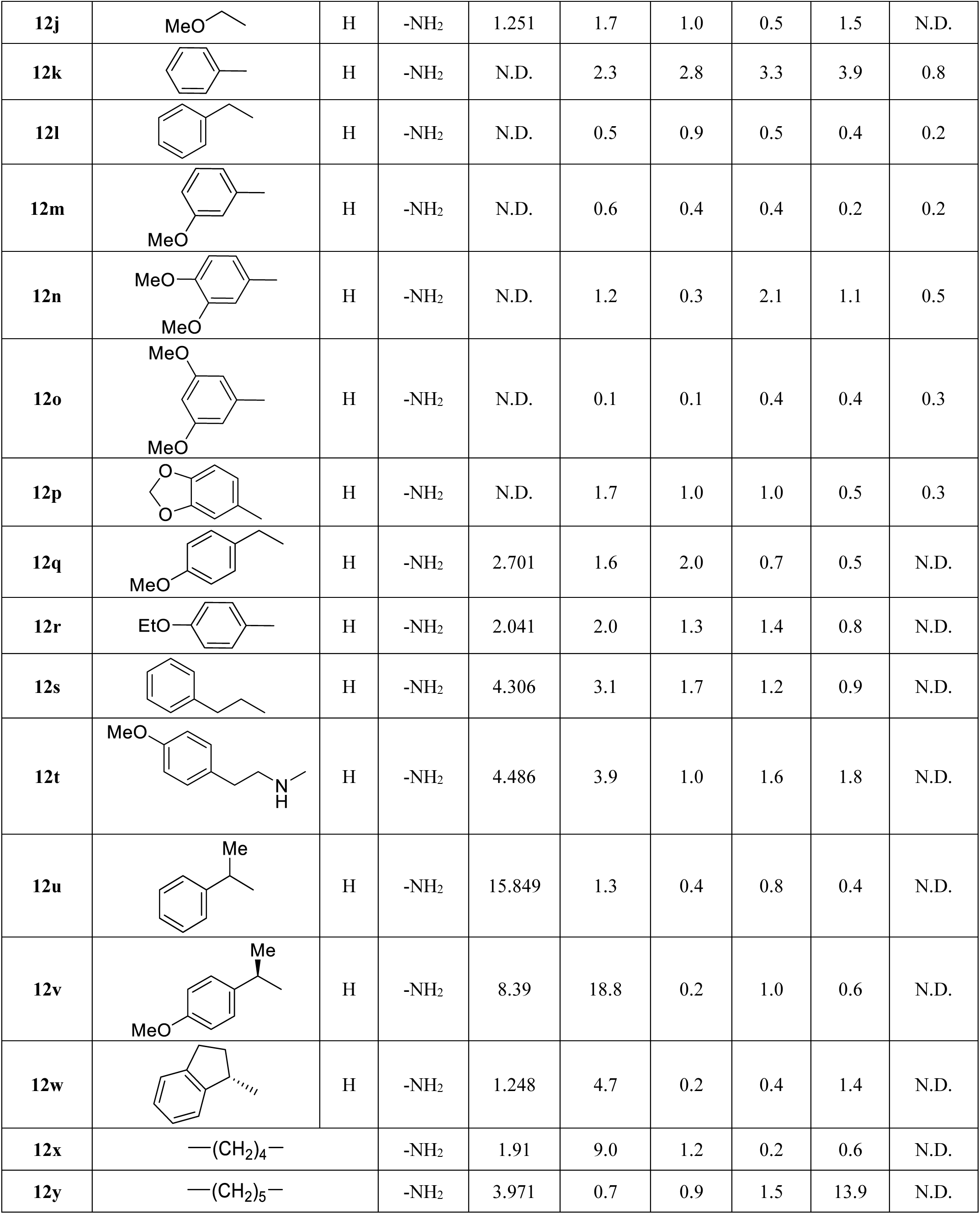
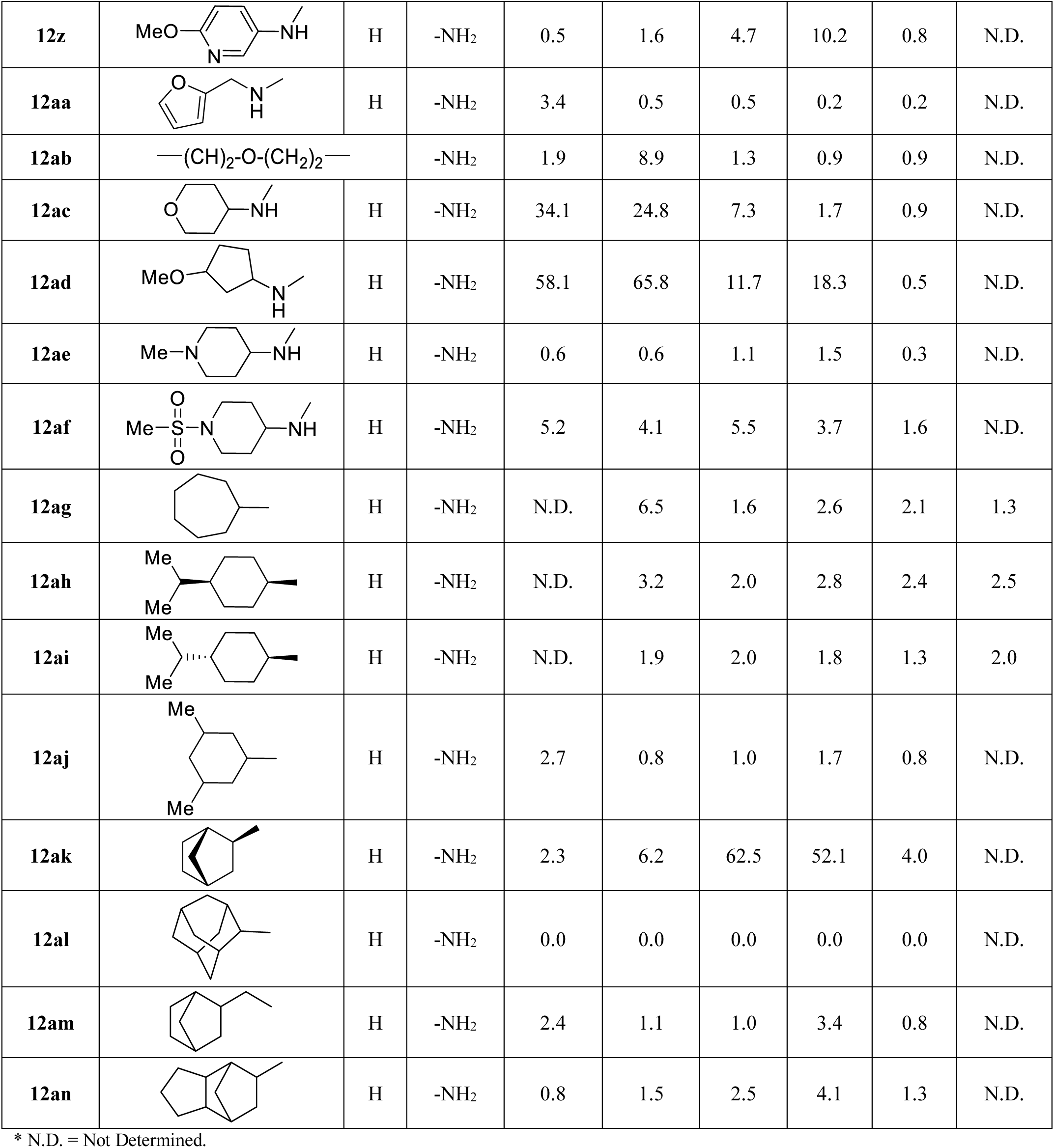
Reversal of epigenetic silencing by compounds 1, 10 and 12a-an, as probed via a GFP reporter gene in YB5 cells. Activity at each compound concentration is reported relative to the activity of depsipeptide at 40 nM.

| Cd. # | R1 | R2 | R3 |  |  |  |  |  |  |
| --- | --- | --- | --- | --- | --- | --- | --- | --- | --- |
|  |  |  |  | % Activity Relative to Depsipeptide * |  |  |  |  |  |
|  |  |  |  | 50 μM | 25 μM | 10 μM | 5 μM | 1 μM | 500 nM |
| **1** |  | H | -NH<sub>2</sub> | N.D. | 110.7 | 59.3 | 6.3 | N.D. | N.D. |
| **10** | MeS- | H | -NH<sub>2</sub> | 3.8 | 0.5 | 1.1 | 0.2 | 0.7 | N.D. |
| **12a** | Me- | H | -NH<sub>2</sub> | N.D. | 0.6 | 0.5 | 0.8 | 0.3 | 0.5 |
| **12b** | EtN- | H | -NH<sub>2</sub> | 2.4 | 0.9 | 0.4 | 0.2 | 0.8 | N.D. |
| **12c** | nPr- | H | -NH<sub>2</sub> | N.D. | 3.5 | 1.2 | 0.9 | 0.3 | 0.6 |
| **12d** | iPr- | H | -NH<sub>2</sub> | N.D. | 38.9 | 1.2 | 0.4 | 0.5 | 0.2 |
| **12e** | Cyclopropyl- | H | -NH<sub>2</sub> | 43.5 | 5.2 | 0.9 | 1.1 | 0.3 | N.D. |
| **12f** | nBu- | H | -NH<sub>2</sub> | N.D. | 2.4 | 0.8 | 0.8 | 0.6 | 0.3 |
| **12g** | iBu- | H | -NH<sub>2</sub> | 41.2 | 8.1 | 5.4 | 1.2 | 0.4 | N.D. |
| **12h** |  | H | -NH<sub>2</sub> | 124.1 | 90.8 | 7.9 | 2.6 | 1.3 | N.D. |
| **12i** |  | H | -NH<sub>2</sub> | 69.2 | 14.0 | 3.3 | 0.8 | 2.3 | N.D. |
| 12j |  | H | -NH<sub>2</sub> | 1.251 | 1.7 | 1.0 | 0.5 | 1.5 | N.D. |
| 12k |  | H | -NH<sub>2</sub> | N.D. | 2.3 | 2.8 | 3.3 | 3.9 | 0.8 |
| 12l |  | H | -NH<sub>2</sub> | N.D. | 0.5 | 0.9 | 0.5 | 0.4 | 0.2 |
| 12m |  | H | -NH<sub>2</sub> | N.D. | 0.6 | 0.4 | 0.4 | 0.2 | 0.2 |
| 12n |  | H | -NH<sub>2</sub> | N.D. | 1.2 | 0.3 | 2.1 | 1.1 | 0.5 |
| 12o |  | H | -NH<sub>2</sub> | N.D. | 0.1 | 0.1 | 0.4 | 0.4 | 0.3 |
| 12p |  | H | -NH<sub>2</sub> | N.D. | 1.7 | 1.0 | 1.0 | 0.5 | 0.3 |
| 12q |  | H | -NH<sub>2</sub> | 2.701 | 1.6 | 2.0 | 0.7 | 0.5 | N.D. |
| 12r |  | H | -NH<sub>2</sub> | 2.041 | 2.0 | 1.3 | 1.4 | 0.8 | N.D. |
| 12s |  | H | -NH<sub>2</sub> | 4.306 | 3.1 | 1.7 | 1.2 | 0.9 | N.D. |
| 12t |  | H | -NH<sub>2</sub> | 4.486 | 3.9 | 1.0 | 1.6 | 1.8 | N.D. |
| 12u |  | H | -NH<sub>2</sub> | 15.849 | 1.3 | 0.4 | 0.8 | 0.4 | N.D. |
| 12v |  | H | -NH<sub>2</sub> | 8.39 | 18.8 | 0.2 | 1.0 | 0.6 | N.D. |
| 12w |  | H | -NH<sub>2</sub> | 1.248 | 4.7 | 0.2 | 0.4 | 1.4 | N.D. |
| 12x | —(CH<sub>2</sub>)<sub>4</sub>— |  | -NH<sub>2</sub> | 1.91 | 9.0 | 1.2 | 0.2 | 0.6 | N.D. |
| 12y | —(CH<sub>2</sub>)<sub>5</sub>— |  | -NH<sub>2</sub> | 3.971 | 0.7 | 0.9 | 1.5 | 13.9 | N.D. |
| **12z** |  | H | -NH<sub>2</sub> | 0.5 | 1.6 | 4.7 | 10.2 | 0.8 | N.D. |
| **12aa** |  | H | -NH<sub>2</sub> | 3.4 | 0.5 | 0.5 | 0.2 | 0.2 | N.D. |
| **12ab** | $\text{---}(\text{CH}_2)_2\text{---O---}(\text{CH}_2)_2\text{---}$ | | -NH<sub>2</sub> | 1.9 | 8.9 | 1.3 | 0.9 | 0.9 | N.D. |
| **12ac** |  | H | -NH<sub>2</sub> | 34.1 | 24.8 | 7.3 | 1.7 | 0.9 | N.D. |
| **12ad** |  | H | -NH<sub>2</sub> | 58.1 | 65.8 | 11.7 | 18.3 | 0.5 | N.D. |
| **12ae** |  | H | -NH<sub>2</sub> | 0.6 | 0.6 | 1.1 | 1.5 | 0.3 | N.D. |
| **12af** |  | H | -NH<sub>2</sub> | 5.2 | 4.1 | 5.5 | 3.7 | 1.6 | N.D. |
| **12ag** |  | H | -NH<sub>2</sub> | N.D. | 6.5 | 1.6 | 2.6 | 2.1 | 1.3 |
| **12ah** |  | H | -NH<sub>2</sub> | N.D. | 3.2 | 2.0 | 2.8 | 2.4 | 2.5 |
| **12ai** |  | H | -NH<sub>2</sub> | N.D. | 1.9 | 2.0 | 1.8 | 1.3 | 2.0 |
| **12aj** |  | H | -NH<sub>2</sub> | 2.7 | 0.8 | 1.0 | 1.7 | 0.8 | N.D. |
| **12ak** |  | H | -NH<sub>2</sub> | 2.3 | 6.2 | 62.5 | 52.1 | 4.0 | N.D. |
| **12al** |  | H | -NH<sub>2</sub> | 0.0 | 0.0 | 0.0 | 0.0 | 0.0 | N.D. |
| **12am** |  | H | -NH<sub>2</sub> | 2.4 | 1.1 | 1.0 | 3.4 | 0.8 | N.D. |
| **12an** |  | H | -NH<sub>2</sub> | 0.8 | 1.5 | 2.5 | 4.1 | 1.3 | N.D. |

\* N.D. = Not Determined.

Removing the 4-methoxy group from the phenyl ring of **1** (**12k, 12l, 12s, 12u, 12w)**, lengthening the distance between the amine and the methoxy groups (**12q, 12t, 12v**), replacing the methoxy group with an ethoxy moiety (**12r**), and moving the methoxy group from the 4-position to the 3-position (**12m-p**) resulted in greatly reduced activity across the concentration range tested. Interestingly, the simple introduction of a heteroatom into the phenyl ring (**12z**) also reduced activity considerably.

The results suggested the presence of a lipophilic binding pocket adjacent to the 2-position on the thiazole. To explore the length and volume requirements of this putative pocket we prepared analogs **12ah-an**. The best results were obtained with the symmetrical exo-norbornyl group (**12ak**), which showed reasonable potency relative to depsipeptide at the middle of the concentration range tested (5 to 10 µM). Subtle changes in length (**12ah, 12ai, 12am, 12an**), width (**12aj**) or overall volume (**12al**) reduced activity significantly. Compound **12ak** emerged as the most potent analog with the dihydrobenzodioxanylmethanone series, showing 50% of the activity obtained with depsipeptide at the 5 µM concentration. The compound demonstrated an “inverted U” shaped concentration/response effect at higher concentrations which was not mirrored by **1**. We believe that this is the result of aqueous solubility limits (maximum aqueous solubility in 2% DMSO/phosphate buffered saline for **1** = 119 µM, compared to that of **12ak** = 18 µM; see *Experimental Section*) rather a reflection of some pharmacological effect.

We next turned to examining structural variation within the arylmethanone moiety in the 5-position of the thiazole (**Table 3**). We included groups from known series of CDK inhibitors [52] as well as undescribed (new) substitution patterns. Adding a third methoxy group to the ring in combination with either the 5-cyclopentyl amine (**14a**) or the 5-(exo-2-aminonorbornyl) (**14b**) group led to compounds that showed greatly reduced activity across the concentration range tested. Expanding (**14d**) or contracting (**14c**) the dioxan ring in combination with the 2-(exo-2-norbornylamine) gave compounds with greatly reduced activity. To explore the effect of substituting the electron rich aryl ring in the 3,4-dimethoxyphenylmethanones with an electron poor ring, we prepared 2-nitrophenylmethanone **14e**. The compound demonstrated good activity across the concentration range of 5 to 25 µM, albeit with reduced activity at 50 µM that could be explained by solubility limitations (maximum aqueous solubility = 46 µM). Moving the nitro group to the meta and para positions (**14j** and **14k**, respectively) resulted in a loss in activity compared to that seen with **14e**. Examination of a previously reported 2-nitrophenylmethanone CDK inhibitor (**14f**) [52] revealed it to be less potent than **14e**. Substitution of the 2-nitro group with 2-cyano provided **14l**, one of the most potent and effective analogs in the screening assay. Compound **14l** was found to be slightly more active at high concentrations than depsipeptide. Compound **14o**, with the oxadiazole group (a known bioisosteric replacement for nitro [53]) also demonstrated good activity. As expected, the oxadiazole bioisostere for the 2-nitro group (**14o**) was more potent than the 3-nitro bioisostere (**14p**). Unfortunately, combining the 2-nitro group with the benzodioxan ring system was not additive, as compounds **14m** and **14n** showed reduced activity. Compound **14i**, which combines the 2-nitrophenylmethanone group with a 4-aminotetrahydropyran moiety in the 2-position of the thiazole ring, proved to be one of the more potent and effective analogs, showing comparable activity to that seen with depsipeptide in the concentration range of 1 to 10 µM. This result was surprising in light of the fact that the corresponding dihydrobenzodioxanylmethanone analog (**12ac**) was significantly less potent across the concentration range tested.

**Table 3.**
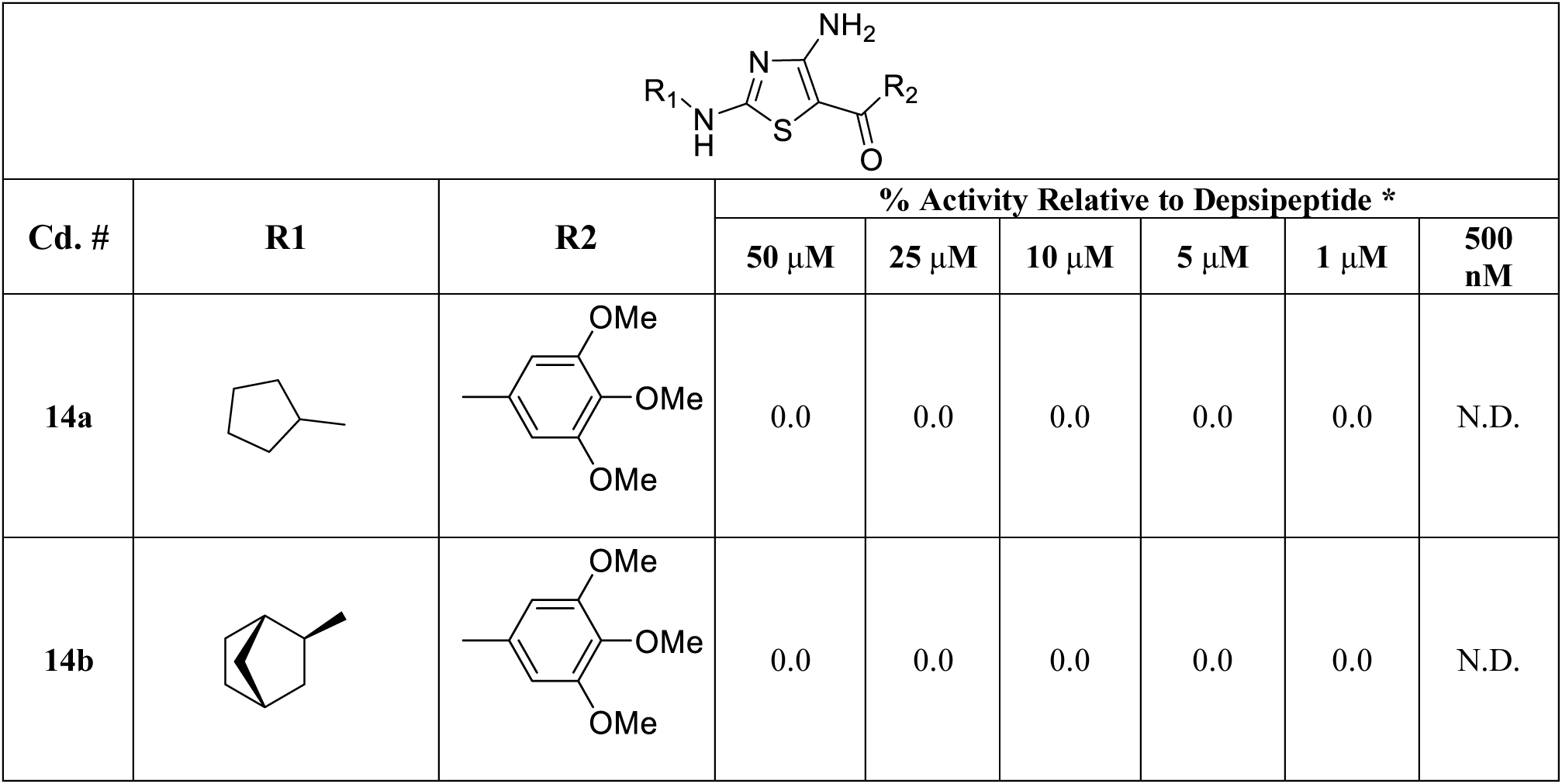
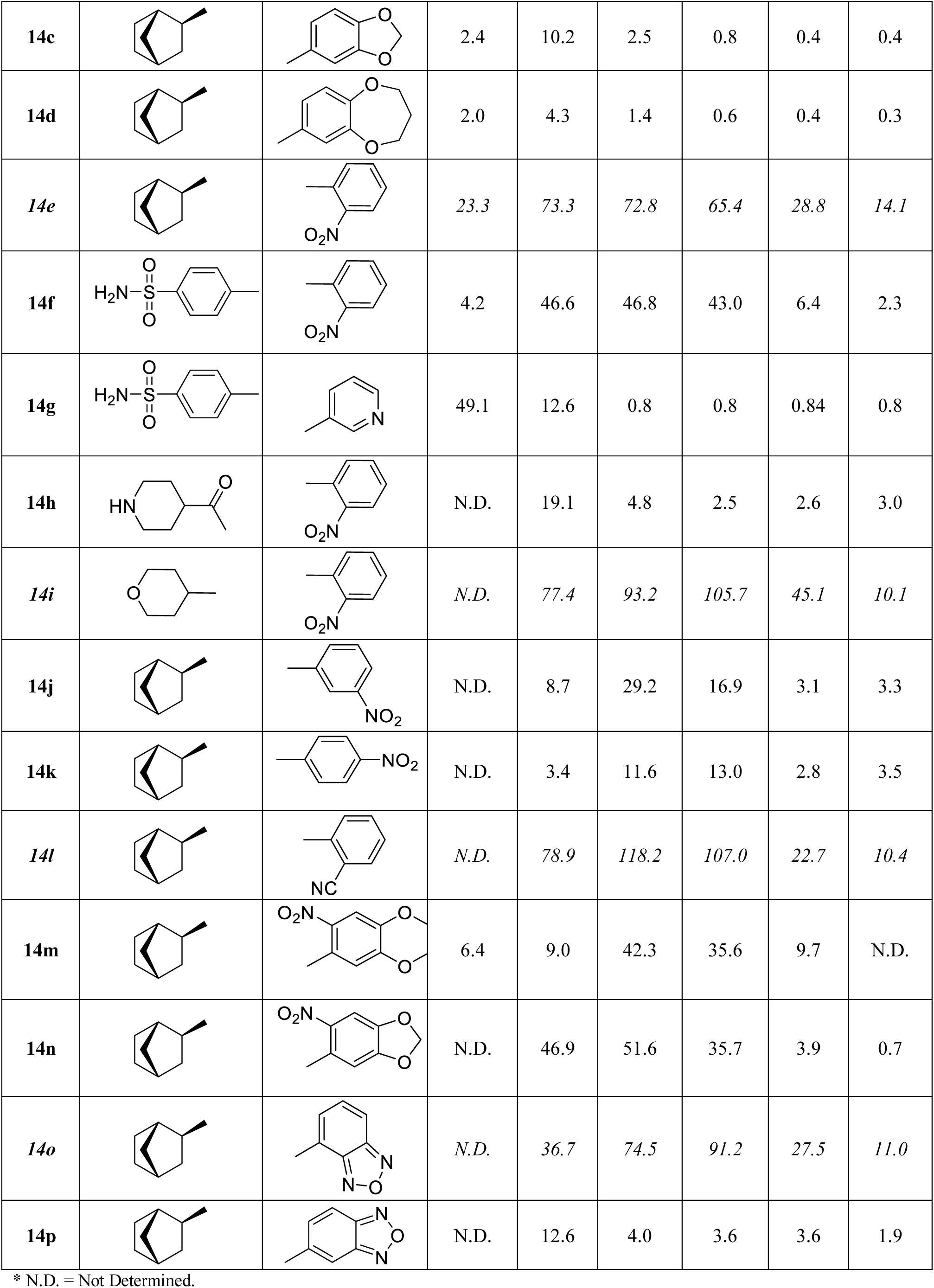
Reversal of epigenetic silencing by compounds 14a-p, as probed via a GFP reporter gene in YB5 cells. Activity at each compound concentration is reported relative to the activity of depsipeptide at 40 nM. Compounds providing >10% activity at 500 nM are *italicized*.

|  |  |  |  |  |  |  |  |  |
| --- | --- | --- | --- | --- | --- | --- | --- | --- |
| Cd. # | R1 | R2 | % Activity Relative to Depsipeptide * |  |  |  |  |  |
| | | | 50 $\mu\text{M}$ | 25 $\mu\text{M}$ | 10 $\mu\text{M}$ | 5 $\mu\text{M}$ | 1 $\mu\text{M}$ | 500 nM |
| **14a** |  |  | 0.0 | 0.0 | 0.0 | 0.0 | 0.0 | N.D. |
| **14b** |  |  | 0.0 | 0.0 | 0.0 | 0.0 | 0.0 | N.D. |
| 14c |  |  | 2.4 | 10.2 | 2.5 | 0.8 | 0.4 | 0.4 |
| 14d |  |  | 2.0 | 4.3 | 1.4 | 0.6 | 0.4 | 0.3 |
| 14e |  |  | 23.3 | 73.3 | 72.8 | 65.4 | 28.8 | 14.1 |
| 14f |  |  | 4.2 | 46.6 | 46.8 | 43.0 | 6.4 | 2.3 |
| 14g |  |  | 49.1 | 12.6 | 0.8 | 0.8 | 0.84 | 0.8 |
| 14h |  |  | N.D. | 19.1 | 4.8 | 2.5 | 2.6 | 3.0 |
| 14i |  |  | N.D. | 77.4 | 93.2 | 105.7 | 45.1 | 10.1 |
| 14j |  |  | N.D. | 8.7 | 29.2 | 16.9 | 3.1 | 3.3 |
| 14k |  |  | N.D. | 3.4 | 11.6 | 13.0 | 2.8 | 3.5 |
| 14l |  |  | N.D. | 78.9 | 118.2 | 107.0 | 22.7 | 10.4 |
| 14m |  |  | 6.4 | 9.0 | 42.3 | 35.6 | 9.7 | N.D. |
| 14n |  |  | N.D. | 46.9 | 51.6 | 35.7 | 3.9 | 0.7 |
| 14o |  |  | N.D. | 36.7 | 74.5 | 91.2 | 27.5 | 11.0 |
| **14p** |  |  | N.D. | 12.6 | 4.0 | 3.6 | 3.6 | 1.9 |

\* N.D. = Not Determined.

To examine the necessity for the 4-amino group on the thiazole ring we prepared a series of acylated analogs that incorporated the 2-cyclopentyl amine and 2-(exo-norbornylamine) groups (**17a-d**) (**Table 4**). We also prepared two derivatives containing the prototype 3,4-dimethoxyphenylmethanone (**17f**) and dihydrobenzodioxanyl (**17g**) moieties which lacked the 4-amine group. All of these compounds demonstrated low across the concentration range tested, indicating that the 3-amine was a critical component of the pharmacophore. As noted earlier, compound **17e** combined the active analog **12h** with a biotin moiety and was originally prepared as an early tool reagent for developing an affinity capture assay [49]; however, the lack of activity seen with this compound precluded its use for that purpose.

**Table 4.**
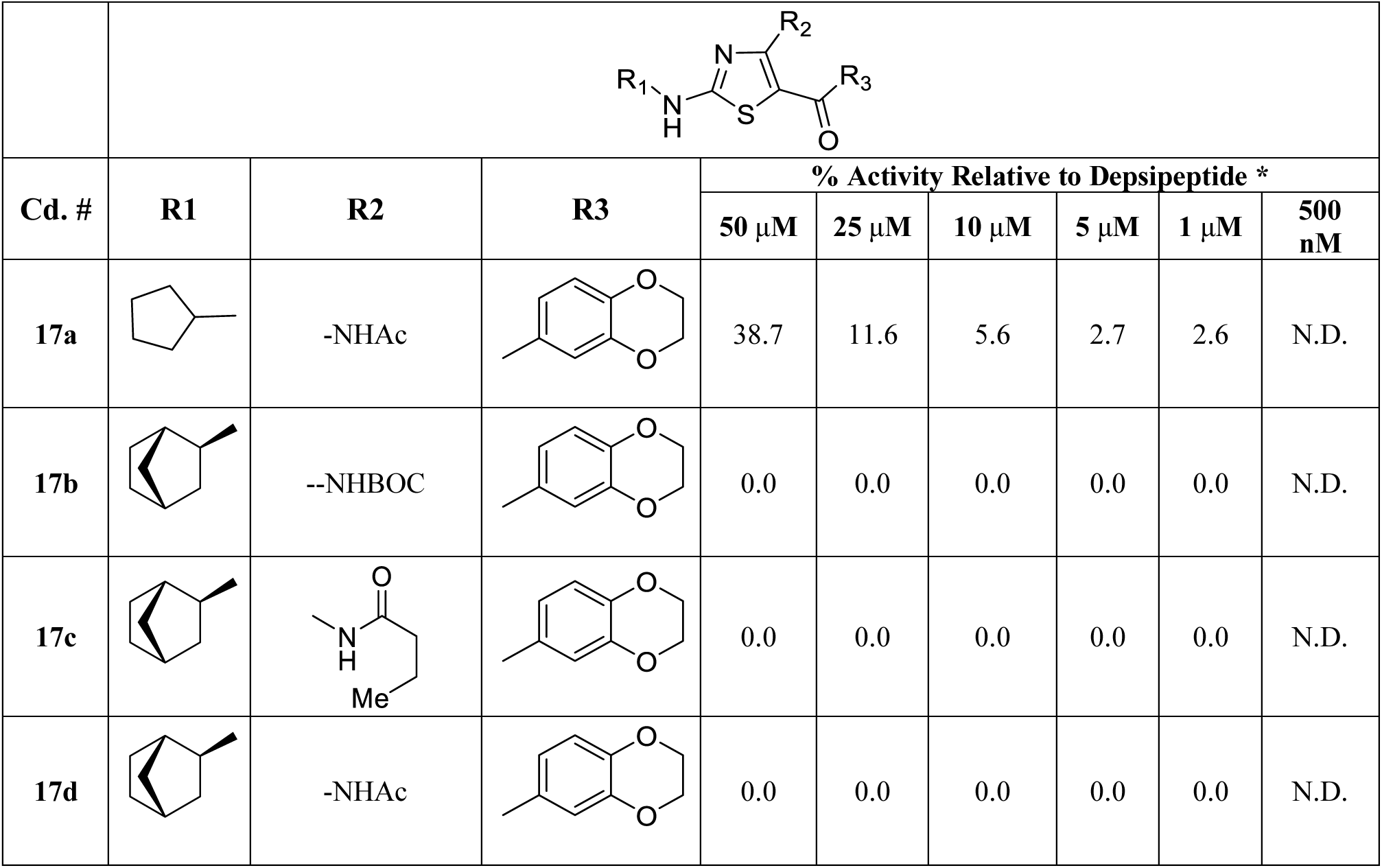
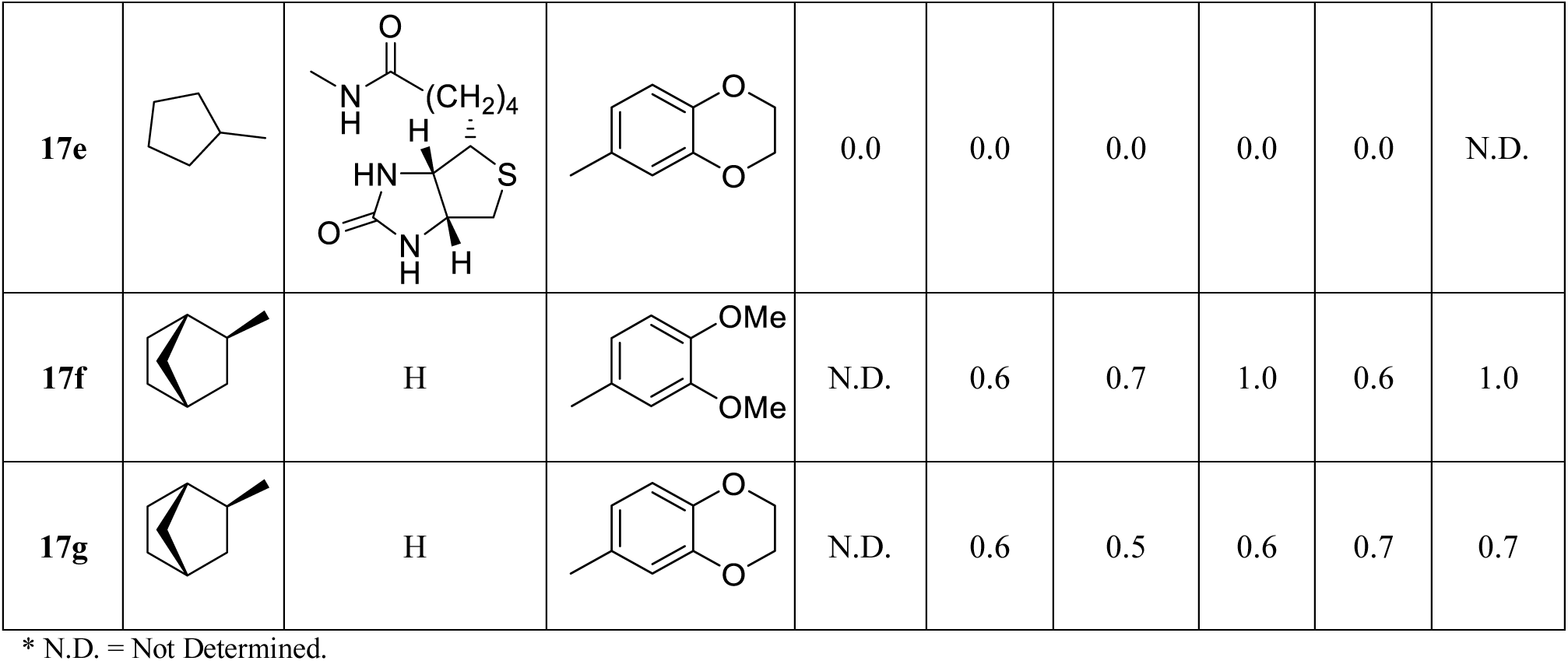
Reversal of epigenetic silencing by compounds 17a-g, as probed via a GFP reporter gene in YB5 cells. Activity at each compound concentration is reported relative to the activity of depsipeptide at 40 nM.

| Cd. # | R1 | R2 | R3 | % Activity Relative to Depsipeptide * |  |  |  |  |  |
| --- | --- | --- | --- | --- | --- | --- | --- | --- | --- |
| | | | | 50 μM | 25 μM | 10 μM | 5 μM | 1 μM | 500 nM |
| **17a** |  | -NHAc |  | 38.7 | 11.6 | 5.6 | 2.7 | 2.6 | N.D. |
| **17b** |  | --NHBOC |  | 0.0 | 0.0 | 0.0 | 0.0 | 0.0 | N.D. |
| **17c** |  |  |  | 0.0 | 0.0 | 0.0 | 0.0 | 0.0 | N.D. |
| **17d** |  | -NHAc |  | 0.0 | 0.0 | 0.0 | 0.0 | 0.0 | N.D. |
| 17e |  |  |  | 0.0 | 0.0 | 0.0 | 0.0 | 0.0 | N.D. |
| 17f |  | H |  | N.D. | 0.6 | 0.7 | 1.0 | 0.6 | 1.0 |
| 17g |  | H |  | N.D. | 0.6 | 0.5 | 0.6 | 0.7 | 0.7 |

\* N.D. = Not Determined.

### Structure activity relationships around 14e: comparative modeling

Our SAR campaign identified four compounds of interest (**Figure 5A**), each of which incorporated an amine in the 4-position of the thiazole ring, and each of which provided >10% activity at 500 nM (relative to HDAC inhibitor depsipeptide) (**Figure 5B**). At higher concentrations (up to 10 µM), each of these four derivatives ultimately reversed epigenetic silencing of the reporter gene at a level comparable to that of depsipeptide (at 40 nM). Overall, the most potent compounds included a norbornyl group at R^1^ and a benzene ring substituted in the ortho position at R^2^. The free amine in the 4-position also appeared to be important for activity.

**Figure 5:**
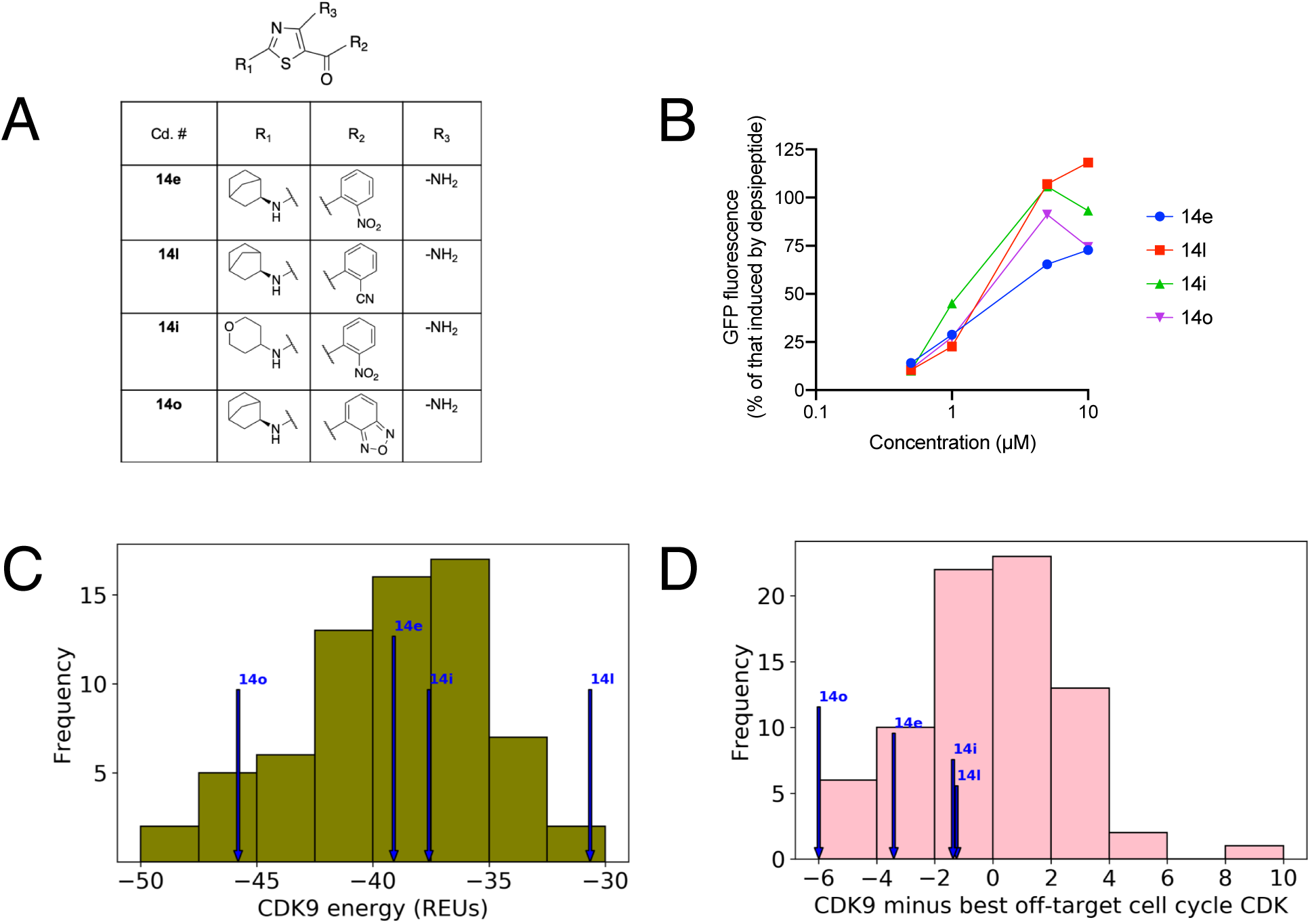
**Optimization in a phenotypic assay led to compounds that are both potent and selective for CDK9. (A)** Optimization over the course of 77 compounds yielded four compounds that reverse epigenetic silencing of a GFP reporter gene. **(B)** At high doses, each of these four compounds restore GFP fluorescence to a level comparable to that of HDAC inhibitor depsipeptide (romidepsin) at 40 nM. These four compounds in particular were prioritized because they retain activity even at lower doses. **(C)** For all 77 compounds, interaction energies were computed from comparative models of the inhibitor/CDK9 complexes. The four prioritized compounds from the phenotypic assay were not among those with the strongest interaction energies. **(D)** For each compound the difference in interaction energy was computed between the CDK9 model and the lowest-energy model from among the cell cycle CDK kinases (CDKs 1/2/4/6/7). All of the compounds prioritized from the phenotypic assay were among those that exhibited a preference for CDK9 over any of the cell cycle CDKs.

Using our comparative modeling approach, we built models of all 77 compounds in complex with their (now established) molecular target CDK9, and we evaluated the inhibitor/kinase interaction energy. Since our pipeline uses the Rosetta energy function [54], we report energies in “Rosetta energy units” (REUs). To our surprise, the compounds that most potently induced GFP fluorescence were *not* those with the best interaction energies (**Figure 5C**): indeed, based on these energies one would not have distinguished these four compounds from the rest of the set. This observation implied that either our comparative models were not yielding interaction energies reflective of these compound’s activity towards CDK9, or that our phenotypic assay was providing richer information than simply binding affinity against CDK9.

From the outset, we were quite surprised that optimization in this phenotypic assay yielded compounds that were not only potent against CDK9, but selective as well. This selectivity was most notable with respect to CDK2, given that most CDK9 inhibitors also inhibit CDK2 [29, 30]. Given that simply optimizing for potency against CDK9 would not necessarily be expected to yield selective inhibitors, we therefore hypothesized that our phenotypic screen was indirectly yielding information on selectivity to guide our optimization. As noted earlier, certain other CDKs are key regulators of the cell cycle (particularly CDKs 1/2/4/6/7): a compound that inhibited one of these CDKs in addition to CDK9 would reverse epigenetic silencing of GFP (increasing fluorescence), but would also restrict cell growth (decreasing fluorescence). In retrospect, optimal potency in this assay would best be achieved by a compound that provides the largest possible discrimination between CDK9 and the cell cycle CDKs.

To test this hypothesis, we next built models of all 77 compounds in complex with each of the cell cycle CDKs. For each compound we extracted from these models the strongest interaction energy across the set of cell cycle CDKs; we then calculated the difference in interaction energy between CDK9 and the top-scoring cell cycle CDK (**Figure 5D**). Of these 77 compounds, about half have negative values (i.e., they are predicted to bind CDK9 more tightly than any of the cell cycle CDKs) and about half have positive values (i.e., CDK9 is not the most preferred target for these compounds). Already this is a notable observation, in that our computational method revealed a preference for CDK9 in particular, over the other five competing CDK kinases for this compound set; this occurs despite the fact that the comparative modeling pipeline is in no way biased towards CDK9. Further, the four compounds that most potently induced GFP fluorescence in this reporter assay were clustered among the more negative values from the computational models.

Collectively these results begin to suggest that our comparative modeling pipeline can provide insights into the selectivity of candidate kinase inhibitors, at least amongst this collection of thiazoles.

### Selectivity of 14e, and of other CDK9 inhibitors

In course of our earlier studies, we had used kinase assays (with purified protein) to test for selectivity of **14e** against five other CDK kinases [28]. The sample of **14e** used for these biochemical studies was synthesized from a racemic building block, and accordingly contained a mixture of the two stereoisomers. We later synthetized enantiomerically-pure **14e**, and found both configurations to exhibit nearly-identical inhibition of CDK9. To facilitation comparison with these experimental data, our initial comparative modeling correspondingly included both stereoisomers of **14e**. Subsequent analysis here too revealed essentially no difference in the interaction energies of the two stereoisomers.

Enabled by our refined computational pipeline, we proceeded to model **14e** in complex with all members of the CDK family. To complement these computational results, we additionally tested **14e** for inhibition of eight more CDK kinases that were not included in our previous study (**Table 5**). Comparison of the results from kinase assays and from comparative modeling are in broad agreement with one another: in both experiments, CDK9 is the most preferred target of **14e** by a considerable margin.

**Table 5.**
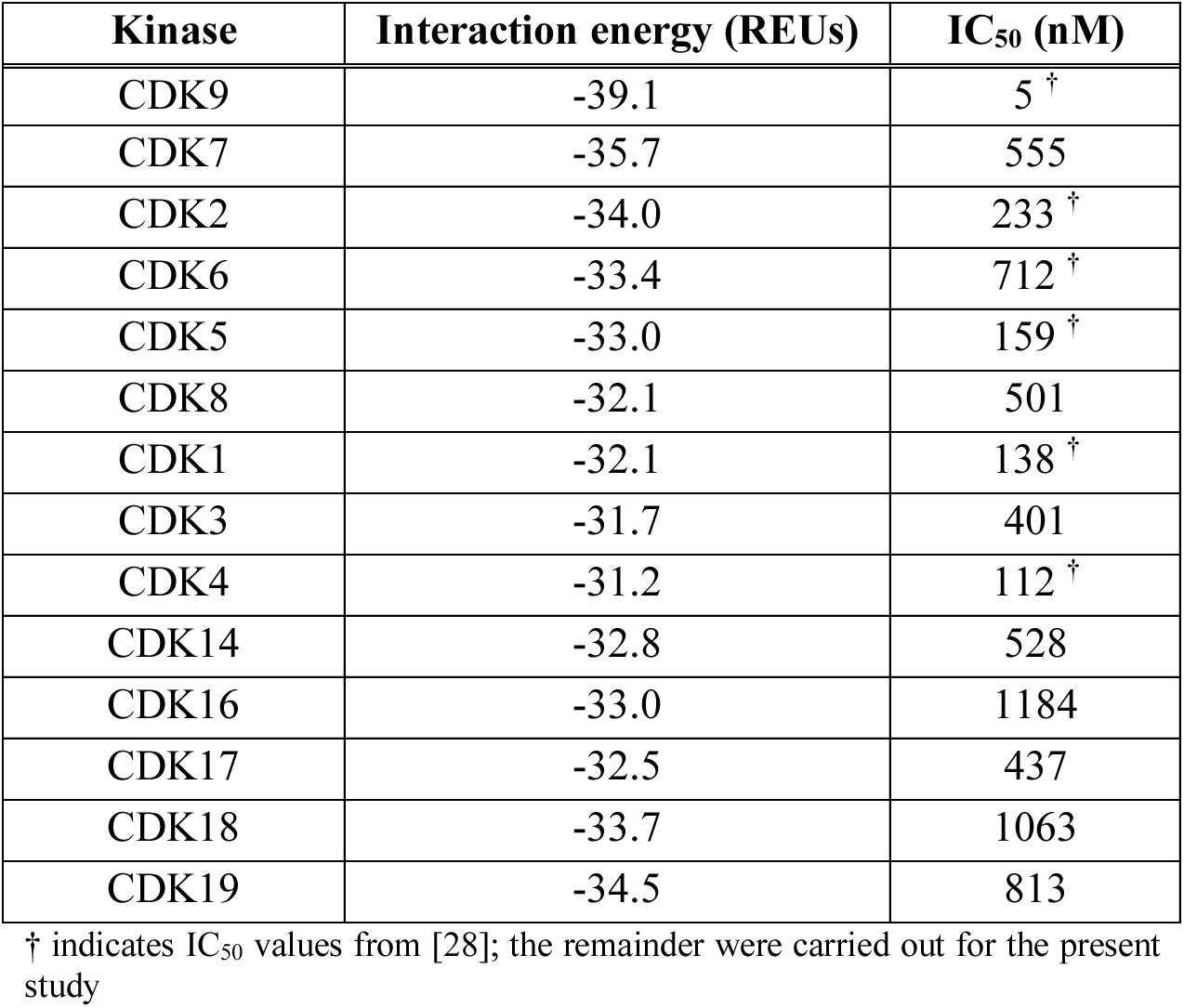
Selectivity of 14e, from comparative modeling and from biochemical kinase assays. In both experiments **14e** is found to exhibit selectivity for CDK9 over all other CDK kinases. Interaction energies from comparative models are reported in Rosetta Energy Units (REUs).

To explore the feasibility of predicting selectivity for CDK9 inhibitors beyond **14e**, we next examined a recently-compiled set of clinical candidates targeting CDK9 for acute myeloid leukemia (AML) [22]. Excluding **14e**, which is still in pre-clinical development, this set included eight compounds: six of these are currently in clinical trials (Alvocidib, AT7519, Dinaciclib, P276-00, SNS-032, TG02) and the other two are in pre-clinical development (CDKI-73, LY2857785). The chemical structures of these eight compounds (**Figure 6**) demonstrate their broad structural diversity.

**Figure 6:**
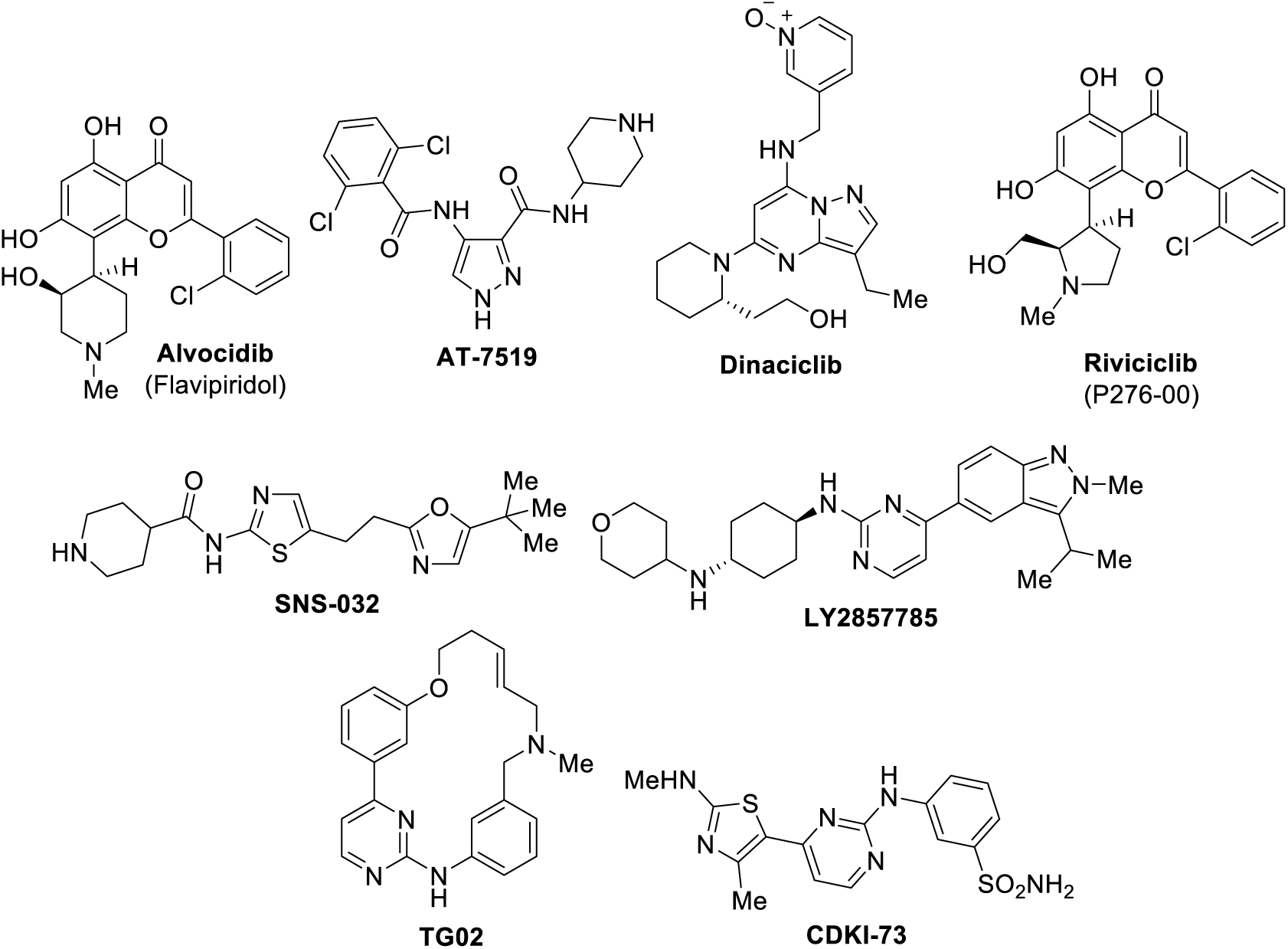
Structures of current clinical candidates targeting CDK9 for acute myeloid leukemia.

For each of these CDK9 inhibitors, activity has been evaluated against various other CDK kinases [55-59,29,60-66]. These selectivity profiles had been conveniently compiled into a recent review [22]. For each compound, we used our comparative modeling approach to build models of the compound in complex with each CDK kinase for which activity had been reported, and then extracted the interaction energy from each model. To report on selectivity, we expressed each interaction energy relative to that of the CDK9 complex, such that positive values indicate a preference for CDK9 and negative values imply that the compound is predicted to bind another CDK kinase more tightly than CDK9.

For each compound, we plotted the relative interaction energy from these comparative models as a function of the experimentally observed activities (**Figure 7**). Thus, a compound that is computationally predicted to be selective for CDK9 will have the red dot positioned far below any of the blue dots, and a compound that is experimentally observed to be selective for CDK9 will have the red dot positioned far to the left of any blue dots. For **14e**, this plot simply re-formats the results presented earlier (**Table 5**). As noted earlier, both the computational and biochemical characterization concur that **14e** has a strong preference for CDK9 over the 13 other CDK kinases tested.

**Figure 7:**
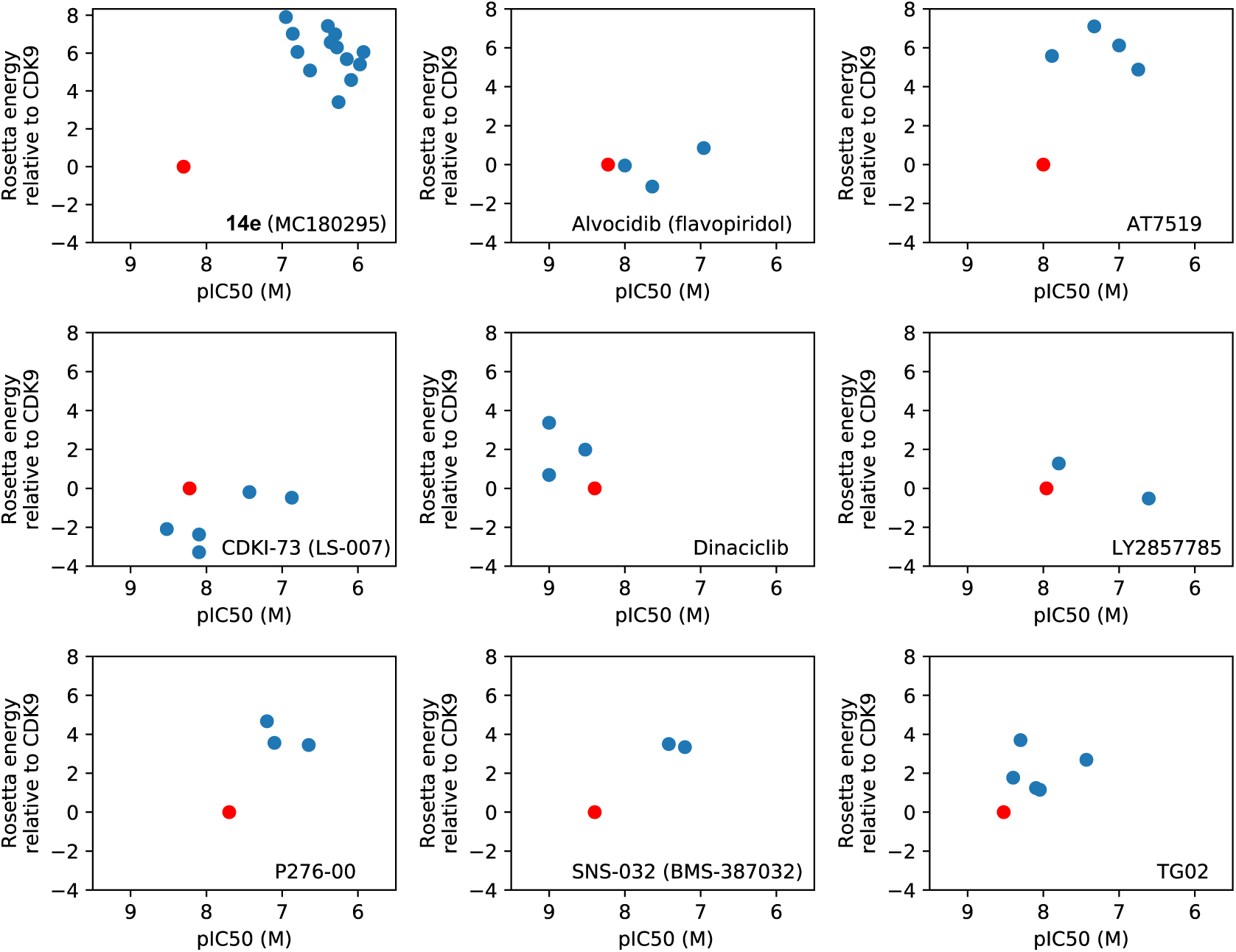
**Evaluating selectivity of other CDK9 inhibitors.** Selectivity profiles are presented for each of nine diverse CDK9 inhibitors in clinical development. Each plot corresponds to a different compound; on each plot the *red* dot shows activity against CDK9 and each *blue* dot corresponds to activity against a different CDK kinase. For each inhibitor/kinase pair, the x-axis denotes experimentally-derived inhibition (-log_10_(IC_50_)), and the y-axis gives the interaction energy from comparative modeling using the computational pipeline developed herein (energies are reported relative to the interaction energy of the corresponding inhibitor/CDK9 complex). Thus, a given compound is computationally predicted to be selective if the red dot is far below the blue dots; it is experimentally found to be selective if the red dot is far to the left of the blue dots.

While most of the other inhibitors have not been evaluated against a similarly comprehensive panel of CDK kinases, certain trends are evident. Other than **14e**, the compounds with the strongest experimentally-observed selectivity for CDK9 are P276-00 and SNS-032 (**Figure 7**, bottom row). In both cases, the computational predictions concur that these compounds are likely to exhibit selectivity for CDK9. Each of the other six compounds has been experimentally shown to have a very small preference for CDK9, or even to prefer a different CDK kinase. Encouragingly, the interaction energies from comparative modeling recapitulate the lack of selectivity for five of these six compounds. The sole failure from the comparative modeling approach is in the case of AT7519 (**Figure 7**, top right): our computational method predicts that this compound will be selective for CDK9, but in fact this compound inhibits CDK5 almost as potently as CDK9. The inappropriately predicted selectivity in this case could be because the model of the AT7519/CDK9 complex was deemed overly favorable by our energy function, or because the model of the AT7519/CDK5 complex missed key interactions, or because the model of the AT7519/CDK5 complex was not scored sufficiently favorably. While it falls beyond the scope of the present study to distinguish between these possibilities, we expect that further exploration of AT7519 selectivity can drive improvements in our computational methods moving forward.

Overall however, this experiment demonstrated that our comparative modeling pipeline can indeed distinguish whether or not a given CDK9 inhibitor is likely to prove selective, even for chemical scaffolds beyond the thiazole core used in **14e** and its analogs.

### Structure determination of 14e in complex with CDK9

While the successful recapitulation of selectivity suggests that our comparative modeling approach is able to provide an accurate representation of how MC180295 engages various kinases, direct structural evidence was still required to support this approach. To validate the modeling result, we obtained crystals of the CDK9/Cyclin-T1/**14e** complex by soaking **14e** into CDK9/Cyclin-T1 crystals, and determined the crystal structure to 3.1 Å (**Figure 8A**, **Table 6**). The CDK9 kinase in the CDK9/Cyclin-T1/**14e** complex adopts a typical kinase fold consisting of an N-terminal lobe and a C-terminal lobe linked by a hinge region.

**Figure 8:**
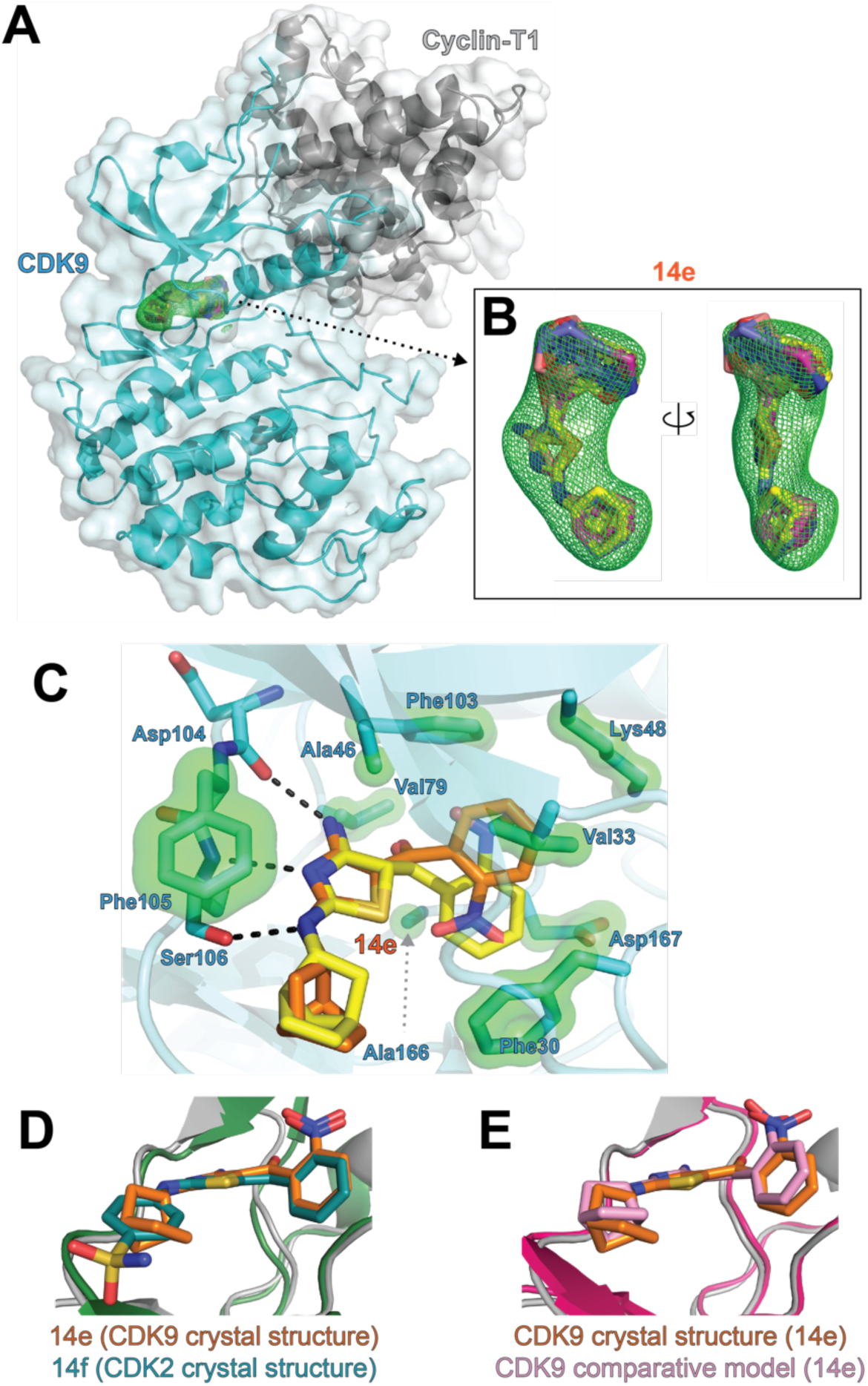
**Crystal structure of the 14e/CDK9 complex. (A)** The compound **14e** in the CDK9-cyclinT1 crystal structure. CDK9 is shown in *cyan*, cyclin-T1 is shown in *gray*, and **14e** is in *orange* covered by electron density in *green* mesh. **(B)** *Fo − Fc* omit map of **14e** contoured at 3.0σ. Four (equally-occupied) copies of **14e** were built into the density, reflecting alternate norbornane stereoisomers and a flipped nitrophenyl conformation. **(C)** Interactions between **14e** and CDK9. Side chains of the residues making hydrophobic interaction with **14e** are represented by a light green surface. Hydrogen interactions are denoted by dotted lines. Two representative models of **14e** are presented; all four make essentially identical interactions with CDK9. **(D)** One model from the **14e**/CDK9 crystal structure is superposed onto a previous crystal structure of a slightly different nitrophenyl-diaminothiazole compound (**14f**) in complex with CDK2 (PDB ID 3QXP). Both inhibitors engage their cognate kinases in a very similar manner. **(E)** This model from the **14e**/CDK9 crystal structure is superposed with our comparative model of this complex, showing very close agreement: 1.0 Å RMSD overall, and 0.83 Å RMSD for the crystallographic bound ligand (for the crystallographic pose with the same stereoisomer and ring orientation as our comparative model, and aligning on the protein only).

**Table 6.** Crystallographic data collection and refinement statistics (PDB ID 6W9E).

|  |  |
| --- | --- |
| **Data collection** | **CDK9/Cyclin-T1/14e** |
| Light Source | NSLS-II AMX |
| Wavelength (Å) | 0.97935 |
| Space group | H3 |
| Cell dimensions |  |
| *a*, *b*, *c* (Å) | 173.34, 173.34, 97.368 |
| $\alpha$, $\beta$, $\gamma$ (°) | 90, 90, 120 |
| Resolution (Å) | 28.89~3.10 (3.18~3.10) |
| *R*<sub>merge</sub> (%) | 6.1 (88.5) |
| *I*/$\sigma$ | 14.90 (1.20) |
| CC(1/2) | 99.9 (56.6) |
| Completeness (%) | 97.8 (93.4) |
| No. reflections | 87946 (6339) |
| Multiplicity | 4.4 (4.5) |
| **Refinement** |  |
| Resolution (Å) | 28.89~3.10 |
| *R*<sub>work</sub>/*R*<sub>free</sub> (%) | 17.2/22.1 |
| No. atoms |  |
| Protein | 4555 |
| Ligand/ion | 25 |
| Water | 28 |
| Wilson B factor (Å<sup>2</sup>) |  |
| Protein | 109.2 |
| Ligands | 96.5 |
| R.m.s. deviations |  |
| Bond lengths (Å) | 0.010 |
| Bond angles (°) | 2.05 |
| Ramachandran |  |
| Favored regions (%) | 90.1 |
| Allowed regions (%) | 98.2 |

*R*<sub>free</sub> was calculated using 5% of the data and the same sums
Values in parentheses are for highest-resolution shell

A fully-occupied **14e** molecule is observed in the crystal structure based on the strong electron density. That said, a mixture of the two **14e** stereoisomers was used in crystallization (due to synthesis from a racemic starting material) and the resolution of this structure does not allow us to determine whether one or both of these equipotent configurations are present. The density also does not unambiguously define the orientation of the nitrophenyl group: for this reason, we ultimately built four (equal occupancy) well-fitted models of the compound (**Figure 8B**).

The compound is accommodated in a hydrophobic pocket in the ATP-binding site, comprising residues Phe30, Val33, Ala46, Val79, Phe103, Phe105, and Ala166. Moreover, the diaminothiazole core makes canonical hydrogen bonding interactions to the main chain of the CDK9 hinge region involving Asp104, Phe105, and Ser106 (**Figure 8C**). Similar interactions are also seen in the crystal structures of aminothiazole-containing inhibitors bound to other kinases, such as dasatinib bound to Abl kinase [67]. Finally, the bulky (and hydrophobic) norbornyl group is located on the surface of the protein complex, interacting with the loop region just C-terminal of the hinge.

Interestingly, the 1.75 Å resolution crystal structure of a similar nitrophenyl-substituted diaminothiazole (**14f**) in complex with CDK2 has been solved [52]. The overall architecture of the kinase in this structure is nearly identical to CDK9 in our crystal structure, and shows this similar compound to be a Type I inhibitor. The DFG-loop in the CDK2 structure occupies the “in” conformation (active), but the αC-helix in an “out” (inactive) conformation: this led to exclusion of this structure from our set of comparative modeling templates, since we restricted available templates to those in a fully active kinase conformation (“BLA-minus”) [43]. A comparison of our crystal structure of **14e**/CDK9 to this **14f**/CDK2 structure showed very similar overall orientation of the inhibitors, and identical interactions with the hinge (**Figure 8D**).

In both cases the nitrophenyl group faces into a hydrophobic pocket in the cleft, which is formed in CDK9 by residues Val33, Ala46, the aliphatic part of Lys48, and gatekeeper residue Phe103 (**Figure 8C**). Each of these residues is conserved in CDK2 (Val18, Ala31, Lys33, and Phe80), leading to a structurally-analogous pocket in both kinases. Both inhibitors occupy the same volume in this part of the active site (next to the gatekeeper residue, and adjacent to the hydrophobic “back-pocket” region of the binding site). Given the lipophilicity of the nitro group, interactions beyond packing are unlikely to contribute to determining the orientation of this part of the compound, and both orientations fit the density well. Thus, we cannot conclude unambiguously whether **14e** samples both conformations when bound to CDK9, or just a single conformation.

To evaluate the success of the comparative modeling approach, we superposed our **14e**/CDK9 comparative model onto our crystal structure of this complex (**Figure 8E**). For this comparison, we used from the crystal structure the nitrophenyl orientation that matched the CDK2 structure. Overall, we find an extremely close match between the crystal structure and the comparative model, with root-mean-square deviation (RMSD) of 1.0 Å over the whole structure and 0.83 Å for the ligand (after aligning on the protein only, and using the crystallographic pose with the same stereoisomer and ring orientation as our comparative model). The hydrogen bonds to the hinge region were perfectly recapitulated in our comparative model, and the interactions of the norbornyl group with the region C-terminal of the hinge were virtually identical as well.

### Effect of the norbornane moiety in driving CDK9-selectivity

In our previous work [28], we had attributed the selectivity of **14e** for CDK9 to the norbornyl group. We noted that in crystal structures of CDK9 bound to other ligands as well as in our compound models of the **14e** complex, the loop C-terminal of the hinge was in a slightly lower conformation than in other CDK kinase structures, which we speculated was necessary to accommodate the steric bulk of the spherically-shaped norbornyl group [28]. Accordingly, we anticipated that incorporation of a flat aromatic ring in place of the norbornyl group would result in loss of selectivity for CDK9. To directly test this hypothesis, we revisited our collection of **14e** analogs used in optimization (**Table 3**). Amongst this set, we noted that **14f** (and only this compound) retained the nitro group at the ortho position and replaced the norbornyl group with a flat (substituted) ring (**Figure 8A**). We further noted that this compound showed worse activity that **14e** in our phenotypic assay for reversal of epigenetic silencing, as would be expected if selectivity for CDK9 was lost (**Table 3**).

Before initiating experiments to characterize selectivity of **14f**, however, we recognized that this compound matches exactly the nitrophenyl-aminothiazole ligand solved in the earlier crystal structure with CDK2 (“compound 51” in that paper) [52]. We proceeded to build comparative models of **14f** in complex with each CDK kinase, as a means to access selectivity of this compound (**Figure 9**). To avoid any potential bias from using the previously-solved structure of this compound (or its analogs), models at this stage were built using our newly-solved **14e**/CDK9 structure as the sole template. Comparison of the resulting **14f**/CDK2 model with the crystal structure of this complex confirmed that the pose of this compound was again correctly predicted, with the exception of the same flipped nitrophenyl group observed previously (**Figure 9B**). We expect that this difference in the nitrophenyl group (relative to our previous models) arose from having used our crystal structure as the sole template, because templates that might have led to alternate nitrophenyl orientations were no longer included in our modeling pipeline.

**Figure 9:**
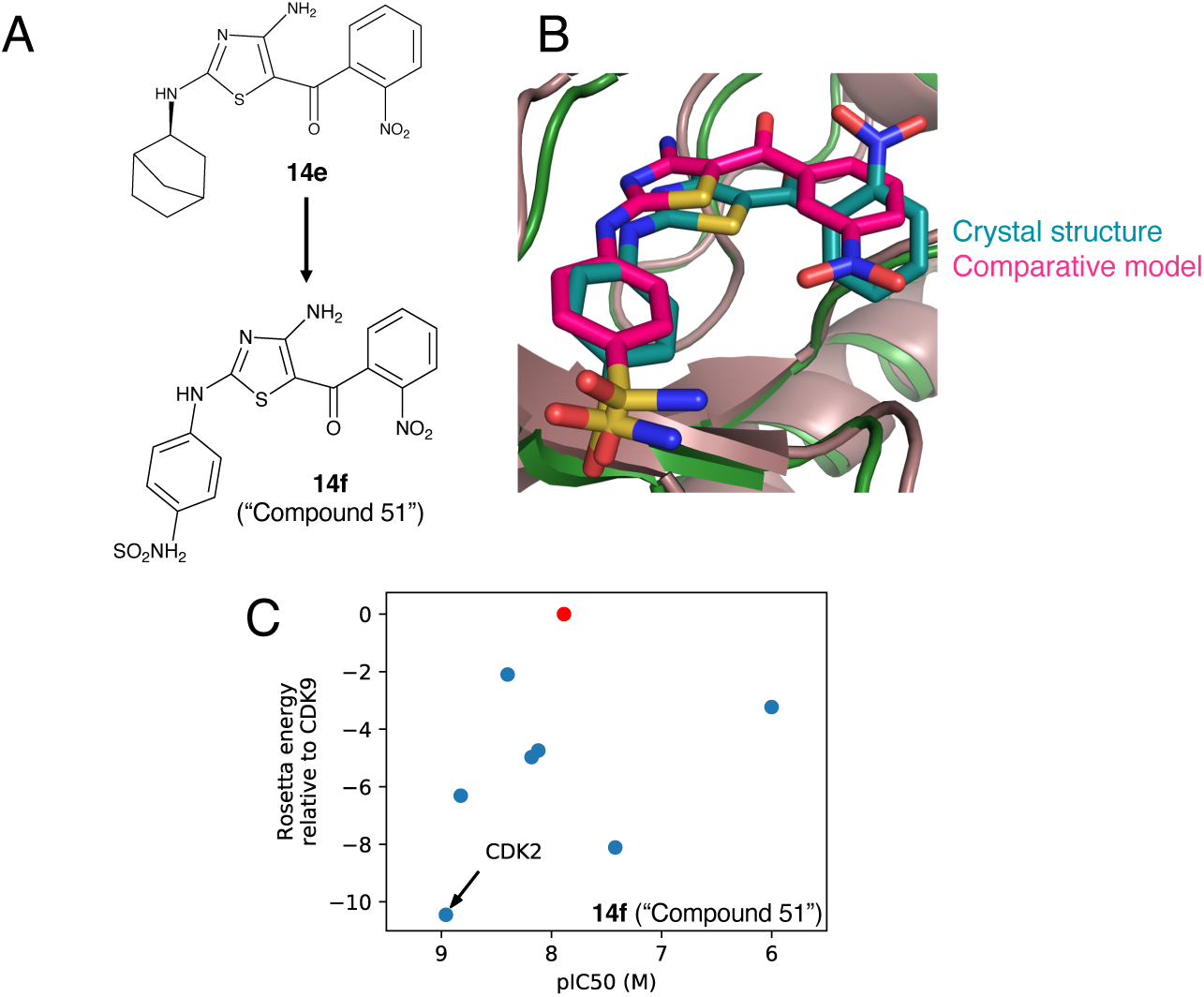
**Probing the basis for 14e selectivity. (A)** Compound **14f** (“compound 51” in Schonbrunn et al. 2013) has a phenylsulfonamide in place of **14e**’s norbornane group. **(B)** Our comparative modeling pipeline provides a prediction of **14f** bound to CDK2 that closely matches the previously-reported crystal structure with CDK2. **(C)** Selectivity profiles from comparative modeling and from kinase assays both identify CDK2 as the preferred target of **14f**, ahead of CDK9.

To our gratification, examination of the study reporting this crystal structure revealed that the selectivity of **14f** had already been defined [52] (**Table 7**). These results confirmed that, consistent with our hypothesis as to the origins of **14e** selectivity, substitution of a flat ring in place of the norbornyl moiety shifted this compound’s preference away from CDK9 and towards CDK2/CDK5. Examining the relative interaction energies of **14f** from the corresponding comparative models showed recapitulation of this preference. In contrast to models of **14e** (**Figure 7**), which ranked CDK9 ahead of any other CDK kinase, the comparative models of **14f** appropriately place CDK2 (and many other CDK kinases) ahead of CDK9 (**Figure 9C**).

**Table 7.** Selectivity of 14f, from comparative modeling and from biochemical kinase assays. In both experiments **14f** is found to exhibit a preference for CDK2 over CDK9. Interaction energies from comparative models are reported in Rosetta Energy Units (REUs).

| Kinase | Interaction energy (REUs) | IC<sub>50</sub> (nM) |
| --- | --- | --- |
| CDK2 | -48.0 | 1.1 |
| CDK3 | -45.6 | 38 |
| CDK5 | -43.8 | 1.5 |
| CDK6 | -42.5 | 6.6 |
| CDK1 | -42.3 | 7.6 |
| CDK7 | -40.8 | >1000 |
| CDK4 | -39.6 | 4.0 |
| CDK9 | -37.5 | 13 |

All IC<sub>50</sub> values are from [52]

In summary, then, the selectivity profile of **14f** strongly supports our earlier assertion that **14e**’s preference for CDK9 derives from its norbornyl moiety. The fact that this conclusion could be reached separately from our earlier (blind) comparative modeling studies [28] and now through analysis of structural analogs supports incorporation of this comparative modeling approach into medicinal chemistry optimization of kinase inhibitors in future.

## Conclusions

In this work, we present SAR around **14e** as interpreted through our novel comparative modeling pipeline. Relative to our earlier comparative modeling strategy, which used only templates from other CDK kinases [28], we have now extended our computational approach to include templates from all structures in the PDB of kinases in their active conformation. The very close structural similarly amongst active conformations of protein kinases [41] – even protein kinases that are distant in phylogeny – enables some of these ostensibly more distant structures to serve as valuable templates; this is especially true for structures that include ligands resembling the inhibitor of interest. By limiting the allowed templates to CDK kinases, our prior approach could not take advantage of templates with clear chemical similarity to **14e**, such as dasatinib: this aminothiazole inhibitor is present in 20 kinase-bound PDB structures, but none with a CDK kinase.

At the same time, however, a surprising observation stemming from our expanded set of templates was that compounds with shared chemical substructures are not necessarily the best templates. Rather, because alignment of the ligands is carried out on the basis of 3D shape and chemical features (i.e., through pharmacophoric matching), our comparative modeling pipeline identifies valuable templates that would not be evident on the basis of their chemical structures. This strength of the approach is highlighted by fact that the template inhibitor yielding the lowest-energy prediction for the **14e**/CDK9 complex used a completely different chemical scaffold from **14e** (**Figure 2**).

An important consequence of broadening the space of potential templates is that this correspondingly expands the space of inhibitors that can be reliably modeled. While direct structural evidence is not available for the majority of reported CDK9 inhibitors, recapitulation of selectivity exhibited by the diverse set of compounds included in our study (**Figure 6**) strongly suggests that our comparative modeling approach has identified and made use of appropriate templates for each of these scaffolds as well. In the case of **14e**, the near-atomic agreement between the model and our crystal structure demonstrates the accuracy that can be attained using diverse templates (**Figure 8**).

A surprising outcome from the medicinal chemistry optimization leading to **14e** was that despite prioritizing only potency, we arrived at a lead compound that was also exquisitely selective across the CDK family (**Table 5**). The close similarity between CDK kinases makes this selectivity unexpected: indeed, most previously-reported inhibitors of CDK9 are notoriously unselective (as seen from the experimentally-derived IC_50_ values in **Figure 7**). We hypothesized that the selectivity of **14e** arose as a by-product of the phenotypic assay used to guide optimization, by steering us away from compounds that inhibit cell-cycle CDKs. The ability of our comparative modeling approach to build models of a given inhibitor with each member of the CDK family allowed us to evaluate each potential interaction pair directly, and ultimately provided evidence supporting our hypothesis (**Figure 5**). Moving forward, we expect that the use of such phenotypic assays may provide a means to simultaneously optimize inhibition of epigenetic kinases such as CDK9, while avoiding potential off-target interactions with cell-cycle kinases.

Intriguingly, it was the SAR from this phenotypic assay that ultimately converged on the use of a norbornane or an alternate puckered ring (tetrahydropyran) at the 2-position of the thiazole ring (**Figure 5**), which we now understand to abut the hinge-adjacent loop of CDK9. This non-obvious choice was enabled by our inadvertent pursuit of selectivity in addition to potency, and indeed it would have been difficult to recognize *a priori* the slight structural difference that distinguishes CDK9 from other CDK family members. Having arrived at **14e**, however, our comparative modeling pipeline was uniquely useful in providing a potential explanation for its selectivity. Ultimately, the models’ attribution of **14e**’s selectivity to its norbornane group is strongly supported by the fact that replacement of this group with a flat ring abrogates its preference for CDK9 (**Figure 9**).

Through these studies, we have found that insights from this comparative modeling approach can rationalize differences in both potency and selectivity of new compounds. We expect that the same approach will prove effective beyond the CDK family, and more broadly to Type I kinase inhibitors in general. Finally, as with any comparative modeling approach, we note that accuracy of these predictions is expected to continuously improve even beyond the current standard, as coverage from the PDB grows to include new potential templates.

## Experimental Section

### Comparative modeling

To build a set of templates, we started from a recent collection of 2573 individual chains of kinases in the PDB in the active (“BLA-minus”) conformation [43]. We restricted this set to those with a ligand in the ATP-binding site, leading to a set of 153 unique kinases. We extracted a single chain from those structures containing multiple chains, leading to a set of 1507 templates. All kinases were aligned to a single reference frame, arbitrarily using PDB ID 1FIN (a CDK2 structure) to define the reference frame. This collection of kinase structures formed the protein template library used in comparative modeling.

The same transformation that brought each kinase into this reference frame was applied to the corresponding ligand from that structure, such that all 1507 ligands were now occupying the same (kinase-bound) reference frame. This collection of aligned ligands formed the ligand template library used in comparative modeling.

For comparative modeling of a given kinase inhibitor (Step I of our pipeline), we used the SMILES string as a starting point for conformer generation with the program OMEGA (version 3.0.1.2) [68, 69]. Using the program ROCS (version 3.2.2.2) [70, 71], we then structurally aligned each of the conformers generated against the crystallographic conformation for each of the 1507 (aligned) ligands in our template library. The aligned matches were evaluated using the ROCS TanimotoCombo score, and the 100 top-scoring alignments were advanced to the next step of modeling.

For comparative modeling of the target kinase (Step II of our pipeline), potential templates were evaluated based on two criteria: sequence similarity of the protein template versus the target kinase, and ligand structural similarity versus the target inhibitor. The rationale for this approach was that a given template may be particularly useful in comparative modeling if it includes a kinase that is similar to the desired kinase (albeit bound to an inhibitor that makes different interactions), or if the template includes an unrelated kinase in complex with an inhibitor that makes similar interactions as the target compound. An ideal template, of course, would include a kinase that is closely related to the desired kinase, in complex with an inhibitor that engages the kinase in a very similar way as the target compound does.

To evaluate sequence similarity of the protein template versus the target kinase, we used the standalone EMBOSS Needleman-Wunsch program (version 6.6.0.0) [72, 73]. All available templates in our library were ranked in order of similarity to the desired kinase (“sequence rank”). To evaluate the similarity (in three-dimensional geometry) of a template’s ligand to that of the desired ligand, we looked up the TanimotoCombo score that was calculated in Step I of our pipeline when evaluating the potential overlay of the target inhibitor with the template ligand. All available templates in our library were ranked in order of their ligand’s similarity to the desired ligand (“ligand rank”). For each template, we then multiplied its sequence rank by its ligand rank to give an overall score to this template. For each of the top-scoring 10 templates by this combined ranking, we then used the PyRosetta software (PyRosetta4.Release.python36.linux.release-191) [74] to update the sequence in the protein structure by mutating any residues needed so that the current sequence matched the desired sequence (since the protein template may have come from a different kinase than the target kinase).

The final modeling building and refinement (Step III of our pipeline), was carried out using PyRosetta pose manipulation protocols [74]. For the single top-scoring protein template in Step II, we superposed onto the protein structure each of the 100 top-scoring ligand conformations from Step I. No further alignment was needed at this stage, since both the kinases and the ligands were already oriented into the same reference frame when we built the template libraries. Each of these 100 conformations was subjected to full-atom gradient-based energy minimization using Rosetta [75]. Any models that did not maintain canonical hydrogen bonding interactions between the ligand and the kinase hinge region were removed, then the remaining models were ranked on the basis of protein-ligand interaction energy. Each of the 10 top-scoring ligand poses were then superposed into all 10 protein structures from Step II, and the minimization step was repeated for these 100 complexes. The resulting models were again filtered to ensure canonical hydrogen bonding interactions at the hinge region, then the top-scoring model (on the basis of protein-ligand interaction energy) was taken as the final prediction for this complex.

Through the use of Rosetta in studying a variety of protein-ligand complexes, we noted that Rosetta’s default energy function is inappropriately parametrized with respect to nitro groups. By default, these are modeled as analogous to carboxyl groups, whereas in reality nitro groups are much more lipophilic than carboxyl groups, and (unlike carboxyl groups) they rarely participate in hydrogen bonds with their environment [76]. This oversight has not yet been widely appreciated, in part because the vast majority of Rosetta’s use has been focused on modeling proteins. To enable modeling of compounds containing nitro groups, we instead use a variation of the Rosetta energy function in which parameters are modified such that the oxygens in a nitro group are not considered to be hydrogen bond acceptors.

### Biology

#### Cellular fluorescence assay

YB5 cell line (derived from SW48 (female) colon cancer cell line) was used to detect GFP reactivation after drug treatments. YB5 cells were maintained in L-15 (Corning) supplemented with 10% FBS (Atlanta Biologicals) and 1% penicillin/streptomycin (Corning) at 1% CO_2_ and 37°C. For this assay, a single-dose, 24-hour treatment schedule was performed. A total of 10000 cells per well were analyzed using Millipore Guava flow cytometry (EMD, Millipore). GFP-positive percentage was calculated by excluding all propidium iodide-positive cells. All analogs were prepared as stocks in 100% DMSO. After dilution, the final concentration of DMSO in drug-treated cultures was 0.5%. Depsipeptide (an HDAC inhibitor, dosed at 40 nM) was used as a positive control and as a standard to calculate relative activities of each newly identified analogs.

#### In vitro kinase assay

CDK enzymatic assays were performed by Reaction Biology Corp. (Malvern PA), using 10 μM ATP for determining IC_50_’s of **14e** with each CDK family member.

### X-ray Crystallography

#### Crystallization and structure determination

The pFastBac plasmid for CDK9-Cyclin-T1 co-expression was a gift from Dr. Ursula Schulze-Gahmen. The plasmid is a modified version of pFastBac Dual, with a TEV cleavable His-Tag at the p10 promoter. CDK9 1-330 is cloned at the P10 promoter and Cyclin-T1 1-264 at the PH promoter. In order to get the best crystallization result, we introduced Q77R, E96G, F241L into CDK9 and truncated 5aa from Cyclin-T1 [77] using a site-directed mutagenesis method. pFastBac plasmids were transformed into Bac10 cells to generate and screen for positive bacmids, and the confirmed bacmids were transfected into *Sf*9 cells with Invitrogen Cellfectin II reagent to produce viruses. For protein production, *Sf*9 insect cells were grown in ESF 921 media and infected with P3 viruses, the cells were pelleted after 48 hours of infection. Cells were lysed in 50 mM Tris pH 8.0, 500 mM NaCl, protein sample was extracted from the supernatant with a HisTrap^TM^ HP column (GE healthcare) and eluted with an imidazole gradient. The His-tag was removed by TEV protease, untagged protein complex was further purified by a Resource Q column.

CDK9/Cyclin-T1 protein was crystallized to 4 mg/ml, in a well solution containing 0.1 M Na/K phosphate pH 6.2, 200 mM NaCl, 4 mM TCEP and 1∼6% PEG1000 at 4°C. The crystals were soaked in 1 mM MC180295 for more than 1 hour and flash frozen in the well solution with additional 30% ethylene glycol. X-ray diffraction data was collected at NSLS-II AMX beamline. The diffraction data were indexed, integrated and scaled with FastDP [78–81]. The crystal structure was determined by molecular replacement method in Phenix [82] and the model was refined using the CCP4 suite [83]. Data collection and refinement statistics are listed in **Table 6**.

### Chemistry

#### General Methods

Reagents and starting materials were purchased from commercial vendors and used without further purification unless otherwise indicated. Reactions were followed by thin layer chromatography and liquid chromatography/mass spectrometry (LC/MS). ^1^H-NMR spectra and ^13^C-NMR spectra were recorded on a Bruker Avance III spectrometer at ambient temperature at 400 MHz and 100 MHz, respectively. Chemical shifts are reported in ppm relative to CDCl_3_ (7.26 ppm for ^1^H-NMR and 77.0 ppm for ^13^C-NMR). LC-MS analysis was performed on an Agilent Technologies 1200 series LC system coupled to a 6300 quadrapole MS. High resolution mass spectrometry analysis was performed by the Mass Spectrometry Lab, School of Chemical Sciences, University of Illinois at Urbana-Champaign on a Waters QTOF Ultima ESI instrument. Flash chromatography was performed on ISCO RediSep Rf^®^ normal phase silica columns using a Teledyne ISCO CombiFlash Instrument. Purity was measured by high pressure liquid chromatography using two different systems with UV detection at 210 and 254 nM: normal phase on silica using gradients of ethyl acetate in hexanes; reversed phase on C-18 using gradients of acetonitrile in water with trifluoroacetic acid as a modifier. All compounds were determined to be > 98.5% pure unless otherwise noted.

### Synthesis of Compounds 7, 8a – 8m

**(3,4-Dimethoxyphenyl)(2-(methylthio)thiazol-5-yl)methanone (7)**. A solution of 2-(methylthio)thiazole (392 uL, 3.81 mmol) in anhydrous tetrahydrofuran (7.5 mL) was cooled to −55°C under nitrogen. N-Butyl lithium (1.6 M in hexanes, 2.38 mL, 3.81 mmol) was added dropwise, keeping the reaction solution temperature below −40°C. A solution of 3,4-dimethoxybenzonitrile (311 mg, 1.91 mmol) in anhydrous tetrahydrofuran (2.5 mL) was then added dropwise while keeping the reaction temperature between −35°C and −20°C. Stirring was continued at this temperature range for 50 minutes. The reaction was quenched with 1M aqueous acetic acid (11.4 mL, 11.43 mmol) at −20°C. The reaction was allowed to warm to room temperature with stirring overnight and then diluted with ethyl acetate and water. The aqueous phase was extracted with two additional portions of ethyl acetate. The combined organic layers were washed with 50% aqueous potassium carbonate solution twice and brine, dried over anhydrous sodium sulfate and concentrated on a rotary evaporator. The crude product was purified by chromatography on silica gel using a gradient of 0 to 80% of ethyl acetate in hexanes to afford the titled compound as an orange-yellow solid (212 mg, 19%).^1^H NMR (400 MHz, CDCl_3_) δ 8.08 (s, 1H), 7.53 (dd, J = 8.32 Hz, J = 2.0 Hz, 1H), 7.43 (d, J = 2.0 Hz, 1H), 6.94 (d, J = 8.36 Hz, 1H), 3.98 (s, 3H), 3.95 (s, 3H), 2.76 (s, 3H); MS (ESI): *m/z* 295.7 [(M+H)^+^].

**Method A.** The synthesis of compound **8a** exemplifies General Method A.

**(4-Amino-2-(butylamino)thiazol-5-yl)(3,4-dimethoxyphenyl)methanone (8a).** Butyl isothiocyanate (24.5 uL, 0.20 mmol) and solid potassium tert-butoxide (50.2 mg, 0.45 mmol) were added sequentially to a solution of cyanamide (8.5 mg, 0.20 mmol) in anhydrous tetrahydrofuran (1.0 mL). The mixture was stirred for 15 minutes. A solution of 2-bromo-1-(3,4-dimethoxyphenyl)ethanone (50 mg, 0.19 mmol) in anhydrous tetrahydrofuran (500 uL) was then added. The resulting solution was stirred at room temperature overnight. The reaction was concentrated on a rotary evaporator and the residue was partitioned between ethyl acetate and water. The organic layer was washed with brine, dried over anhydrous sodium sulfate and concentrated on a rotary evaporator. The crude product was purified by chromatography on silica gel using a gradient of 0 to 100% of ethyl acetate in hexanes to afford the titled compound as a light yellow solid (10 mg, 15%). ^1^H NMR (400 MHz, CDCl_3_) δ 7.38 (m, 2H), 6.87 (d, J = 8.28 Hz, 1H), 5.72 (bs, 1H), 3.93 (d, J = 1.20 Hz, 6H), 3.25 (q, J = 6.80, 2H), 1.63 (m, 2H), 1.40 (m, 2H), 0.95 (t, J = 7.35 HZ, 2H); ^13^C NMR (100 MHz, CDCl_3_) δ 190.3, 165.4, 158.7, 150.8, 143.5, 139.9, 131.1, 126.2, 114.7, 115.3, 58.9, 58.7, 45.6, 28.8, 21.1, 18.9; MS (ESI): *m/z* 336.1 [(M+H)^+^].

The following compounds were prepared using similar methodology:

**(4-Amino-2-(cyclopentylamino)thiazol-5-yl)(3,4-dimethoxyphenyl)methanone (8b)**. General Method A was followed using cyclopentyl isothiocyanate (25.1 uL, 0.20 mmol), potassium tert-butoxide (50.2 mg, 0.45 mmol), cyanamide ((8.5 mg, 0.20 mmol) and 2-bromo-1-(3,4-dimethoxyphenyl)ethanone (50 mg, 0.19 mmol). The desired product was isolated as an orange-yellow solid (19 mg, 14%). ^1^H NMR (400 MHz, CDCl_3_) δ 7.39 (m, 2H), 6.88 (d, J = 8.28 Hz, 1H), 5.62 (bd, J = 7.28, Hz,1H), 3.93 (s, 6H), 3.82 (m, 1H), 2.05 (m, 2H), 1.65 (m, 6H); ^13^C NMR (100 MHz, CDCl_3_) δ 190.0, 166.1, 157.9, 149.9, 143.7, 138.8, 131.0, 126.4, 113.9, 113.3, 58.8, 58.3, 56.7, 33.5, 24.1; MS (ESI): *m/z* 348.1 [(M+H)^+^].

**(4-Amino-2-(cyclohexylamino)thiazol-5-yl)(3,4-dimethoxyphenyl)methanone (8c)**. General Method A was followed using cyclohexyl isothiocyanate (27.8 uL, 0.20 mmol), potassium tert-butoxide (50.2 mg, 0.45 mmo.l), cyanamide ((8.5 mg, 0.20 mmol) and 2-bromo-1-(3,4-dimethoxyphenyl)ethanone (50 mg, 0.19 mmol). The desired product was isolated as an orange-yellow glassy solid (12.1 mg, 9%). ^1^H NMR (400 MHz, CDCl_3_) δ 7.38 (m, 2H), 6.88 (d, J = 8.28 Hz, 1H), 5.59 (bd, J = 7.92 Hz, 1H), 3.93 (s, 6H), 3.82 (m, 1H), 2.05 (m, 2H), 1.76 (m, 2H), 1.61 (m, 1H), 1.31 (m, 5H); ^13^C NMR (100 MHz, CDCl_3_) δ 190.1, 166.2, 157.7, 150.1, 143.5, 138.7, 130,.9, 126.2, 113.8, 113.5, 58.9, 58.5, 54.9, 33.3, 26.1, 25.7; MS (ESI): *m/z* 362.2 [(M+H)^+^].

**(4-Amino-2-((methoxymethyl)amino)thiazol-5-yl)(3,4-dimethoxyphenyl)methanone (8d)**. General Method A was followed using methoxymethyl isothiocyanate (18.9 uL, 0.20 mmol), potassium tert-butoxide (50.2 mg, 0.45 mmol), cyanamide ((8.5 mg, 0.20 mmol) and 2-bromo-1-(3,4-dimethoxyphenyl)ethanone (50 mg, 0.19 mmol). The desired product was isolated as an orange solid (7.7 mg, 12%). ^1^H NMR (400 MHz, CDCl_3_) δ 7.40 (m, 1H), 7.35 (d, J = 2.00 Hz, 1H), 6.88 (d, J = 8.32 Hz, 1H), 4.73 (d, J = 6.16 Hz, 2H), 3.93 (s, 6H), 3.38 (s, 3H); ^13^C NMR (100 MHz, CDCl_3_) δ 190.1, 165.9, 155.8, 149.9, 143.1, 138.6, 131.0, 126.3, 113.8, 113.4, 85.2, 58.7, 58.2, 56.3; MS (ESI): *m/z* 669.2 [(2M+Na)^+^].

**(4-Amino-2-(benzylamino)thiazol-5-yl)(3,4-dimethoxyphenyl)methanone (8e).** General Method A was followed using benzyl isothiocyanate (26.9 uL, 0.20 mmol), potassium tert-butoxide (50.2 mg, 0.45 mmol), cyanamide ((8.5 mg, 0.20 mmol) and 2-bromo-1-(3,4-dimethoxyphenyl)ethanone (50 mg, 0.19 mmol). The desired product was isolated as an orange-yellow glassy solid (17.3 mg, 24%). ^1^H NMR (400 MHz, CDCl_3_) δ 7.35 (m, 7H), 6.86 (d, J = 8.36 Hz, 1H), 6.41 (bs, 1H), 4.46 (bd, J = 4.24 Hz, 2H), 3.92 (s, 3H), 3.90 (s, 3H); ^13^C NMR (100 MHz, CDCl_3_) δ 190.0, 166.1, 156.1, 150.3, 144.0, 159.8, 141.2, 131.9, 128.9, 126.9, 126.6, 126.4, 113.8, 113.3, 58.9, 58.5. 50.7; MS (ESI): *m/z* 370.1 [(M+H)^+^].

**(4-Amino-2-((4-methoxybenzyl)amino)thiazol-5-yl)(3,4-dimethoxyphenyl)methanone (8f)**. General Method A was followed using 4-methoxybenzyl isothiocyanate (31.7 uL, 0.20 mmol), potassium tert-butoxide (50.2 mg, 0.45 mmol), cyanamide ((8.5 mg, 0.20 mmol) and 2-bromo-1-(3,4-dimethoxyphenyl)ethanone (50 mg, 0.19 mmol). The desired product was isolated as an orange-yellow glassy solid (12.6 mg, 16%). ^1^H NMR (400 MHz, CDCl_3_) δ 7.38 (m, 2H), 7.26 (m, 2H), 6.88 (m, 3H), 4.39 (bs, 2H), 3.92 (s, 3H), 3.91 (s, 3H), 3.81 (s, 3H);^13^C NMR (100 MHz, CDCl_3_) δ 190.0, 165.7, 160.1, 155.7, 152.4, 144.0, 139.2, 133.5, 132.1, 130.8, 126.5, 115.5, 113.7, 113.4, 58.9, 58.4, 57.0, 51.3; MS (ESI): *m/z* 400.1 [(M+H)^+^].

**(4-Amino-2-(phenethylamino)thiazol-5-yl)(3,4-dimethoxyphenyl)methanone (8g)**. General Method A was followed using 4-methoxybenzyl isothiocyanate (30.4 uL, 0.20 mmol), potassium tert-butoxide (50.2 mg, 0.45 mmol), cyanamide ((8.5 mg, 0.20 mmol) and 2-bromo-1-(3,4-dimethoxyphenyl)ethanone (50 mg, 0.19 mmol). The desired product was isolated as an orange-yellow solid (16.9 mg, 23%). ^1^H NMR (400 MHz, CDCl_3_) δ 7.41 (m, 1H), 7.36 (m, 2H), 7.28 (m, 2H), 7.22 (m, 2H), 6.90 (d, J = 8.28 Hz, 1H), 3.95 (s, 6H), 3.57 (m, 2H), 2.98 (t, J = 6.84 Hz, 2H); ^13^C NMR (100 MHz, CDCl_3_) δ 190.0, 165.3, 155.0, 150.2, 144.1, 142.7, 139.5, 113.6, 113.3, 130.9, 128.1, 127.8, 126.6, 125.7, 58.6, 58.2, 46.3, 37.1; MS (ESI): *m/z* 384.1 [(M+H)^+^].

**(4-Amino-2-((4-methoxyphenethyl)amino)thiazol-5-yl)(3,4-dimethoxyphenyl)methanone (8h)**. General Method A was followed using 4-methoxyphenethyl isothiocyanate (37.4 uL, 0.20 mmol), potassium tert-butoxide (50.2 mg, 0.45 mmol), cyanamide ((8.5 mg, 0.20 mmol) and 2-bromo-1-(3,4-dimethoxyphenyl)ethanone (50 mg, 0.19 mmol). The desired product was isolated as an orange-yellow solid (16,2 mg, 20%). ^1^H NMR (400 MHz, CDCl_3_) δ 7.38 (m, 2H), 7.11 (m, 2H), 6.87 (m, 3H), 3.93 (s, 6H), 3.79 (s, 3H), 3.50 (q, J = 6.12 Hz, 2H), 2.89 (t, J = 6.80 Hz, 2H); ^13^C NMR (100 MHz, CDCl_3_) δ 190.1, 165.6, 159.4, 155.3, 150.4, 143.9, 142.4, 132.7, 116.0, 113.9, 113.6, 127.9, 126.9, 125.4, 58.8, 58.5, 56.3, 45.3, 36.8; MS (ESI): *m/z* 414.1 [(M+H)^+^].

**(4-Amino-2-(benzo[d][1,3]dioxol-5-ylamino)thiazol-5-yl)(3,4-dimethoxyphenyl)methanone (8i)**. General Method A was followed using benzyl isothiocyanate (29.7 uL, 0.20 mmol), potassium tert-butoxide (50.2 mg, 0.45 mmol), cyanamide ((8.5 mg, 0.20 mmol) and 2-bromo-1-(3,4-dimethoxyphenyl)ethanone (50 mg, 0.19 mmol). The desired product was isolated as an orange-yellow solid (12.9 mg, 17%). ^1^H NMR (400 MHz, CDCl_3_) δ 7.35 (m, 2H), 6.92 (d, J = 1.92 Hz, 1H), 6.86 (d, J = 8.20 Hz, 1H), 6.78 (m, 2H), 6.01 (s, 2H), 3.92 (s, 6H); ^13^C NMR (100 MHz, CDCl_3_) δ 190.2, 160.9, 155.2, 150.8, 150.4, 143.6, 140.3, 139.7, 137.6, 130.1, 124.2, 115.0, 113.7, 113.3, 109.7, 107.8, 102.7, 58.7, 58.3; MS (ESI): *m/z* 400.0 [(M+H)^+^].

**(4-Amino-2-((1-phenylethyl)amino)thiazol-5-yl)(3,4-dimethoxyphenyl)methanone (8j).** General Method A was followed using 1-phenylethyl isothiocyanate (31.3 uL, 0.20 mmol), potassium tert-butoxide (50.2 mg, 0.45 mmol), cyanamide ((8.5 mg, 0.20 mmol) and 2-bromo-1-(3,4-dimethoxyphenyl)ethanone (50 mg, 0.19 mmol). The desired product was isolated as an orange-yellow glassy solid (30.2 mg, 41%). ^1^H NMR (400 MHz, CDCl_3_) δ 7.32 (m, 7H), 6.85(d, J = 8.32 Hz, 1H), 6.19 (bs, 1H), 4.63 (bs, 1H), 3.92 (s, 3H), 3.86 (s, 3H), 1.60 (d, J = 6.80 Hz, 3H); ^13^C NMR (100 MHz, CDCl_3_) δ 190.0, 164.9, 154.8, 150.3, 144.0, 142.5, 138.4, 128.9, 128.7, 127.3, 127.1, 123.6, 113.6, 113.3, 60.9, 58.7, 58.4, 22.3; MS (ESI): *m/z* 384.1 [(M+H)^+^].

**(R)-(4-Amino-2-((1-(4-methoxyphenyl)ethyl)amino)thiazol-5-yl)(3,4-dimethoxyphenyl)methanone (8k)**. General Method A was followed using (R)-1-phenylethyl isothiocyanate (37.4 uL, 0.20 mmol), potassium tert-butoxide (50.2 mg, 0.45 mmol), cyanamide ((8.5 mg, 0.20 mmol) and 2-bromo-1-(3,4-dimethoxyphenyl)ethanone (50 mg, 0.19 mmol). The desired product was isolated as an orange-yellow glassy solid (23.3 mg, 29%). ^1^H NMR (400 MHz, CDCl_3_) δ 7.34 (dd, J = 2.00 Hz, J = 8.24 Hz, 1H), 7.29 (d, J = 2.00 Hz, 1H), 7.23 (m, 2H), 6.86 (m, 3H), 6.06 (bd, J = 6.32 Hz, 1H), 4.58 (m, 1H), 3.92 (s, 3H), 3.87 (s, 3H), 3.79 (s, 3H), 1.57 (d, J = 6.76 Hz, 3H); ^13^C NMR (100 MHz, CDCl3) δ 190.1, 164.9, 159.7, 155.0, 150.8, 143.7, 138.1, 136.9, 129.6, 127.2, 124.4, 115.5, 113.8, 113.5, 61.0, 58.7, 58.5, 56.9, 22.4; MS (ESI): *m/z* 414.1 [(M+H)^+^].

**(S)-(4-Amino-2-((2,3-dihydro-1H-inden-1-yl)amino)thiazol-5-yl)(3,4-dimethoxyphenyl)methanone (8l)**. General Method A was followed using (S)-1-indanyl isothiocyanate (30.7 uL, 0.20 mmol), potassium tert-butoxide (50.2 mg, 0.45 mmol), cyanamide ((8.5 mg, 0.20 mmol) and 2-bromo-1-(3,4-dimethoxyphenyl)ethanone (50 mg, 0.19 mmol). The desired product was isolated as an orange-yellow glassy solid (24.4 mg, 32%). ^1^H NMR (400 MHz, CDCl_3_) δ 7.23-7.42 (m, 6H), 6.86 (d, J = 8.32 Hz, 1H), 5.04 (bd, J = 7.04 Hz, 1H), 3.92 (m, 7H), 3.04 (m, 1H), 2.88 (m, 1H), 2.68 (m, 1H), 1.94 (m, 1H); ^13^C NMR (100 MHz, CDCl_3_) δ 190.1, 164.5, 154.9, 150.6, 144.2, 143.0, 141.8, 138.0, 129.2, 127.0, 126.9, 125.3, 125.1, 123.9, 113.6, 113,.3, 69.9, 58.8, 58.5, 34.7, 30.4; MS (ESI): *m/z* 396.1 [(M+H)^+^].

**(2-((1,4-Dioxaspiro[4.5]decan-8-yl)amino)-4-aminothiazol-5-yl)(3,4-dimethoxyphenyl)methanone (8m**). General Method A was followed using 8-isothiocyanato-1,4-dioxaspiro[4,5]decane (31.1 uL, 0.20 mmol) [46], potassium tert-butoxide (50.2 mg, 0.45 mmol), cyanamide ((8.5 mg, 0.20 mmol) and 2-bromo-1-(3,4-dimethoxyphenyl)ethanone (50 mg, 0.19 mmol). The desired product was isolated as an orange-yellow glassy solid (12.8 mg, 6 %). ^1^H NMR (400 MHz, CDCl_3_) 7.38 (dd, J = 2.00 Hz, J = 8.20 Hz, 1H), 7.35 (d, J = 1.96 Hz, 1H), 6.88 (m, 1H), 5.90 (bs, 1H), 3.95 (bt, J = 2.24 Hz, 4H), 3.93 (s, 6H), 3.90 (bs, 1H), 2.08 (m, 2H), 1.79 (m, 2H), 1.66 (m, 4H); ^13^C NMR (100 MHz, CDCl_3_) δ 190.2, 164.7, 155.0, 150.9, 143.2, 138.1, 129.2, 123.8, 118.8, 113.7, 113.3, 64.9, 58.7, 58.4, 56.0, 29.7, 22.5 ; MS (ESI): *m/z* 420.1 [(M+H)^+^].

### Synthesis of compounds 10, 12a – 12an

**4-Amino-2-(methylthio)thiazol-5-yl)(2,3-dihydrobenzo[b][1,4]dioxin-6-yl)methanone (10).** 2-Bromo-1-(2,3-dihydro-1,4-benzodioxin-6-yl)ethan-1-one (100 mg, 0.3890 mmol) and triethylamine (70 uL, 0.50 mmol) were added sequentially to a solution of cyanimidodithiocarbonic acid S-methyl ester S-potassium salt (59.5 mg, 0.35 mmol) in anhydrous dimethylformamide (2.0 mL). The resulting mixture was stirred at 80°C for 3 hours. The reaction was cooled to room temperature and concentrated on a rotary evaporator. The residue was partitioned between ethyl acetate and water. The organic layer was washed with brine, dried over anhydrous sodium sulfate and concentrated on a rotary evaporator. The crude product was purified by chromatography on silica gel using a gradient of 0 to 100% of ethyl acetate in hexanes to afford compound the desired compound as a yellow solid (77 mg, 71%). ^1^H NMR (400 MHz, CDCl_3_) δ 7.33 (m, 2H), 6.90 (d, J = 8.28 Hz, 1H), 4.30 (m, 4H), 2.66 (s, 3H); ^13^C NMR (100 MHz, CDCl_3_) δ 189.9, 166.6, 155.5, 150.4, 147.9, 147.3, 128.3, 122.5, 112.5, 112.2, 65.6, 65.4, 19.8; MS (ESI): *m/z* 309.1 [(M+H)^+^].

**Method B.** The synthesis of compound **12a** exemplifies General Method B.

**(4-amino-2-(methylamino)thiazol-5-yl)(2,3-dihydrobenzo[b][1,4]dioxin-6-yl)methanone (12a)**. Methyl isothiocyanate (14.0 µL, 021 mmol) and solid potassium tert-butoxide (50.5 mg, 0.45 mmol) were added sequentially to a solution of cyanamide (8.6 mg, 0.21 mmol) in anhydrous tetrahydrofuran (1.0 mL). This mixture was stirred for 15 minutes. A solution of 2-bromo-1-(2,3-dihydro-1,4-benzodioxin-6-yl)ethan-1-one (50 mg, 0.20 mmol) in anhydrous tetrahydrofuran (500 µL) was added. The resulting orange solution was stirred at room temperature overnight. The reaction was concentrated on a rotary evaporator and the residue was partitioned between ethyl acetate and water. The organic layer was washed with brine, dried over anhydrous sodium sulfate and concentrated on a rotary evaporator. The crude product was purified by chromatography on silica gel using a gradient of 0 to 100% of ethyl acetate in hexanes to afford the titled compound as an orange-yellow oil (4.1 mg, 7%). ^1^H NMR (400 MHz, CD_3_OD) δ 7.19 (m, 2H), 6.87 (m, 1H), 4.28 (m, 4H), 2.94 (s, 3H); ^13^C NMR (100 MHz, CDCl_3_) δ 189.9, 165.1, 155.8, 150.1, 143.3,138.2, 128.4, 122.9, 112.9, 112.6, 65.4, 65.2, 31.4; MS (ESI): *m/z* 292.1 [(M+H)^+^].

The following compounds were prepared using a similar procedure:

**(4-Amino-2-(ethylamino)thiazol-5-yl)(2,3-dihydrobenzo[b][1,4]dioxin-6-yl)methanone (12b)**. General Method B was followed using ethyl isothiocyanate (17.9 µL, 0.21 mmol), potassium tert-butoxide (50.5 mg, 0.45 mmol), cyanamide (8.6 mg, 0.21 mmol) and 2-bromo-1-(2,3-dihydro-1,4-benzodioxin-6-yl)ethan-1-one (50 mg, 0.20 mmol). The desired product was isolated as an orange-yellow oil (10.9 mg, 18%). ^1^H NMR (400 MHz, CDCl_3_) δ 7.31 (m, 2H), 6.88 (m, 1H), 4.29 (m, 4H), 3.30 (m, 2H), 1.29 (t, J = 7.20 Hz, 3H); ^13^C NMR (100 MHz, CDCl_3_) δ 190.0, 165.2, 155.7, 150.3, 143.1,138.5, 128.7, 123.2, 112.8, 112.4, 65.5, 65.4, 31.1, 17.6; MS (ESI): *m/z* 306.1 [(M+H)^+^].

**(4-Amino-2-(propylamino)thiazol-5-yl)(2,3-dihydrobenzo[b][1,4]dioxin-6-yl)methanone (12c)** General Method B was followed using propyl isothiocyanate (21.2 µL, 0.21 mmol), potassium tert-butoxide (50.5 mg, 0.45 mmol), cyanamide (8.6 mg, 0.21 mmol) and 2-bromo-1-(2,3-dihydro-1,4-benzodioxin-6-yl)ethan-1-one (50 mg, 0.20 mmol). The desired product was isolated as an orange-yellow glassy solid (8.9 mg,14%). ^1^H NMR (400 MHz, CD_3_OD) δ 7.19 (m, 2H), 6.87 (m. 1H), 4.28 (m, 4H), 3.30 (m, 2H), 1.65 (m, 2H), 0.98 (t, J = 7.40 Hz, 3H); ^13^C NMR (100 MHz, CDCl_3_) δ 189.9, 165.2, 155.7, 150.0, 143.1,138.3, 128.2, 123.0, 112.7, 112.4, 65.3, 65.1, 45.8, 23.2, 16.8 ; MS (ESI): *m/z* 320.1 [(M+H)^+^].

**(4-Amino-2-(isopropylamino)thiazol-5-yl)(2,3-dihydrobenzo[b][1,4]dioxin-6-yl)methanone (12d)** General Method B was followed using isopropyl isothiocynate (21.8 µL, 0.21 mmol), potassium tert-butoxide (50.5 mg, 0.45 mmol), cyanamide (8.6 mg, 0.21 mmol) and 2-bromo-1-(2,3-dihydro-1,4-benzodioxin-6-yl)ethan-1-one (50 mg, 0.20 mmol). The desired product was isolated as a yellow solid (4.0 mg, 6%). ^1^H NMR (400 MHz, CD_3_OD) δ 7.18 (m, 1H), 6.86 (m, 1H), 4.27 (m, 4H), 3.30 (m, 1H), 1.25 (s, 3H), 1.24 (s, 3H); ^13^C NMR (100 MHz, CDCl_3_) δ 190.0, 165.3, 155.7, 150.2, 143.1,138.3, 128.2, 122.8, 112.9, 112.6, 65.5, 65.4, 51.5, 23.4 ; MS (ESI): *m/z* 320.1 [(M+H)^+^].

**(4-Amino-2-(butylamino)thiazol-5-yl)(2,3-dihydrobenzo[b][1,4]dioxin-6-yl)methanone (12f)** General Method B was followed using butyl isothiocynate (24.7 µL, 0.21 mmol), potassium tert-butoxide (50.5 mg, 0.45 mmol), cyanamide (8.6 mg, 0.21 mmol) and 2-bromo-1-(2,3-dihydro-1,4-benzodioxin-6-yl)ethan-1-one (50 mg, 0.20 mmol). The desired product was isolated as a yellow solid (5.4 mg, 8%). ^1^H NMR (400 MHz, CD_3_OD) δ 7.18 (m, 2H), 6.86 (m, 1H), 4.27 (m, 4H), 3.30 (m, 2H), 1.60 (m, 2H), 1.40 (m, 2H), 0.96 (t, J = 7.32 Hz, 3H); ^13^C NMR (100 MHz, CDCl_3_) δ 190.1, 165.2, 155.6, 150.0, 143.0,138.1, 128.1, 123.0, 112.7, 112.4, 65.5, 65.3, 44.0, 27.5, 22.8, 15.7; MS (ESI): *m/z* 334.1 [(M+H)^+^].

**(4-Amino-2-(cyclopentylamino)thiazol-5-yl)(2,3-dihydrobenzo[b][1,4]dioxin-6-yl)methanone (12h)** General Method B was followed using cyclopentyl isothiocyanate (23.5 µL, 0.21 mmol), potassium tert-butoxide (50.5 mg, 0.45 mmol), cyanamide (8.6 mg, 0.21 mmol) and 2-bromo-1-(2,3-dihydro-1,4-benzodioxin-6-yl)ethan-1-one (50 mg, 0.20 mmol). The desired product was isolated as a yellow solid (9.2 mg, 14%). ^1^H NMR (400 MHz, CDCl_3_) δ 7.31 (m, 2H), 6.88 (d, J = 8.28 Hz, 1H), 5.67 (m, 1H), 4.29 (m, 4H), 3.80 (m, 1H), 2.04 (m, 2H), 1.50-1.80 (m, 6H); ^13^C NMR (100 MHz, CDCl_3_) δ 190.1, 164.9, 155.6, 150.1, 143.0,138.3, 128.3, 122.8, 112.8, 112.5, 65.6, 65.5, 55.9, 32.7, 24.3; MS (ESI): *m/z* 346.1 [(M+H)^+^].

**(4-Amino-2-(cyclohexylamino)thiazol-5-yl)(2,3-dihydrobenzo[b][1,4]dioxin-6-yl)methanone (12i)** General Method B was followed using cyclohexyl isothiocyanate (29.0 µL, 0.21 mmol.), potassium tert-butoxide (50.5 mg, 0.45 mmol.), cyanamide (8.6 mg, 0.21 mmol.) and 2-bromo-1-(2,3-dihydro-1,4-benzodioxin-6-yl)ethan-1-one (50 mg, 0.20 mmol.). The desired product was isolated as an orange-yellow glassy solid (3.8 mg, 5%). ^1^H NMR (400 MHz, CDCl_3_) δ 7.30 (m, 2H), 6.88 (d, J = 8.28 Hz, 1H), 5.62 (m, 1H), 4.29 (m, 4H), 3.30 (m, 1H), 2.04 (m, 2H), 1.78 (M, 2H), 1.20-1.42 (m, 6H); ^13^C NMR (100 MHz, CDCl_3_) δ 190.1, 165.4, 155.6, 149.9, 143.0,138.4, 128.3, 123.0, 112.8, 112.5, 65.5, 65.3, 55.0, 33.5, 25.7, 25.3; MS (ESI): *m/z* 360.1 [(M+H)^+^].

**(4-Amino-2-((methoxymethyl)amino)thiazol-5-yl)(2,3-dihydrobenzo[b][1,4]dioxin-6-yl)methanone (12j)** General Method B was followed using methoxymethyl isothiocyanate (20.5 µL, 0.21 mmol.), potassium tert-butoxide (50.5 mg, 0.4504 mmol.), cyanamide (8.6 mg, 0.21 mmol.) and 2-bromo-1-(2,3-dihydro-1,4-benzodioxin-6-yl)ethan-1-one (50 mg, 0.20 mmol.). The desired product was isolated as an orange-yellow glassy solid (8.2 mg, 13%). ^1^H NMR (400 MHz, CDCl_3_) δ 7.30 (m, 2H), 6.88(d, J = 8.28 Hz, 1H), 6.70 (bs, 1H), 4.29 (m, 4H), 3.37 (s, 3H); ^13^C NMR (100 MHz, CDCl_3_) δ 189.9, 165.3, 156.0, 150.3, 143.5,138.3, 128.5, 123.1, 112.7, 112.3, 65.5, 65.2, 84.1, 57.0; MS (ESI): *m/z* 322.0 [(M+H)^+^].

**(4-amino-2-(phenylamino)thiazol-5-yl)(2,3-dihydrobenzo[b][1,4]dioxin-6-yl)methanone (12k).** General Method B was followed using pheny isothiocyanate (39 µL, 0.21 mmol.), potassium tert-butoxide (50.5 mg, 0.45 mmol.), cyanamide (8.6 mg, 0.21 mmol.) and 2-bromo-1-(2,3-dihydro-1,4-benzodioxin-6-yl)ethan-1-one (50 mg, 0.20 mmol.). The desired product was isolated as yellow solid (24 mg, 36%). ^1^H NMR (400 MHz, CD_3_OD) δ 7.61 (m, 2H), 7.37 (m, 2H), 7.24 (m, 2H), 7.13 (m, 1H), 6.90 (m, 1H), 4.30 (m, 4H); ^13^C NMR (100 MHz, CDCl_3_) δ 190.0, 162.2, 155.7, 150.0, 143.1,140.7, 138.0, 129.9, 128.2, 123.5, 122.8, 118.0, 112.8, 112.4, 65.6, 65.4; MS (ESI): *m/z* 354.1 [(M+H)^+^].

**(4-Amino-2-(benzylamino)thiazol-5-yl)(2,3-dihydrobenzo[b][1,4]dioxin-6-yl)methanone (12l)** General Method B was followed using benzyl isothiocyanate (27.1 µL, 0.21 mmol.), potassium tert-butoxide (50.5 mg, 0.4504 mmol.) cyanamide (8.6 mg, 0.21 mmol.) and 2-bromo-1-(2,3-dihydro-1,4-benzodioxin-6-yl)ethan-1-one (50 mg, 0.20 mmol.). The desired product was isolated as an orange-yellow glassy solid (12 mg, 17%). ^1^H NMR (400 MHz, CD_3_OD) δ 7.34 (m, 4H), 7.28 (m, 1H), 7.18 (m, 2H), 6.86 (m, 1H), 4.55 (m, 2H), 4.27 (m, 4H); ^13^C NMR (100 MHz, CDCl_3_) δ 190.1, 163.2, 155.5, 149.9, 143.2, 140.8, 137.9, 128.7, 128.1, 126.9, 126.5, 122.2, 112.4, 112.2, 65.4, 65.3, 50.5; MS (ESI): *m/z* 368.1 [(M+H)^+^].

**(4-Amino-2-((3-methoxyphenyl)amino)thiazol-5-yl)(2,3-dihydrobenzo[b][1,4]dioxin-6-yl)methanone (12m).** General Method B was followed using 3-methoxyphenylisothiocyanate (29.4 µL, 0.21 mmol.), potassium tert-butoxide (50.5 mg, 0.4504 mmol.), cyanamide (8.6 mg, 0.21 mmol.) and 2-bromo-1-(2,3-dihydro-1,4-benzodioxin-6-yl)ethan-1-one (50 mg, 0.20 mmol.). The desired product was isolated as a yellow solid (13 mg, 17%). ^1^H NMR (400 MHz, DMSO-d_6_) δ 10.72 (s, 1H), 8.17 (bs, 2H), 7.34 (m, 1H), 7.26 (t, J = 8.12 Hz, 1H), 7.21 (dd, J = 2.04 Hz, J = 8.24 Hz, 1H), 7.17 (d, J = 2.04 Hz, 1H), 7.09 (m, 1H), 6.93 (d, J = 8.32 Hz, 1H), 6.66 (dd, J = 2.08 Hz, J = 8.12 Hz, 1H), 4.29 (s, 4H), 3.76 (s, 3H); ^13^C NMR (100 MHz, CDCl_3_) δ 190.1, 162.0, 160.2, 155.7, 150.4, 144.1, 142.9, 138.0, 131.3, 122.8, 127.9, 112.4, 112.1, 111.6, 111.3, 99.9, 65.5, 65.3. 56.4; MS (ESI): *m/z* 384.1 [(M+H)^+^].

**(4-Amino-2-((3,4-dimethoxyphenyl)amino)thiazol-5-yl)(2,3-dihydrobenzo[b][1,4]dioxin-6-yl)methanone (12n).** General method B was followed using 3,4-dimethoxyphenyl isothiocyanate (31.3 µL, 0.21 mmol.), potassium tert-butoxide (50.5 mg, 0.4504 mmol.), cyanamide (8.6 mg, 0.21 mmol.) and 2-bromo-1-(2,3-dihydro-1,4-benzodioxin-6-yl)ethan-1-one (50 mg, 0.20 mmol.). The desired product was isolated as an orange solid (20.7 mg, 26%). ^1^H NMR (400 MHz, CD_3_OD) δ 7.27 (bd, J = 2.04 Hz, 1H), 7.20 (m, 2H), 7.04 (dd, J = 2.48 Hz, J = 8.64 Hz, 1H), 6.93 (d, J = 8.68 Hz, 1H), 6.07 (m, 1H), 4.27 (m, 4H), 3.85 (s, 3H), 3.82 (s, 3H); ^13^C NMR (100 MHz, CDCl_3_) δ 190.0, 160.3, 155.4, 150.4, 150.2, 143.0, 140.5, 138.1, 135.9, 128.1, 122.5, 114.6, 113.7, 112.6, 112.2, 106.8, 65.5, 65.4, 56.8, 56.7; MS (ESI): *m/z* 414.1 [(M+H)^+^].

**(4-Amino-2-((3,5-dimethoxyphenyl)amino)thiazol-5-yl)(2,3-dihydrobenzo[b][1,4]dioxin-6-yl)methanone (12o)**. General Method B was followed using 3,5-dimethoxyphenylisothiocyanate (41 mg, 0.21 mmol.), potassium tert-butoxide (50.5 mg, 0.4504 mmol.), cyanamide (8.6 mg, 0.21 mmol.) and 2- bromo-1-(2,3-dihydro-1,4-benzodioxin-6-yl)ethan-1-one (50 mg, 0.20 mmol.). The desired product was isolated as a yellow solid (5.6 mg, 7%). ^1^H NMR (400 MHz, DMSO-d_6_) δ 10.68 (s, 1H), 8.15 (bs, 2H), 7.20 (m, 2H), 6.94 (d, J = 8.28 Hz, 1H), 6.84 (d, J = 1.92 Hz, 1H), 6.25 (t, J = 2.04 Hz, 1H), 4.29 (s, 4H), 3.75 (s, 6H); ^13^C NMR (100 MHz, CDCl_3_) δ 189.8, 161.3, 160.7, 155.6, 149.6, 145.0, 142.7, 138.0, 128.0, 122.5, 112.8, 112.4, 92.3, 91.1, 65.5, 65.3, 56.4; MS (ESI): *m/z* 414.1 [(M+H)^+^].]

**(4-Amino-2-(benzo[d][1,3]dioxol-5-ylamino)thiazol-5-yl)(2,3-dihydrobenzo[b][1,4]dioxin-6-yl)methanone (12p)**. General Method B was followed using 3,4-methylenedioxyphenyl isothiocyanate (37.6 mg, 0.21 mmol.), potassium tert-butoxide (50.5 mg, 0.4504 mmol.), cyanamide (8.6 mg, 0.21 mmol.) and 2-bromo-1-(2,3-dihydro-1,4-benzodioxin-6-yl)ethan-1-one (50 mg, 0.20 mmol.). The desired product was isolated as a yellow solid (32 mg, 41%). ^1^H NMR (400 MHz, DMSO-d_6_) δ 10.59 (s, 1H), 8.14 (bs, 2H), 7.34 (bs, 1H), 7.18 (dd, J = 2.12 Hz, J = 8.32 Hz, 1H), 7.15 (d, J = 2.04 Hz, 1H), 6.92 (m, 3H), 6.02 (s, 2H), 4.28 (d, J = 2.84 Hz, 4H); ^13^C NMR (100 MHz, CDCl_3_) δ 190.1, 180.0, 155.2, 149.8, 148.3, 142.9, 139.1, 138.0, 136.1, 127.9, 122.5, 114.7, 112.5, 112.1, 109.1, 106.6, 101.8, 65.4, 65.2; MS (ESI): *m/z* 398.1 [(M+H)^+^].

**(4-Amino-2-((4-methoxybenzyl)amino)thiazol-5-yl)(2,3-dihydrobenzo[b][1,4]dioxin-6-yl)methanone (12q)** General Method B was followed using 4-methoxybenzyl isothiocynate (34.6 µL, 0.21 mmol.), potassium tert-butoxide (50.5 mg, 0.4504 mmol.), cyanamide (8.6 mg, 0.21 mmol.) and 2-bromo-1-(2,3-dihydro-1,4-benzodioxin-6-yl)ethan-1-one (50 mg, 0.20 mmol.). The desired product was isolated as an orange-yellow glassy solid (17.3 mg, 22%). ^1^H NMR (400 MHz, CDCl_3_) δ 7.23-7.32 (m, 4H), 6.88 (m, 3H), 6.35 (bs, 1H), 4.39 (m, 2H), 4.28 (m, 4H), 3.80 (s, 3H); ^13^C NMR (100 MHz, CDCl_3_) δ 190.1, 163.9, 159.4, 155.6, 150.0, 142.9, 138.1, 133.0, 131.1, 127.9, 122.5, 114.7, 112.7, 112.3, 65.6, 65.4, 56.3, 49.9; MS (ESI): *m/z* 398.1 [(M+H)^+^].

**(4-Amino-2-((4-ethoxyphenyl)amino)thiazol-5-yl)(2,3-dihydrobenzo[b][1,4]dioxin-6-yl)methanone (12r)**. General Method B was followed using 4-ethoxyphenylisothiocyanate (37.6 mg, 0.21 mmol.), potassium tert-butoxide (50.5 mg, 0.4504 mmol.), cyanamide (8.6 mg, 0.21 mmol.) and 2-bromo-1-(2,3-dihydro-1,4-benzodioxin-6-yl)ethan-1-one (50 mg, 0.20 mmol.). The desired product was isolated as an orange-yellow glassy solid (13.3 mg, 17%). ^1^H NMR (400 MHz, CDCl_3_) δ 7.26 (m, 4H), 6.90 (m, 2H), 6.85 (d, J = 8.28 Hz, 1H), 4.27 (m, 4H), 4.03 (q, J = 7.00 Hz, 2H), 1.42 (t, J = 6.96 Hz, 3H); ^13^C NMR (100 MHz, CDCl_3_) δ 189.9, 164.1, 158.2, 155.4, 150.2, 142.9, 138.0, 132.1, 130.7, 127.5, 122.5, 114.8,112.7, 112.3, 65.7, 65.6, 65.4; 49.9, 16.7; MS (ESI): *m/z* 398.1 [(M+H)^+^].

**(4-Amino-2-(phenethylamino)thiazol-5-yl)(2,3-dihydrobenzo[b][1,4]dioxin-6-yl)methanone (12s)** General Method B was followed using phenylethyl isothiocynate (30.5 µL, 0.21 mmol.), potassium tert-butoxide (50.5 mg, 0.4504 mmol.), cyanamide (8.6 mg, 0.21 mmol.) and 2-bromo-1-(2,3-dihydro-1,4-benzodioxin-6-yl)ethan-1-one (50 mg, 0.20 mmol.). The desired product was isolated as an orange-yellow glassy solid (17.1 mg, 23%). ^1^H NMR (400 MHz, CDCl_3_) δ 7.27-7.35 (m, 5H), 7.19 (m, 2H), 6.88 (d, J = 8.28 Hz, 1H), 5.72 (bs, 1H), 4.29 (m, 4H), 3.53 (m, 2H), 2.94 (t, J = 6.80 Hz, 2H); ^13^C NMR (100 MHz, CDCl_3_) δ 190.1, 163.9, 155.7, 150.1, 143.0, 140.2, 138.1, 129.5, 128.3, 128.0, 126.4, 122.9, 112.4, 112.1, 65.7, 65.5, 44.4, 35.8; MS (ESI: *m/z* 382.1 [(M+H)^+^].

**(4-Amino-2-(4-methoxyphenethylamino)thiazol-5-yl)(2,3-dihydrobenzo[b][1,4]dioxin-6-yl)methanone (12t)** General Method B was followed using 1-(2-isothiocyanatoethyl)-4-methoxybenzene (41.2 µL, 0.21 mmol.), potassium tert-butoxide (50.5 mg, 0.4504 mmol.), cyanamide (8.6 mg, 0.21 mmol.) and 2-bromo-1-(2,3-dihydro-1,4-benzodioxin-6-yl)ethan-1-one (50 mg, 0.20 mmol.). The desired product was isolated as an orange-yellow solid (24.3 mg, 30%). ^1^H NMR (400 MHz, CDCl_3_) δ 7.30 (m, 2H), 7.11 (M, 2H), 6.87 (M, 3H), 5.64 (bs, 1H), 4.29 (m, 4H), 3.80 (s, 3H), 3.49 (m, 2H), 2.88 (t, J = 6.80 Hz, 2H); ^13^C NMR (100 MHz, CDCl_3_) δ 190.2, 164.1, 158.6, 155.8, 150.4, 142.9, 138.1, 132.5, 130.8, 127.9, 122.7, 114.9, 112.8, 112.4, 65.6, 65.5, 56.2, 44.5, 35.9; MS (ESI): *m/z* 412.1 [(M+H)^+^].

**(4-Amino-2-((1-phenylethyl)amino)thiazol-5-yl)(2,3-dihydrobenzo[b][1,4]dioxin-6-yl)methanone (12u)** General Method B was followed using 1-phenylethyl isothiocyanate (36.7 µL, 0.21 mmol.), potassium tert-butoxide (50.5 mg, 0.4504 mmol.), cyanamide (8.6 mg, 0.21 mmol.) and 2-bromo-1-(2,3-dihydro-1,4-benzodioxin-6-yl)ethan-1-one (50 mg, 0.20 mmol.). The desired product was isolated as an orange-yellow solid (26.7 mg, 36%). ^1^H NMR (400 MHz, CDCl_3_) δ 7.22-7.37 (m, 7H), 6.85 (d, J = 8.32 Hz, 1H), 6.11 (m, 1H), 4.66 (m 1H), 4.27 (m, 4H), 1.60 (d, J = 6.76 Hz, 3H); ^13^C NMR (100 MHz, CDCl_3_) δ 189.9, 163.6, 155.4, 150.0, 144.1, 142.6, 137.9, 129.1, 128.3, 127.7, 127.3, 122.5, 112.6, 112.2, 65.2, 65.1, 60.7, 23.2; MS (ESI): *m/z* 382.1 [(M+H)^+^].

**(R)-(4-Amino-2-((1-(4-methoxyphenyl)ethyl)amino)thiazol-5-yl)(2,3-dihydrobenzo[b][1,4]dioxin-6-yl)methanone (12v)** General Method B was followed using (R)-1-(4-methoxyphenyl)ethyl isothiocyanate (37.7 µL, 0.21 mmol.), potassium tert-butoxide (50.5 mg, 0.4504 mmol.), cyanamide (8.6 mg, 0.21 mmol.) and 2-bromo-1-(2,3-dihydro-1,4-benzodioxin-6-yl)ethan-1-one (50 mg, 0.20 mmol.). The desired product was isolated as an orange-yellow glassy solid (29.5 mg, 37%). ^1^H NMR (400 MHz, CDCl_3_) δ 7.25 (m, 4H), 6.86 (m, 3H), 6.02 (M, 1H), 4.60 (m, 1H), 4.28 (m, 4H), 3.80 (s, 3H), 1.57 (d, J = 6.76 Hz, 3H); ^13^C NMR (100 MHz, CDCl_3_) δ 189.8, 163.9, 159.8, 155.7, 149.9, 143.0, 137.9, 136.6, 128.9, 127.3, 122.5, 114.8, 112.8, 112.5, 65.5, 65.3, 60.7, 56.4, 23.0; MS (ESI): *m/z* 412.1 [(M+H)^+^].

**(S)-(4-Amino-2-((2,3-dihydro-1H-inden-1-yl)amino)thiazol-5-yl)(2,3-dihydrobenzo[b][1,4]dioxin-6-yl)methanone (12w)** General Method B was followed using (S)-1-indanyl isothiocyanate (30.9 µL, 0.21 mmol.), potassium tert-butoxide (50.5 mg, 0.4504 mmol.), cyanamide (8.6 mg, 0.21 mmol.) and 2- bromo-1-(2,3-dihydro-1,4-benzodioxin-6-yl)ethan-1-one (50 mg, 0.20 mmol.). The desired product was isolated as an orange-yellow glassy solid (37.8 mg, 49%). ^1^H NMR (400 MHz, CDCl_3_) δ 7.39 (bd, J = 7.32 Hz, 1H), 7.24-7.32 (m, 5H), 6.87 (d, J = 8.32 Hz, 1H), 5.01 (m, 1H), 4.28 (m, 4H), 3.03 (M, 1H), 2.89 (m, 1H), 2.69 (m, 1H), 1.93 (m, 1H); ^13^C NMR (100 MHz, CDCl_3_) δ 190.1, 163.9, 155.7, 150.0, 144.0, 142.9, 140.5, 138.1, 127.8, 126.4, 126.2, 125.1, 125.0, 122.5, 112.8, 112.5, 69.1, 65.8, 65.5, 34.6, 30.8; MS (ESI): *m/z* 394.1 [(M+H)^+^].

**(4-Amino-2-((6-methoxypyridin-3-yl)amino)thiazol-5-yl)(2,3-dihydrobenzo[b][1,4]dioxin-6-yl)methanone (12z)**. General Method B was followed using 5-isothiocyanato-2-methoxypyridine (34.9 mg, 0.21 mmol.), potassium tert-butoxide (50.5 mg, 0.4504 mmol.), cyanamide (8.6 mg, 0.21 mmol.) and 2-bromo-1-(2,3-dihydro-1,4-benzodioxin-6-yl)ethan-1-one (50 mg, 0.20 mmol.). The desired product was isolated as an orange-yellow solid (19.3 mg, 26%). ^1^H NMR (400 MHz, CD_3_OD) δ 8.36 (d, J = 2.40 Hz, 1H), 7.98 (dd, J = 2.80 Hz, J = 8.92 Hz, 1H), 7.20 (m, 2H), 6.88 (m, 1H), 6.81 (d, J = 8.84 Hz, 1H), 4.28 (m, 4H), 3.90 (s, 3H); ^13^C NMR (100 MHz, CDCl_3_) δ 189.8, 160.3, 158.3, 155.5, 150.1, 143.1, 138.1, 134.9, 132.7, 128.0. 122.9, 122.4, 112.5, 112.1, 112.0, 65.8, 65.6, 54.7; MS (ESI): *m/z* 385.1 [(M+H)^+^].

**Method C**. The synthesis of compound **12e** exemplifies General Method C.

**(4-Amino-2-(cyclopropylamino)thiazol-5-yl)(2,3-dihydrobenzo[b][1,4]dioxin-6-yl)methanone (12e).** A solution of **11** (20 mg, 0.07 mmol.) and cyclopropylamine (90 uL, 1.30 mmol.) in ethanol (500 uL) was stirred at 100°C in a sealed glass pressure vessel overnight. The reaction was then cooled to room temperature and concentrated on a rotary evaporator. The residue was purified by chromatography on silica gel using a gradient of 0 to 100% ethyl acetate in hexanes to afford the desired compound as an orange-yellow glassy solid (14.4 mg, 65%). ^1^H NMR (400 MHz, CDCl_3_) δ 7.33 (m, 2H), 6.90 (d, J = 8.24 Hz, 1H), 6.30 (bs, 1H), 4.29 (m, 4H), 2.60 (m, 1H), 0.84 (m, 2H), 0.71 (m, 2H); ^13^C NMR (100 MHz, CDCl_3_) δ 190.1, 164.0, 156.0, 150.2, 143.1, 137.5, 127.8, 121.9, 112.8, 112.4, 65.9, 65.7, 26.8, 10.2; MS (ESI): *m/z* 657.2 [(2M+Na)^+^].

The following compounds were prepared using a similar procedure:

**(4-Amino-2-(isobutylamino)thiazol-5-yl)(2,3-dihydrobenzo[b][1,4]dioxin-6-yl)methanone (12g).** General method C was followed using **11** (20 mg, 0.07 mmol.) and isobutylamine (129 uL, 1.30 mmol.). The desired product was isolated as a yellow solid (15.6 mg, 72%). ^1^H NMR (400 MHz, CDCl_3_) δ 7.30 (m, 2H), 6.88 (d, J = 8.28 Hz, 1H), 5.69 (m, 1H), 4.29 (m, 4H), 3.07 (t, J = 6.40 Hz, 2H), 1.93 (m, 1H), 0.97 (d, J = 6.68 Hz, 6H); ^13^C NMR (100 MHz, CDCl_3_) δ 190.0, 164.0, 155.5, 149.9, 143.1, 138.3, 128.0, 122.4, 112.6, 112.2, 65.5, 65.3, 61.0, 30.1, 23.6; MS (ESI): *m/z* 689.3 [(2M+Na)^+^].

**(4-Amino-2-((furan-2-ylmethyl)amino)thiazol-5-yl)(2,3-dihydrobenzo[b][1,4]dioxin-6-yl)methanone (12aa).** General method C was followed using **11** 20 mg, (0.07 mmol.) and 2-aminomethylfuran (115 uL, 1.30 mmol.). The desired product was isolated as a reddish-tan solid (15.7 mg, 68%). ^1^H NMR (400 MHz, CDCl_3_) δ 7.39 (m, 1H), 7.30 (m, 2H), 6.88 (d, J = 8.32 Hz, 1H), 6.34 (m, 2H), 6.07 (m, 1H), 4.47 (m, 2H), 4.29 (m, 4H); ^13^C NMR (100 MHz, CDCl_3_) δ 190.1, 164.0, 155.7, 150.2, 143.9, 143.4, 143.3, 138.0, 128.1, 122.7, 112.4, 112.1, 111.1, 107.2, 65.7, 65.2, 40.6, ; MS (ESI): *m/z* 737.2 [(2M+Na)^+^].

**(4-Amino-2-((tetrahydro-2H-pyran-4-yl)amino)thiazol-5-yl)(2,3-dihydrobenzo[b][1,4]dioxin-6-yl)methanone (12ac).** General method C was followed using **11** 20 mg, (0.07 mmol.) and 4-aminopyran (135 uL, 1.30 mmol.). The desired product was isolated as an orange-yellow solid (5.1 mg, 36%). ^1^H NMR (400 MHz, CDCl_3_) δ 7.30 (m, 2H), 6.89 (d, J = 8.28 Hz, 1H), 5.43 (m, 1H), 4.29 (m, 4H), 3.99 (m, 2H), 3.62 (m, 1H), 3.49 (m, 2H), 2.06 (m, 2H), 1.58 (m, 2H); ^13^C NMR (100 MHz, CDCl_3_) δ 189.8, 163.9, 156.0, 150.3, 143.2, 138.2, 127.9, 122.4, 112.6, 112.2, 66.4, 65.7, 65.3, 55.7, 37.2; MS (ESI): *m/z* 362.1 [(M+H)^+^].

**(4-Amino-2-((3-methoxycyclopentyl)amino)thiazol-5-yl)(2,3-dihydrobenzo[b][1,4]dioxin-6-yl)methanone (12ad).** General method C was followed using **11** (20 mg, 0.07 mmol.) and 3-methoxycyclopentyl amine (157 uL, 1.30 mmol.). The desired product was isolated as an orange-yellow glassy solid (7.0 mg, 58%). ^1^H NMR (400 MHz, CDCl_3_) δ 7.31 (m, 2H), 6.89 (m,1H), 5.64 (m, 1H), 4.29 (m, 4H), 3.97 (m, 1H), 3.91 (m, 1H), 3.28 (d, J = 3.80 Hz, 3H), 2.26 (m, 1H), 1.48-1.99 (m, 5H); ^13^C NMR (100 MHz, CDCl_3_) δ 189.9, 164.7, 155.7, 150.1, 143.3, 138.2, 128.0, 122.8, 112.8, 112.5, 85.7, 65.8, 65.3, 58.1, 50.5, 45.9, 31.1, 30.4; MS (ESI): *m/z* 376.2 [(M+H)^+^].

**(4-amino-2-((1-methylpiperidin-4-yl)amino)thiazol-5-yl)(2,3-dihydrobenzo[b][1,4]dioxin-6-yl)methanone (12ae).** General method C was followed using **11** (20 mg, 0.07 mmol.) and 1-methyl-4-aminopiperidine (163 uL, 1.30 mmol.). The desired product was isolated as an orange-yellow glassy solid (6.2 mg, 20%). ^1^H NMR (400 MHz, CD_3_OD) δ 7.17 (m, 2H), 6.87 (m, 1H), 4.28 (m, 4H), 3.98 (m, 1H), 3.58 (m, 2H), 3.14 (m, 2H), 2.89 (s, 3H), 2.36 (m, 2H), 1.76 (m, 2H); ^13^C NMR (100 MHz, CDCl_3_) δ 190.0, 163.7, 154.9, 150.3, 142.9, 137.9, 127.8, 122.5, 112.5, 112.0, 65.3, 64.9, 56.6, 55.1, 47.4, 32.2; MS (ESI): *m/z* 771.3 [(2M+Na)^+^].

**(4-Amino-2-((1-(methylsulfonyl)piperidin-4-yl)amino)thiazol-5-yl)(2,3-dihydrobenzo[b][1,4]dioxin-6-yl)methanone (12af).** General Method C was followed using **11** (20 mg, 0.07 mmol) and 4-amino-1-methanesulfonylpiperidine (232 mg, 1.30 mmol.). The desired product was isolated as a yellow solid (9.7 mg, 17%). ^1^H NMR (400 MHz, DMSO-d_6_) δ 8.65 (bs, 1H), 7.15 (m, 2H), 6.90 (d, J = 8.28 Hz, 1H), 4.28 (m, 4H), 3.50 (m, 2H), 3.32 (m, 1H), 2.88 (m, 5H), 2.02 (m, 2H), 1.54 (m, 2H); ^13^C NMR (100 MHz, CDCl_3_) δ 190.0, 155.4, 150.2, 164.7, 143.0, 137.9, 122.5, 127.9, 112.7, 112.3, 65.2, 65.1, 56.8, 45.2, 41.3, 29.7; MS (ESI): *m/z* 439.1 [(M+H)^+^].

**(4-Amino-2-(cycloheptylamino)thiazol-5-yl)(2,3-dihydrobenzo[b][1,4]dioxin-6-yl)methanone (12ag).** General method C was followed using **11** (30 mg, 0.07 mmol.) and cycloheptanamine (166 uL, 1.30 mmol.). The desired product was isolated as a yellow solid (15.4 mg, 64%). ^1^H NMR (400 MHz, CD_3_OD) δ 7.17 (m, 2H), 6.86 (m, 1H), 4.27 (m, 4H), 2.02 (m, 2H), 1.47-1.74 (m, 11H); ^13^C NMR (100 MHz, CDCl_3_) δ 190.1, 164.0, 155.6, 150.0, 142.9, 138.1, 128.1, 122.5, 112.4, 112.0, 65.2, 65.1, 51.4, 37.3, 27.1, 23.5; MS (ESI): *m/z* 374.1 [(M+H)^+^].

**(4-Amino-2-((cis-4-isopropylcyclohexyl)amino)thiazol-5-yl)(2,3-dihydrobenzo[b][1,4]dioxin-6-yl)methanone (12ah).** General method C was followed using **11** (30 mg, 0.07 mmol.) and rac-(1s,4s)-4-propan-2-yl)cyclohexan-1-amine, cis (216 uL, 1.30 mmol.). The desired product was isolated as a yellow solid (7.0 mg, 27%). ^1^H NMR (400 MHz, CD_3_OD) δ 7.18 (m, 2H), 6.86 (m, 1H), 4.27 (m, 4H), 1.88 (m, 2H), 1.57 (m, 5H), 1.42 (m, 2H), 1.09-1.31 (m, 2H), 0.91 (d, J = 6.76 Hz, 6H); ; ^13^C NMR (100 MHz, CDCl_3_) δ 190.0, 164.4, 155.4, 150.1, 143.1, 138.0, 128.3, 122.5, 112.7, 112.3, 65.7, 65.5, 55.3, 43.5, 32.9, 30.8, 26.6, 22.0; MS (ESI): *m/z* 402.1 [(M+H)^+^].

**(4-Amino-2-((trans-4-isopropylcyclohexyl)amino)thiazol-5-yl)(2,3-dihydrobenzo[b][1,4]dioxin-6-yl)methanone (12ai).** General method C was followed using **11** (30 mg, 0.07 mmol.) and rac-(1r,4r)-4-propan-2-yl)cyclohexan-1-amine, trans (216 uL, 1.30 mmol.). The desired product was isolated as a yellow solid (7.1 mg, 28%). ^1^H NMR (400 MHz, CD_3_OD) δ 7.18 (m, 2H), 6.86 (m, 1H), 4.27 (m, 4H), 2.09 (m, 2H), 1.81 (m, 2H), 1.46 (m, 1H), 1.26 (m, 3H), 1.11 (m, 3H), 0.91 (d, J = 6.84 Hz, 6H); ^13^C NMR (100 MHz, CDCl_3_) δ 190.2, 164.8, 155.7, 150.2, 143.5, 138.3, 128.4, 122.6, 112.7, 112.4, 65.8, 65.6, 56.5, 43.7, 33.1, 31.0, 26.9, 22.3; MS (ESI): *m/z* 402.1 [(M+H)^+^].

**(4-Amino-2-((3,5-dimethylcyclohexyl)amino)thiazol-5-yl)(2,3-dihydrobenzo[b][1,4]dioxin-6-yl)methanone (12aj).** General method C was followed using **11** (30 mg, 0.07 mmol.) and 3,5-dimethylcyclohexan-1-amine (200 uL, 1.30 mmol.). The desired product was isolated as a yellow solid (4.8 mg, 19%). ^1^H NMR (400 MHz, CDCl_3_) δ 7.29 (m, 1H), 7.26 (m, 1H), 6.90 (d, J = 8.24 Hz, 1H), 4.29 (m, 4H), 3.30 (m, 1H), 2.04 (m, 2H), 1.68 (m, 4H), 1.53 (m, 3H), 0.93 (d, J = 6.56 Hz, 6H); ^13^C NMR (100 MHz, CDCl_3_) δ 190.1, 164.3, 155.7, 150.3, 164.0, 142.8, 128.2, 122.6, 112.7, 112.3, 65.5, 65.3, 50.3, 44.7, 43.9, 29.0, 21.9; MS (ESI): *m/z* 388.2 [(M+H)^+^].

**(4-amino-2-((exo)-bicyclo[2.2.1]heptan-2-ylamino)thiazol-5-yl)(2,3-dihydrobenzo[b][1,4]dioxin-6-yl)methanone (12ak).** General method C was followed using **11** (20 mg, 0.07 mmol.) and exo-2-aminonorbornane (154 uL, 1.30 mmol.). The desired product was isolated as a yellow solid (18.2 mg, 75%). ^1^H NMR (400 MHz, CDCl_3_) δ 7.31 (m, 2H), 6.89 (d, J = 8.24 Hz, 1H), 5.52 (d, J = 7.32 Hz, 1H), 4.29 (m, 4H), 3.27 (m, 2H), 2.35 (m, 2H), 1.89 (m, 1H), 1.62-1.45 (m, 3H), 1.38 (m, 1H), 1.12-1.32 (m, 4H); ^13^C NMR (100 MHz, CDCl_3_) δ 190.2, 165.0, 155.7, 150.2, 143.7, 138.2, 128.4, 123.3, 112.8, 112.4, 65.6, 65.5, 59.3, 43.4, 40.0, 36.7, 36.1, 29.1, 27.4; MS (ESI): *m/z* 372.1 [(M+H)^+^].

**(2-(Adamantan-2-ylamino)-4-aminothiazol-5-yl)(2,3-dihydrobenzo[b][1,4]dioxin-6-yl)methanone (12al).** General method C was followed using **11** (20 mg, 0.07 mmol.) and 2-aminoadamantane (196 mg, 1.30 mmol.). The desired product was isolated as an off-white solid (2.6 mg, 10%). ^1^H NMR (400 MHz, CDCl_3_) δ 7.30 (m, 2H), 6.89 (d, J = 8.24 Hz, 1H), 5.97 (m, 1H), 4.29 (m, 4H), 3.55 (m, 1H), 2.08 (bs, 2H), 1.66-1.92 (m, 12H); ^13^C NMR (100 MHz, CDCl_3_) δ 190.2, 164.5, 155.5, 149.9, 143.1, 138.2, 128.1, 122.6, 112.7, 112.5, 65.7, 65.3, 64.9, 42.3, 37.2, 36.4, 31.5; MS (ESI): *m/z* 412.1 [(M+H)^+^].

**(4-Amino-2-(((bicyclo[2.2.1]heptan-2-yl)methyl)amino)thiazol-5-yl)(2,3-dihydrobenzo[b][1,4]dioxin-6-yl)methanone (12am).** General method C was followed using **11** (20 mg, 0.07 mmol.) and bicycle[2.2.1]heptane-2-methanamine (163 mg, 1.03 mmol.). The desired product was isolated as a yellow solid (22.4 mg, 45%). ^1^H NMR (400 MHz, CD_3_OD) δ 7.18 (m, 2H), 6.86 (m, 1H), 4.28 (m, 4H), 3.27 (m, 1H), 2.17 (m, 3H), 1.77 (m, 1H), 1.57 (m, 2H), 1.39 (m, 3H), 1.16 (m, 2H), 0.72 (m, 1H); ^13^C NMR (100 MHz, CDCl_3_) δ 190.2, 155.4, 164.4, 150.0, 142.8, 137.9, 128.0, 122.4, 112.4, 112.1, 65.5, 65.3, 53.6, 38.9, 43.3, 40.2, 36.5, 35.0, 29.0, 27.2 ; MS (ESI): *m/z* 386.1 [(M+H)^+^].

**(4-Amino-2-(([3a,4,7,7a]-octahydro-1H-4,7-methanoinden-5-yl)amino)thiazol-5-yl)(2,3-dihydrobenzo[b][1,4]dioxin-6-yl)methanone (12an**). General method C was followed using 11 (20 mg, 0.07 mmol.) and tricyclo[5.2.1.0,3,6]decan-8-amine (196 mg, 1.30 mmol.). The desired product was isolated as a yellow solid (14 mg, 52%). ^1^H NMR (400 MHz, CDCl_3_) δ 7.29 (m, 2H), 6.89 (m, 1H), 4.28 (m, 4H), 3.65 (m, 1H), 2.35 (bd, J = 3.84 Hz, 1H), 2.15 (m, 2H), 2.02 (bd, J = 4.64 Hz,1H), 1.85 (m, 2H), 1.65 (m, 1H), 1.48 (m, 1H), 1.17 (m, 3H), 0.96 (m, 2H), 0.80 (m, 1H); ^13^C NMR (100 MHz, CDCl_3_) δ 190.0, 164.7, 155.6, 150.1, 143.3, 138.1, 128.1, 122.5, 112.7, 112.3, 65.6, 65.5, 57.0. 46.4, 45.9. 42.3. 40.2, 38.1, 32.7. 32.6, 32.0, 27.9; MS (ESI): *m/z* 412.1 [(M+H)^+^].

**Method D**. The synthesis of compound **12x** exemplifies General Method D.

**(4-Amino-2-(pyrrolidin-1-yl)thiazol-5-yl)(2,3-dihydrobenzo[b][1,4]dioxin-6-yl)methanone (12x)**. Pyrrolidine (16.7 µL, 0.20 mmol.) was added to a solution of dimethyl N-cyanodithioimidocarbonate (28.4 mg, 0.20 mmol.) in anhydrous dimethylformamate (1 mL). The reaction was stirred at 70°C for 1 hour. Sodium sulfide nonahydrate (47 mg, 0.20 mmol) was added and stirring at 70°C was continued for 1.5 hours. The reaction was cooled to 50°C and a solution of **10** (100 mg, 0.39 mmol) in anhydrous dimethylformamide (1 mL) was added dropwise. Stirring at 50°C was continued for 2 hours. Potassium carbonate (27 mg, 0.20 mmol.) was added to the reaction and stirring at 50°C was continued for an additional 1 hour. The orange reaction mixture was cooled to room temperature and concentrated on a rotary evaporator. The residue was partitioned between ethyl acetate and water. The aqueous phase was extracted with two additional portions of ethyl acetate. The combined organic phases were washed with water and brine, dried over anhydrous sodium sulfate and concentrated on a rotary evaporator. The residue was purified by chromatography on silica gel using a gradient of 0 to 100% ethyl acetate in hexanes to afford the desired product as an off-white solid (40.3 mg, 63%). ^1^H NMR (400 MHz, CDCl_3_) δ 7.32 (m, 2H), 6.88 (d, J = 8.28 Hz, 1H), 4.28 (m, 4H), 3.48 (bs, 4H), 2.05 (m, 4H); ^13^C NMR (100 MHz, CDCl_3_) δ 190.2, 156.3, 155.4, 150.4, 143.2, 138.4, 128.0, 122.7, 112.5, 112.1, 65.8, 65.4, 55.6, 26.9; MS (ESI): *m/z* 685.2 [(2M+Na)^+^].

The following compounds were prepared using a similar procedure:

**(4-Amino-2-(piperidin-1-yl)thiazol-5-yl)(2,3-dihydrobenzo[b][1,4]dioxin-6-yl)methanone (12y)**. General Method D was followed using piperidine (19.8 µL, 0.20 mmol.), dimethyl N-cyanodithioimidocarbonate (28.4 mg, 0.20 mmol.), Sodium sulfide nonahydrate (47 mg, 0.20 mmol) and **10** (100 mg, 0.39 mmol). The desired product was obtained as a yellow solid (21.2 mg, 32%). ^1^H NMR (400 MHz, CDCl_3_) δ 7.31 (m, 2H), 6.87 (d, J = 8.32 Hz, 1H), 4.29 (s, 4H), 3.52 (bs, 4H), 1.67 (bs, 6H); MS (ESI): *m/z* 346.1 [(M+H)^+^].

**(4-Amino-2-morpholinothiazol-5-yl)(2,3-dihydrobenzo[b][1,4]dioxin-6-yl)methanone (12ab)**. General Method D was followed using morpholine piperidine (17.5 µL, 0.20 mmol.), dimethyl N-cyanodithioimidocarbonate (28.4 mg, 0.20 mmol.), Sodium sulfide nonahydrate (47 mg, 0.20 mmol) and **10** (100 mg, 0.39 mmol). The desired product was obtained as a glassy yellow solid (25.6 mg, 38%). ^1^H NMR (400 MHz, CDCl_3_) δ 7.30 (m, 2H), 6.88 (d, J = 8.28 Hz, 1H), 4.28 (m, 4H), 3.77 (m, 4H), 3.55 (m, 4H); ^13^C NMR (100 MHz, CDCl_3_) δ 190.0, 156.7, 155.3, 150.4, 142.8, 137.9, 128.0, 122.5, 112.6, 112.2, 65.7, 65.5, 55.5, 26.4, 25.1; MS (ESI): *m/z* 717.2 [(2M+Na)^+^].

### Synthesis of Compounds 13a – l

**(4-Amino-2-(methylthio)thiazol-5-yl)(3,4,5-trimethoxyphenyl)methanone (13a).** A solution of 2-bromo-1-(3,4,5-trimethoxyphenyl)ethan-1-one (647 mg, 2.24 mmol.) [84] in anhydrous DMF (5 mL) was added dropwise to a second solution of cyanimidodithiocarbonic acid S-methyl ester S-potassium salt (381 mg, 2.24 mmol.) in anhydrous DMF (5 mL) at 50°C. This mixture was stirred at 50°C for two hours. Potassium carbonate (309 mg, 2.24 mmol,) was added and stirring was continued for one hour. The reaction was cooled to room temperature and concentrated on a rotary evaporator. The residue was partitioned between ethyl acetate and water. The aqueous layer was extracted with two additional portions of ethyl acetate. The combined organic layers were washed with water and brine, dried over anhydrous sodium sulfate and concentrated on a rotary evaporator. The residue was purified by chromatography on silica gel using a gradient of 0 to 100% of ethyl acetate in hexanes to afford the desired compound as an orange-yellow solid (329 mg, 43%).^1^H NMR (400 MHz, CDCl_3_) δ 7.05 (s, 2H), 3.93 (m, 9H), 2.69 (s, 3H); ESIMS: *m/z* 363.0 [(M+Na)^+^].

**(4-Amino-2-(methylthio)thiazol-5-yl)(benzo[d][1,3]dioxol-5-yl)methanone (13b).** Prepared using the same procedure as described for the synthesis of compound **19** using 1-(1,3-benzodioxol-5-yl)-2-bromoethan-1-one (244 mg, 1.01 mmol.), triethylamine (180 uL, 1.30 mmol.) and cyanimidodithiocarbonic acid S-methyl ester S-potassium salt (154 mg, 0.90 mmol.). The desired product was obtained as an orange solid (208 mg, 78%). ^1^H NMR (400 MHz, CDCl_3_) δ 7.36 (dd, J = 8.04 Hz, J = 1.72 Hz, 1H), 7.27 (d, J = 1.68 Hz, 1H), 6.84 (d, J = 8.08 Hz, 1H), 6.04 (s, 2H), 2.66 (s, 3H); MS (ESI): *m/z* 295.0 [(M+H)^+^].

**(4-Amino-2-(methylthio)thiazol-5-yl)(3,4-dihydro-2H-benzo[b][1,4]dioxepin-7-yl)methanone (13c)**. Prepared using the same procedure as described for the synthesis of compound **10** using 7-bromoacetyl-3,4-dihydro-1,5-benzodioxepin (272 mg, 1.01 mmol.), triethylamine (180 uL, 1.30 mmol.) and cyanimidodithiocarbonic acid S-methyl ester S-potassium salt (154 mg, 0.90 mmol.). The desired product was obtained as an orange solid (212 mg, 73%). ^1^H NMR (400 MHz, CDCl_3_) δ 7.39 (m, 2H), 6.99 (m, 1H), 4.28 (m, 4H), 2.65 (s, 3H), 2.22 (m, 2H); MS (ESI): *m/z* 323.0 [(M+H)^+^].

**<u>(4-Amino-2-(methylthio)thiazol-5-yl)(2-nitrophenyl)methanone</u> (13d)**. Prepared using the same procedure as described for the synthesis of compound **10** using 2-bromo-2’-nitroacetophenone (245 mg, 1.00 mmol.), triethylamine (180 uL, 1.31 mmol.) and cyanimidodithiocarbonic acid S-methyl ester S-potassium salt (154 mg, 0.90 mmol,). The desired product was obtained as a yellow solid (182 mg, 68%). ^1^H NMR (400 MHz, DMSO) δ 8.13 (dd, J = 8.08 Hz, J = 0.92 Hz, 1H), 7.97 (bs, 2H), 7.84 (td, J = 7.48 Hz, J = 1.08 Hz, 1H), 7.74 (td J = 8.08 Hz, J = 1.44 Hz, 1H), 7.68 (dd, J = 7.48 Hz, J = 1.36 Hz, 1H), 2.62 (s, 3H); MS (ESI): *m/z* 296.0 [(M+H)^+^].

**(4-Amino-2-(methylthio)thiazol-5-yl)(pyridin-3-yl)methanone (13e)**. Prepared using the same procedure as described for the synthesis of compound **10** using 3-(bromoacetyl)pyridine HBr (282 mg, 1.00 mmol.), triethylamine (348 uL, 2.51 mmol.) and cyanimidodithiocarbonic acid S-methyl ester S-potassium salt (154 mg, 0.90 mmol,). The desired product was obtained as a bright yellow solid (99 mg, 65%). ^1^H NMR (400 MHz, CDCl_3_) δ 9.01 (d, J = 1.68 Hz, 1H), 8.73 (dd, J = 1.64 Hz, J = 4.84 Hz, 1H), 8.04 (dt, J = 7.88 Hz, J = 1.96 Hz, 1H), 7.39 (m, 1H), 2.67 (s, 3H); MS (ESI): *m/z* 252.0 [(M+H)^+^].

**(4-Amino-2-(methylthio)thiazol-5-yl)(3-nitrophenyl)methanone (13f)**. Prepared using the same procedure as described for the synthesis of compound **10** using 2-bromo-3’-nitroacetophenone (245 mg, 1.00 mmol.), triethylamine (180 uL, 1.31 mmol.) and cyanimidodithiocarbonic acid S-methyl ester S-potassium salt (154 mg, 0.90 mmol,). The desired product was obtained as a yellow solid (123 mg, 46%). ^1^H NMR (400 MHz, DMSO) δ 8.38 (m, 2H), 8.23 (bs, 2H), 8.14 (m, 1H), 7.80 (t, J = 7.96 Hz, 1H), 2.68 (s, 3H); MS (ESI): *m/z* 295.9 [(M+H)^+^].

**(4-Amino-2-(methylthio)thiazol-5-yl)(4-nitrophenyl)methanone (13g)**. Prepared using the same procedure as described for the synthesis of compound **10** using 2-bromo-4’-nitroacetophenone (245 mg, 1.00 mmol.), triethylamine (180 uL, 1.31 mmol.) and cyanimidodithiocarbonic acid S-methyl ester S-potassium salt (154 mg, 0.90 mmol,). The desired product was obtained as a reddish-orange solid (100 mg, 38%). ^1^H NMR (400 MHz, DMSO) δ 8.33 (m, 2H), 8.25 (bs, 2H), 7.93 (m, 2H), 2.68 (s, 3H); MS (ESI): *m/z* 296.7 [(M+H)^+^].

**2-(4-Amino-2-(methylthio)thiazole-5-carbonyl)benzonitrile (13h)**. Prepared using the same procedure as described for the synthesis of compound **10** using 2-(2-bromoacetyl)benzonitrile (140 mg, 0.63 mmol.), triethylamine (113 uL, 0.81 mmol.) and cyanimidodithiocarbonic acid S-methyl ester S-potassium salt (196 mg, 0.56 mmol,). The desired product was obtained as an orange-yellow solid (49 mg, 32%). ^1^H NMR (400 MHz, CDCl_3_) δ 7.79 (d, J = 7.64 Hz, 1H), 7.67 (m, 2H), 7.56 (td, J = 7.52 Hz, J = 1.48 Hz, 1H), 2.65 (s, 3H); MS (ESI): *m/z* 275.7 [(M+H)^+^].

**(4-Amino-2-(methylthio)thiazol-5-yl)(7-nitro-2,3-dihydrobenzo[b][1,4]dioxin-6-yl)methanone (13i)** *Step 1. 2-Bromo-1-(7-nitro-2,3-dihydrobenzo[b][1,4]dioxin-6-yl)ethan-1-one*. A solution of bromine (138 uL, 2.67 mmol.) in anhydrous dioxane (3 mL) was added dropwise to a solution of 1-(7-nitro-2,3-dihydrobenzo[b][1,4]dioxin-6-yl)ethan-1-one [85] (590 mg, 2.64 mmol.) in anhydrous dioxane (5mL). The reaction was stirred at room temperature for 4 hours and then concentrated on a rotary evaporator. The residue was dissolved in ethyl acetate, washed with saturated aqueous sodium bicarbonate and brine, dried over anhydrous sodium sulfate and concentrated on a rotary evaporator. The residue was purified by column chromatography using dichloromethane. The desired product was obtained as a yellow solid (650 mg, 81%). ^1^H NMR (400 MHz, CDCl_3_) δ 7.75 (s, 1H), 6.93 (s, 1H), 4.37(m, 4 H), 4.23 (s, 2H); MS (ESI): *m/z* 301.9 [(M+H)^+^].

*Step 2. (4-Amino-2-(methylthio)thiazol-5-yl)(7-nitro-2,3-dihydrobenzo[b][1,4]dioxin-6-yl)methanone (**13i**).* Prepared using the same procedure as described for the synthesis of compound **10** using 2- bromo-1-(7-nitro-2,3-dihydrobenzo[b][1,4]dioxin-6-yl)ethan-1-one (190 mg, 0.63 mmol.), triethylamine (113 uL, 0.81 mmol.) and cyanimidodithiocarbonic acid S-methyl ester S-potassium salt (96 mg, 0.56 mmol,). The desired product was obtained as a yellow solid (101 mg, 51%). ^1^H NMR (400 MHz, DMSO-d_6_) δ 7.88 (bs, 2H), 7.68 (s, 1H), 7.13 (s, 1H), 4.38 (m, 4H), 2.61 (s, 3H); MS (ESI): *m/z* 354.0 [(M+H)^+^].

**(4-Amino-2-(methylthio)thiazol-5-yl)(6-nitrobenzo[d][1,3]dioxol-5-yl)methanone (13j)**. Prepared using the same procedure as described for the synthesis of compound **10** using 2-bromo-1-(6-nitrobenzo[d][1,3]dioxol-5-yl)ethan-1-one [86] (200 mg, 0.69 mmol.), triethylamine (125 uL, 0.90 mmol.) and cyanimidodithiocarbonic acid S-methyl ester S-potassium salt (106 mg, 0.62 mmol,). The desired product was obtained as an orange-brown solid (178 mg, 84%). ^1^H NMR (400 MHz, CDCl_3_) δ 7.60 (s, 1H), 6.85 (s, 1H), 6.19 (s, 2H), 2.60 (s, 3H); MS (ESI): *m/z* 339.9 [(M+H)^+^].

**(4-Amino-2-(methylthio)thiazol-5-yl)(benzo[c][1,2,5]oxadiazol-4-yl)methanone (13k)*. Step 1. 1,3-****Dibromo-2-nitrosobenzene*. [87] 3-Chloroperoxybenzoic acid (77%, 2.65 g, 11.97 mmol.) was added to a solution of 2,6-dibromoaniline (1.00 g, 3.39 mmol) in chloroform (24 mL). The resulting thick light green suspension was stirred at room temperature overnight. The reaction mixture was diluted with chloroform, washed twice with saturated aqueous sodium thiosulfate, washed three times with saturated aqueous sodium bicarbonate, washed with brine, dried over anhydrous sodium sulfate and concentrated on a rotary evaporator to afford the desired compound as a light tan solid (976 mg, 92%). The product was taken on to the next step without further purification. ^1^H-NMR (400 MHz, CDCl_3_) δ 7.74 (d, J = 7.96 Hz, 2H), 7.25 (m, 1H).

*Step 2. 4-Bromobenzo[c][1,2,5’oxadiazole*. Sodium azide (263 mg, 4.05 mmol.) was added to a solution of 1,3-dibromo-2-nitrosobenzene (976 mg, 3.68 mmol.) in anhydrous dimethylsulfoxide (30 mL). The solution was stirred at room temperature under a nitrogen atmosphere for 2 hours and then heated at 120°C for 10 minutes. The reaction was cooled to room temperature and poured over ice. The resulting precipitate was filtered, washed with water and dissolved in dichloromethane. The solution was dried over anhydrous sodium sulfate and concentrated on a rotary evaporator to afford the desired compound as an orange-tan solid (573 mg, 73%). The product was taken on to the next step without further purification. ^1^H-NMR (400 MHz, CDCl_3_) δ 7.82 (d, J = 9.00 Hz, 1H), 7.64 (d, J = 7.64 Hz, 1H), 7.30 (m, 1H).

*Step 3. 1-(benzo[c][1,2,5]oxadiazol-4-yl)ethan-1-one*. A solution of 4-bromobenzo[c][1,2,5]oxadiazole (100 mg, 0.50 mmol.) in anhydrous toluene (2 mL) was degassed and placed under a nitrogen atmosphere. 1-Ethoxyvinyl tributyl tin (20.2 µL, 0.06 mmol.) and bis(triphenylphosphine)palladium (II) dichloride (39 mg, 0.06 mmol.) were added sequentially. The reaction was stirred at 90°C overnight. The resulting dark reaction mixture was cooled to room temperature and filtered through a plug of Celite to yield an orange filtrate. 6N Aqueous HCl (2 mL) was added and the resulting biphasic mixture was vigorously stirred at room temperature for one hour. The mixture was concentrated to dryness on a rotary evaporator and the resulting residue was partitioned between dichloromethane and saturated aqueous sodium bicarbonate. The organic layer was dried over anhydrous sodium sulfate and concentrated on a rotary evaporator. The residue was purified by chromatography on silica gel using a gradient of 0 to 40% ethyl acetate in hexanes to afford the desired compound as a pale yellow solid (42 mg, 52%). ^1^H-NMR (400 MHz, CDCl_3_) δ 8.12 (m, 2H), 7.57 (m, 1H), 2.94 (s, 2H).

*Step 4. 1-(Benzo[c][1,2,5]oxadeiazol-4-yl)-2-bromoethan-1-one*. 1-(benzo[c][1,2,5]oxadiazol-4-yl)ethan-1-one (216 mg, 1.33 mmol.), ammonium acetate (10.3 mg, 0.13 mmol.) and N-bromosuccinimide (296 mg, 1.67 mmol.) were combined in anhydrous diethyl ether (6 mL) and stirred at room temperature for one hour. Carbon tetrachloride (6 mL) wad added and the reaction was heated to 80°C. Approximately half of the diethyl ether was allowed to boil off and the remaining reaction was refluxed at 80°C. Extra N-bromosuccinimide was added in portions until the starting material was consumed by thin layer chromatography. The reaction as cooled to room temperature and the resulting precipitate removed by filtration. The filtrate was concentrated on a rotary evaporator. The residue was dissolved in dichloromethane, washed with water, dried over anhydrous sodium sulfate and concentrated on a rotary evaporator. The resulting residue was purified by chromatography on silica gel using a gradient of 0 to 30% ethyl acetate in hexanes to afford the desired compound as an arrange-yellow solid (198 mg, 62%). ^1^H-NMR (400 MHz, CDCl_3_) δ 8.28 (dd, J = 6.80 Hz, J = 0.76 Hz, 1H), 8.16 (dd, J = 9.00 Hz, J = 0.76 Hz, 1H), 7.62 (m, 1H), 4.90 (s, 2H).

*Step 5. (4-Amino-2-(methylthio)thiazol-5-yl)(benzo[c][1,2,5]oxadiazol-4-yl)methanone (**13k**)*. Prepared using the same procedure as described for the synthesis of compound **10** using 1-(benzo[c][1,2,5]oxa-deiazol-4-yl)-2-bromoethan-1-one (139 mg, 0.91 mmol.), triethylamine (162 uL, 1.17 mmol) and cyanimido-dithiocarbonic acid S-methyl ester S-potassium salt (137 mg, 0.81 mmol). Product was obtained as a black solid (128 mg, 54%). ^1^H-NMR (400 MHz, CD_3_OD) δ 8.24 (bs, 2H), 8.22 (dd, J = 9.12 Hz, J = 0.64 Hz, 1H), 7.90 (dd, J = 6.60 Hz, J = 0.64 Hz, 1H), 7.71 (m, 1H), 2.65 (s, 3H); MS (ESI) *m/z* 293.0 [(M+H)^+^].

**(4-Amino-2-(methylthio)thiazol-5-yl)(benzo[c][1,2,5]oxadiazol-5-yl) methanone (13l).** *Step 1. 1-(Benzo[c][1,2,5]oxadiazol-5-yl)ethan-1-one.* A solution of 5-bromobenzo[c][1,2,5]oxadiazole [88] (1.00 g, 5.03 mmol.) in anhydrous toluene (20 mL) was degassed under vacuum for 10 minutes. The solution was then blanketed with nitrogen gas. 1-Ethoxy vinyl tributyltin (1.87 mL, 5.53 mmol) and bis(triphenylphosphine)palladium(II) dichloride (388 mg, 0.55 mmol) were added sequentially. The resulting mixture was stirred under nitrogen at 90°C overnight. After cooling to room temperature, the dark mixture was filtered through a plug of Celite. The resulting orange filtrate was treated with 6N aqueous hydrochloric acid (20 mL) and the mixture was vigorously stirred for one hour. The toluene was removed on a rotary evaporator. The remaining aqueous layer was neutralized to pH 8 with saturated aqueous sodium bicarbonate solution and extracted with dichloromethane. The dichloromethane extract was dried over anhydrous sodium sulfate and concentrated on a rotary evaporator. The crude product was purified by column chromatography on silica gel using a gradient of 0 to 40% ethyl acetate in hexanes to afford the desired product as a pale yellow solid (604 mg, 74%). ^1^H-NMR (400 MHz, CDCl_3_) δ 8.47 (t, J = 1.20 Hz, 1H), 8.02 (dd, J = 1.40 Hz, J = 9.48 Hz, 1H), 7.90 (dd, J = 1.04 Hz, J = 9.44 Hz, 1H), 2.72 (s, 3H); MS (ESI) *m/z* 162.9 [(M+H)^+^].

*Step 2. 1-(Benzo[c][1,2,5]oxadiazol-5-yl)-2-bromoethan-1-one*. A solution of 1-(benzo[c][1,2,5]oxadiazol-5-yl)ethan-1-one (600 mg, 3.70 mmol.), ammonium acetate (29 mg, 0.37 mmol.), and N-bromosuccinimide (823 mg, 4.63 mmol.) in anhydrous diethyl ether (20 mL) was stirred at room temperature for one hour. Carbon tetrachloride (20 mL) was added and the reaction was heated at 80°C to remove half of the diethyl ether. The remaining solution was refluxed at 80°C for ten minutes. An additional portion of N-bromosuccinimide (600 mg) was added portionwise at reflux until the reaction went to completion (as determined by LC/MS). The reaction was cooled to room temperature and the resulting precipitate was removed by filtration. The filtrate was concentrated on a rotary evaporator. The residue was dissolved in dichloromethane and the organic solution was washed with water, dried over anhydrous sodium sulfate and concentrated on a rotary evaporator. The crude product was purified by column chromatography on silica gel using a gradient of 0 to 30% ethyl acetate in hexanes to afford the desired product as a pale orange crystalline solid (636 mg, 71%). ^1^H-NMR (400 MHz, CDCl_3_) δ 8.57 (t, J = 1.24 Hz, 1H), 7.98 (m, 2H), 4.49 (s, 2H); MS (ESI) *m/z* 240.7 [(M+H)^+^].

*Step 3. (4-Amino-2-(methylthio)thiazol-5-yl)(benzo[c][1,2,5]oxadiazol-5-yl)methanone (**13l**)*. Prepared using the same procedure as described for the synthesis of compound **10** using 1- (benzo[c][1,2,5]oxadiazol-5-yl)-2-bromoethan-1-one (242 mg, 1.01 mmol.), triethylamine (180 uL, 1.30 mmol.) and cyanimidodithiocarbonic acid S-methyl ester S-potassium salt (154 mg, 0.90 mmol,). Product was obtained as a black solid (59 mg, 22%). ^1^H-NMR (400 MHz, DMSO-d_6_) δ 8.41 (t, J = 1.16 Hz, 1H), 8.23 (bs, 2H), 8.17 (dd, J = 1.04 Hz, J = 9.27 Hz, 1H), 7.76 (dd, J = 1.28 Hz, J = 9.28 Hz, 1H), 2.68 (s, 3H); MS (ESI) *m/z* 292.7 [(M+H)^+^].

### Synthesis of compounds 14a – p

**(4-Amino-2-(cyclopentylamino)thiazol-5-yl)(3,4,5-trimethoxyphenyl)methanone (14a).** General method C was followed using **13a** (40 mg, 0.12 mmol,) and cyclopentylamine (23.2 uL, 2.35 mmol.). The desired product was isolated as an orange-yellow solid (25 mg, 55%). ^1^H NMR (400 MHz, CDCl_3_) δ 7.05 (s, 2H), 3.93 (m, 9H), 2.69 (s, 3H); ^13^C NMR (100 MHz, CDCl_3_) δ 189.9, 164.3, 154.7, 144.0, 142.1, 138.4, 130.8, 104.9, 62.0, 57.1, 56.8, 24.7, 24.5; MS (ESI): *m/z* 363.0 [(M+Na)^+^].

**4-Amino-2-(((exo)-bicyclo[2.2.1]heptan-2-yl)amino)thiazol-5-yl)(3,4,5-trimethoxyphenyl)methanone (14b)**. General method C was followed using **13a** (40 mg, 0.12 mmol,) and exo-2-aminonorbornane (278 uL, 2.35 mmol.). The desired product was isolated as an orange-yellow glassy solid (28 mg, 60%).^1^H NMR (400 MHz, CD_3_OD) δ 7.02 (s, 2H), 5.57 (bd, J = 6.32 Hz, 1H), 3.91 (s, 6H), 3.89 (s, 3H), 3.30 (m, 1H), 2.35 (m, 2H), 1.89 (m, 1H), 1.55 (m, 2H), 1.38 (m, 1H), 1.28 (m, 2H), 1.18 (m, 2H); ^13^C NMR (100 MHz, CDCl_3_) δ 190.2, 164.0, 143.1, 138.4, 154.7, 144.1, 130.3, 104.6, 59.3, 61.8, 57.0, 43.4, 40.2, 37.0, 36.0, 29.3, 27.0; MS (ESI): *m/z* 404.2 [(M+H)^+^].

**(4-Amino-2-(((exo)-bicyclo[2.2.1]heptan-2-yl)amino)thiazol-5-yl)(benzo[d][1,3]dioxol-5-yl)methanone (14c)**. General method C was followed using **13b** (50 mg, 0.17 mmol.) and exo-2-aminonorbornane (394 uL, 3.33 mmol.). The desired product was isolated as a glassy orange-yellow solid (356mg, 58%). ^1^H NMR (400 MHz, CD_3_OD) δ 7.24 (dd, J = 1.72 Hz, J = 8.04 Hz, 1H), 7.13 (d, J = 1.64 Hz, 1H), 6.86 (d, J = 8.04 Hz, 1H), 6.01 (s, 2H), 2.32 (m, 2H), 1.81 (m, 1H), 1.49 (m, 4H), 1.20 (m, 4H); ^13^C NMR (100 MHz, CDCl_3_) δ 190.0, 164.5, 154.5, 144.7, 143.0, 138.5, 130.2, 104.0, 61.9, 57.0, 58.9, 43.5, 40.1, 37.2, 36.1, 29.4, 27.1; MS (ESI): *m/z* 358.1 [(M+H)^+^].

**(4-Amino-2-(((exo)-bicyclo[2.2.1]heptan-2-yl)amino)thiazol-5-yl)(3,4-dihydro-2H-benzo[b][1,4]dioxepin-7-yl)methanone (14d)**. General method C was followed using **13c** (48 mg, 0.15 mmol.) and exo-2-aminonorbornane (348 uL, 2.94 mmol.). The desired product was isolated as a glassy orange-yellow solid (24 mg, 40%). ^1^H NMR (400 MHz, CD_3_OD) δ 7.26 (m, 2H), 6.97 (m, 1H), 4.22 (m, 4H), 2.32 (m, 2H), 2.18 (m, 2H), 1.81 (m, 1H), 1.49 (m, 4H), 1.20 (m, 4H); ^13^C NMR (100 MHz, CDCl_3_) δ 190.1, 164.2, 154.9, 150.0, 143.2, 138.1, 128.3, 122.7, 112.8, 112.4, 72.0, 58.8, 43.3, 40.0, 37.1, 36.3, 32.7, 29.5, 27.0; MS (ESI): *m/z* 386.1 [(M+H)^+^].

**4-Amino-2-(exo-bicyclo[2.2.1]heptan-2-ylamino-thiazol-5-yl)(2-nitrophenyl)methanone (14e)**. General method C was followed using **13d** (40 mg, 0.14 mmol) and exo-2-aminonorbornane (324 uL, 2.74 mmol). The desired product was isolated as orange glassy solid (145 mg, 99%). ^1^H NMR (400 MHz, CDCl_3_) δ 8.08 (d, J = 7.84 Hz, 1H), 7.67 (t, J = 7.52 Hz, 1H), 7.56 (t, J = 7.678 Hz, 2 Hz), 5.65 (bd, J = 6.12 Hz, 1H), 3.17 (bs, 1H), 2.31 (bs, 2H), 1.82 (m, 1H), 1.49 (m, 3H), 1.32 (m, 1H), 1.25 (m, 1H), 1.11 (m, 2H); ^13^C NMR (100 MHz, CD_3_OD) δ 190.9, 165.4, 148.9, 143.7, 137.8, 133.0, 131.3, 130.0, 122.6, 125.6, 59.1, 43.3, 40.1, 36.9, 36.2, 29.2, 27.2; MS (ESI): *m/z* 359.0 [(M+H)^+^].

**4-((4-Amino-5-(2-nitrobenzoyl)thiazol-2-yl)amino)benzenesulfonamide (14f)**. General method C was followed using **13d** (40 mg, 0.14 mmol) and sulfanilamide (472 mg, 2.74 mmol.). The desired product was isolated as a glassy orange solid (12 mg, 14%). ^1^H NMR (400 MHz, CD_3_OD) δ 8.13 (d, J = 8.16 Hz, 1H), 7.83 (m, 5H), 7.68 (m, 1H), 7.61 (dd, J = 1.32 Hz, J = 7.52 Hz, 1H); 190.8, 160.2, 148.7, 144.0, 143.5, 137.7, 133.1, 131.6, 131.2, 130.8, 130.2, 122.5, 125.7, 114.6; MS (ESI): *m/z* 420.0 [(M+H)^+^].

**4-((4-Amino-5-nicotinoylthiazol-2-yl)amino)benzenesulfonamide (14g)**. General method C was followed using **13e** (53 mg, 0.21 mmol) and sulfanilamide (708 mg, 4.11 mmol). The desired product was isolated as a yellow solid (25 mg, 32%). ^1^H NMR (400 MHz, DMSO-d_6_) δ 11.17 (s, 1H), 8.85 (d, J = 1.60 Hz, 1H), 8.68 (dd, J = 1.60 Hz, J = 4.80 Hz, 1H), 8.31 (bs, 2H), 8.05 (m, 1H), 7.79 (s, 4H), 7.51 (m, 1H), 7.27 (s, 2H); ^13^C NMR (100 MHz, CDCl_3_) δ 190.3, 160.4, 156.3, 152.5, 144.3, 143.1, 138.2, 137.3, 131.9, 131.2, 130.9, 125.5, 114.6; MS (ESI): *m/z* 376.0 [(M+H)^+^].

**N-(4-Amino-5-(2-nitrobenzoyl)thiazol-2-yl)piperidine-4-carboxamide (14h)**. *Step 1. Tert-butyl (2-(methylthio)-5-(2-nitrobenzoyl)thiazol-4-yl)carbamate.* Compound **13d** (500 mg, 1.69 mmol.) was dissolved in dchloromethane (10 mL). A solution of di-tert-butyl dicarbonate (406 mg, 1.86 mmol.) in dichloromethane (10 mL) was added, followed by 4-(dimethylamino)pyridine (21 mg, 0.17 mmol.). The resulting solution was stirred at room temperature overnight and then concentrated on a rotary evaporator. The residue was purified by chromatography on silica gel using a gradient of 0 to 2.5% ethyl acetate in dichloromethane to afford the desired product as a yellow crystalline solid (355 mg, 53%). ^1^H NMR (400 MHz, CDCl_3_) δ 10.12 (s, 1H), 8.18 (dd, J = 0.96 Hz, J = 8.16 Hz, 1H), 7.76 (td, J = 1.20 Hz, J = 7.52 Hz, 1H), 7.66 (td, J = 1.48 Hz, J = 8.24 Hz, 1H), 7.49 (dd, J = 1.40 Hz, J = 7.52 Hz, 1H), 2.66 (s, 3H), 1.56 (s, 9H); MS (ESI): m/z 813.2 [(2M+Na)^+^].

*Step 2. Tert-butyl (2-(methylsulfonyl)-5-(2-nitrobenzoyl)thiazol-4-yl)carbamate*. 3-Chloroperbenzoic acid (70%, 382 mg, 1.52 mmol.) was added to a solution of tert-butyl (2-(methylthio)-5-(2-nitrobenzoyl)thiazol-4-yl)carbamate (241 mg, 0.61 mmol.) in dichloromethane (4 mL). The resulting solution was stirred at room temperature for 5 hours and then concentrated on a rotary evaporator. The residue was dissolved in ethyl acetate, washed with saturated aqueous sodium thiosulfate, saturated aqueous sodium bicarbonate and brine, dried over anhydrous sodium sulfate and concentrated on a rotary evaporator. The desired product was isolated as a yellow solid (252 mg, 97%) and used without further purification.^1^H NMR (400 MHz, CDCl_3_) δ 9.77 (s, 1H), 8.27 (dd, J = 1.04 Hz, J = 8.20 Hz, 1H), 7.83 (td, J = 1.12 Hz, J = 7.48 Hz, 1H), 7.76 (td, J = 1.48 Hz, J = 8.12 Hz, 1H), 7.51 (dd, J = 1.40 Hz, J = 7.48 Hz, 1H), 3.39 (s, 3H), 1.57 (s, 9H); MS (ESI): m/z 877.1 [(2M+Na)^+^].

*Step 3. Tert-butyl (2-amino-5-(2-nitrobenzoyl)thiazol-4-yl)carbamate*. A 0.5 M solution of ammonia in dioxane (2.98 mL, 1.49 mmol.) and triethylamine (0.28 mL, 1.99 mmo.l) were added sequentially to a solution of tert-butyl (2-(methylsulfonyl)-5-(2-nitrobenzoyl)thiazol-4-yl)carbamate (318 mg, 0.74 mmol.) in dioxane (10 mL). The reaction was stirred at room temperature for 4 hours and then concentrated on a rotary evaporator. The residue was purified by column chromatography on silica gel using a gradient of 0 to 100% ethyl acetate in hexanes to afford the desired product as an orange solid (133 mg, 49%). ^1^H NMR (400 MHz, CDCl_3_) δ 10.60 (s, 1H), 8.12 (dd, J = 0.96 Hz, J = 8.04 Hz, 1H), 7.71 (td, J = 1.12 Hz, J = 7.48 Hz, 1H), 7.61 (td, J = 1.44 Hz, J = 8.16 Hz, 1H), 7.49 (dd, J = 1.36 Hz, J = 7.48 Hz, 1H), 7.04 (bs, 2H), 1.51 (s, 9H); MS (ESI): m/z 751.3 [(2M+Na)^+^].

*Step 4. Tert-butyl 4-((4-((tert-butoxycarbonyl)amino)-5-(2-nitrobenzoyl)thiazol-2-yl)carbamoyl)piperidine-1-carboxylate*. Tert-butyl (2-amino-5-(2-nitrobenzoyl)thiazol-4-yl)carbamate (50 mg, 0.135 mmol.), N-Boc-piperidine-4-carboxylic acid (31.5 mg, 0.135 mmol.), N-(3-Dimethylaminopropyl)-N′-ethylcarbodiimide hydrochloride (79 mg mg, 0.41 mmol.) and 1-hydroxybenzotriazole hydrate (55.5 mg, 0.41 mmol.) were combined in dichloromethane (10 mL) and stirred at room temperature overnight. The reaction was concentrated down and the residue was partitioned between ethyl acetate and water. The organic layer was separated, washed with water, 1N aqueous hydrochloric acid, 1N sodium hydroxide and brine, dried over anhydrous sodium sulfate and concentrated on a rotary evaporator. The residue was purified by column chromatography on silica gel using 0 to 100% ethyl acetate in hexanes to afford the desired product as a yellow solid (40 mg, 51%). ^1^H NMR (400 MHz, CDCl_3_) δ 10.23 (s, 1H), 8.20 (d, J = 7.48 Hz, 1H), 7.75 (t, J = 7.40 Hz, 1H), 7.67 (td, J = 1.32 Hz, J = 8.24 Hz, 1H), 7.49 (dd, J = 1.00 Hz, J = 7.44 Hz, 1H), 4.12 (m, 2H), 2.81 (t, J = 11.80 Hz, 2H), 2.48 (t, J = 11.00 Hz, 1H), 1.85 (m, 2H), 1.65 (m, 2H), 1.56 (s, 9H), 1.45 (s, 9H); MS (ESI): m/z 598.5 [(M+Na)^+^].

*Step 5. N-(4-amino-5-(2-nitrobenzoyl)thiazol-2-yl)piperidine-4-carboxamide **(14h)***. Tert-butyl 4-((4-((tert-butoxycarbonyl)amino)-5-(2-nitrobenzoyl)thiazol-2-yl)carbamoyl)piperidine-1-carboxylate (36 mg, 0.06 mmol.) was dissolved into a solution of 4N hydrochloric acid in dioxane (3 mL). The reaction was stirred at room temperature for two hours. Acetonitrile (2 mL) was added to dissolve the resulting precipitate and stirring was continued for one hour. The reaction was then concentrated on a rotary evaporator. The residue was partitioned between ethyl acetate and 1N aqueous hydrochloric acid. The aqueous layer was separated, washed with ethyl acetate and basified to pH 12 with 1N aqueous sodium hydroxide. The resulting basic aqueous solution was extracted with three portions of ethyl acetate, saturated with sodium chloride and then extracted twice more with ethyl acetate. The combined orgnanic layers were dried over anhydrous sodium sulfate and concentrated on a rotary evaporator to afford the desired product as an orange solid (11.4 mg, 49%) that was pure enough to use without further purification. ^1^H NMR (400 MHz, CD_3_OD) δ 8.15 (dd, J = 0.92 Hz, J = 8.20 Hz, 1H), 7.80 (td, J = 1.12 Hz, J = 7.48 Hz, 1H), 7.69 (td, J = 1.44 Hz, J = 8.20 Hz, 1H), 7.60 (dd, J = 1.32 Hz, J = 7.52 Hz, 1H), 3.17 (m, 2H), 2.75 (td, J = 2.96 Hz, J = 12.48 Hz, 2H), 2.60 (m, 1H), 1.88 (m, 2H), 1.71 (m, 2H); ^13^C NMR (100 MHz, CDCl_3_) δ 190.7, 173.8, 163.4, 149.2, 143.0, 138.2, 135.6, 134.3, 131.4, 129.9, 124.8, 45.9, 39.3, 33.1; MS (ESI): m/z 376.7 [(M+H)^+^].

**(4-Amino-2-((tetrahydro-2H-pyran-4-yl)amino)thiazol-5-yl)(2-nitrophenyl)methanone (14i)**. General method C was followed using **13d** (40 mg, 0.12 mmol.) and 4-aminotetrahydropyran (284 uL, 2.74 mmol.). The desired product was isolated as an orange solid (18 mg, 44%). ^1^H NMR (400 MHz, CD_3_OD) δ 8.09 (dd, J = 0.96 Hz, J = 8.20 Hz, 1H), 7.76 (td, J = 1.16 Hz, J = 7.52 Hz, 1H), 7.64 (td, J = 1.44 Hz, J = 8.20 Hz, 1H), 7.56 (dd, J = 1.27 Hz, J = 7.56 Hz, 1H), 3.93 (m, 2H), 3.47 (m, 2H), 3.35 (m, 1H), 1.96 (m, 2H), 1.55 (m, 2H); ^13^C NMR (100 MHz, CDCl_3_) δ 190.8, 164.1, 149.0, 143.1, 138.2, 135.5, 134.3, 131.4, 129.1, 124.9, 66.4, 55.6, 37.7; MS (ESI): m/z 349.7 [(M+H)^+^].

**(4-Amino-2-(((exo)-bicyclo[2.2.1]heptan-2-yl)amino)thiazol-5-yl)(3-nitrophenyl)methanone (14j)**. General method C was followed using **13f** (40 mg, 0.14 mmol.) and exo-2-aminonorbornane (324 uL, 2.74 mmol). The desired product was isolated as a glassy orange solid (23 mg, 46%). ^1^H NMR (400 MHz, CDCl_3_) δ 8.60 (t, J = 1.88 Hz, 1H), 8.30 (m, 1H), 8.07 (dt, J = 1.24 Hz, J = 7.68 Hz, 1H), 7.62 (t, J = 7.96 Hz, 1H), 5.83 (bs, 1H), 3.30 (bs, 1H), 2.35 (m, 2H), 1.90 (m, 1H), 1.54 (m, 3H), 1.38 (m, 1H), 1.31 (m, 1H), 1.16 (m, 2H); ^13^C NMR (100 MHz, CDCl_3_) δ 190.3, 164.0, 149.1, 143.1, 138.2, 137.0, 136.6, 130.5, 128.6, 124.4, 57.1, 43.2, 40.0, 37.1, 35.9, 29.2, 27.3; MS (ESI): m/z 359.8 [(M+H)^+^].

**(4-Amino-2-(((exo)-bicyclo[2.2.1]heptan-2-yl)amino)thiazol-5-yl)(4-nitrophenyl)methanone (14k)**. General method C was followed using **13g** (40 mg, 0.14 mmol.) and exo-2-aminonorbornane (324 uL, 2.74 mmol). The desired product was isolated as orange glassy solid (26 mg, 52%). ^1^H NMR (400 MHz, CDCl_3_) δ 8.28 (m, 2H), 8.87 (m, 2H), 5.75 (bs, 1H), 3.27 (bs, 1H), 2.35 (m, 2H), 1.90 (m, 1H), 1.54 (m, 3H), 1.38 (m, 1H), 1.27 (m, 1H), 1.16 (m, 2H); 57.0, 43.4, 40.2, 37.0, 36.0, 29.3, 27.0; ^13^C NMR (100 MHz, CDCl_3_) δ 190.9, 164.2, 143.3, 138.4, 153.2, 145.4, 129.8, s124.9, 57.3, 43.3, 40.2, 37.4, 36.3, 30.2, 27.4; MS (ESI): m/z 359.7 [(M+H)^+^].

**2-(4-Amino-2-(((exo)-bicyclo[2.2.1]heptan-2-yl)amino)thiazole-5-carbonyl)benzonitrile (14l)**. General method C was followed using **13h** (47 mg, 0.17 mmol.) and exo-2-amnonorbornane (393 uL, 3.33 mmol.). The desired product was isolated as a glassy yellow solid (19.5 mg, 34%). ^1^H NMR (400 MHz, CDCl_3_) δ 7.76 (d, J = 7.56 Hz, 1H), 7.65 (m, 2H), 7.52 (td, J = 1.56 Hz, J = 7.56 Hz, 1H), 6.08 (bs, 1H), 3.22 (bs, 1H), 2.34 (bs, 2H), 1.86 (m, 1H), 1.53 (m, 2H), 1.32 (m, 3H), 1.14 (m, 2H); ^13^C NMR (100 MHz, CDCl_3_) δ 190.5, 164.0, 143.3, 138.4, 137.5, 134.3, 134.0, 133.1, 131.4, 116.7, 113.3, 57.5, 43.2, 40.1, 37.2, 36.1, 30.3, 27.2; MS (ESI): m/z 339.7 [(M+H)^+^].

**(4-Amino-2-(((exo)-bicyclo[2.2.1]heptan-2-yl)amino)thiazol-5-yl)(7-nitro-2,3-dihydrobenzo[b]-[1,4]dioxin-6-yl)methanone (14m)**. General method C was followed using **13i** (42 mg, 0.12 mmol.) and exo-2-aminonorbornane (284 uL, 2.74 mmol.). The desired product was isolated as an orange sold (14.6 mg, 31%). ^1^H NMR (400 MHz, CD_3_OD) δ 7.66 (s, 1H), 6.93 (s, 1H), 4.37 (m, 4H), 2.29 (m, 2H), 1.78 (m, 1H), 1.47 (m, 5H), 1.19 (m, 3H); ^13^C NMR (100 MHz, CDCl_3_) δ 190.4, 164.3, 156.3, 154.8, 143.0, 141.4,138.0, 121.3, 116.3, 110.1, 65.1, 64.9,, 57.4, 43.4, 40.3, 37.3, 36.2, 30.5, 27.4; MS (ESI): m/z 417.1 [(M+H)^+^].

**(4-Amino-2-(((exo)-bicyclo[2.2.1]heptan-2-yl)amino)thiazol-5-yl)(6-nitrobenzo[d][1,3]dioxol-5-yl)methanone (14n)**. Method C was followed using **13**j (41 mg, 0.12 mmol.) and exo-2-aminonorbornane (284 uL, 2.74 mmol.). The desired product was isolated as a glassy orange solid (44 mg, 92%). ^1^H NMR (400 MHz, CDCl_3_) δ 7.57 (s, 1H), 6.88 (s, 1H), 6.17 (s, 2H), 5.71 (bd, J = 6.04 Hz, 1H), 3.16 (bs, 1H), 2.32 (m, 2H), 1.83 (m, 1H), 1.51 (m, 3H), 1.34 (m, 1H), 1.23 (m, 2H), 1.12 (m, 2H); ^13^C NMR (100 MHz, CDCl_3_) δ 190.5, 164.2, 156.1, 154.7, 143.1, 141.8, 138.3, 122.4, 116.5, 110.3, 102.0, 57.7, 43.1, 40.2, 37.0, 36.3, 30.5, 27.6; MS (ESI): m/z 403.0 [(M+H)^+^].

**(4-Amino-2-(((exo)-bicyclo[2.2.1]heptan-2-yl)amino)thiazol-5-yl)(benzo[c][1,2,5]oxadiazol-4-yl)methanone (14o)**. General method C was followed using **13k** (50 mg, 0.17 mmol.) and exo-2-aminonorbornane (393 uL, 3.33 mmol.). The desired product was isolated as a glassy reddish-brown solid (20 mg, 33%). ^1^H NMR (400 MHz, CD_3_OD) δ 8.99 (dd, J = 0.48 Hz, J = 9.04 Hz, 1H), 7.72 (d, J = 6.52 Hz, 1H), 7.59 (m, 1H), 3.30 (bs, 1H), 2.31 (m, 2H), 1.79 (m, 1H), 1.48 (m, 3H), 1.22 (m, 4H); ^13^C NMR (100 MHz, CDCl_3_) δ 190.3, 164.0, 149.4, 146.3, 143.1, 138.0, 132.0, 129.1, 120.6, 116.8, 57.8, 43.0, 40.1, 37.1, 36.2, 30.3, 27.9; MS (ESI): m/z 356.1 [(M+H)^+^].

**(4-Amino-2-(((exo)-bicyclo[2.2.1]heptan-2-yl)amino)thiazol-5-yl)(benzo[c][1,2,5]oxadiazol-5-yl)methanone (14p)**. General method Cwas followed using 13l (50 mg, 0.17 mmol.) and exo-2-aminonorbornane (393 uL, 3.33 mmol.). The desired product was isolated as a glassy orange solid (17.7 mg, 29%). ^1^H NMR (400 MHz, CDCl_3_) δ 8.17 (t, J = 1.12 Hz, 1H), 7.88 (dd, J = 1.00 Hz, J = 9.32 Hz, 1H), 7.77 (dd, J = 1.28 Hz, J = 9.28 Hz, 1H), 6.00 (bs, 1H), 3.27 (bs, 1H), 2.35 (m, 2H), 1.90 (m, 1H), 1.54 (m, 2H), 1.30 (m, 3H), 1.15 (m, 2H); ^13^C NMR (100 MHz, CDCl_3_) δ 190.2, 164.3, 143.1, 138.3, 153.4, 149.4, 132.8, 129.0, 116.7, 113.1, 57.6, 42.9, 40.0, 37.2, 36.0, 30.1, 27.6; MS (ESI): m/z 356.1 [(M+H)^+^].

### Synthesis of compounds 17a-g

**N-(2-(Cyclopentylamino)-5-(2,3-dihydrobenzo[b][1,4]dioxine-6-carbonyl)thiazol-4-yl)acetamide (17a)**. *Step 1. N-(5-(2,3-Dihydrobenzo[b][1,4]dioxine-6-carbonyl)-2-(methylthio)thiazol-4-yl)acetamide*. Pyridine (39 uL, 0.49 mmol.) and acetyl chloride (35 uL, 0.49 mmol.) were added to a suspension of **10** (100 mg, 0.32 mmol.) in acetonitrile (4 mL). Dichloromethane (3 mL) was added to fully dissolve all of the starting materials. The reaction was stirred at room temperature for 24 hours and then additional portions of pyridine (39 uL, 0.49 mmol.) and acetyl chloride (35 uL, 0.49 mmol.) were added sequentially. The resulting reaction mixture was stirred at room temperature overnight. The mixture was concentrated on a rotary evaporator and the residue was partitioned between ethyl acetate and water.

The organic layer was separated, washed with brine, dried over anhydrous sodium sulfate and concentrated on a rotary evaporator. The residue was purified by column chromatography on silica gel using 0 to 100% ethyl acetate in hexanes to afford the desired product as an orange-yellow solid (69 mg, 61%). ^1^H NMR (400 MHz, CDCl_3_) δ 11.06 (s, 1H), 7.35 (m, 2H), 6.93 (d, J = 8.24 Hz, 1H), 4.32 (m, 4H), 2.74 (s, 3H), 2.43 (s, 3H); MS (ESI): m/z 723.1 [(2M+Na)^+^].

*Step 2. N-(5-(2,3-dihydrobenzo[b][1,4]dioxine-6-carbonyl)-2-(methylsulfonyl)thiazol-4-yl)acetamide*. Pepared using the same procedure as described for the synthesis of compound **14h**, Step 2 using N-(5-(2,3-dihydrobenzo[b][1,4]dioxine-6-carbonyl)-2-(methylthio)thiazol-4-yl)acetamide (61 mg, 0.17 mmol.) and 3-chloroperbenzoic acid (70%, 107 mg, 0.44 mmol.). The product was obtained as a yellow solid (64 mg, 96%). ^1^H NMR (400 MHz, CDCl_3_) δ 10.46 (s, 1H), 7.41 (m, 2H), 6.98 (m, 1H), 4.34 (m, 4H), 3.40 (s, 3H), 2.40 (s, 3H); ESIMS: m/z 787.0 [(2M+Na)^+^].

*Step 3. N-(2-(Cyclopentylamino)-5-(2,3-dihydrobenzo[b][1,4]dioxine-6-carbonyl)thiazol-4-yl)acetamide **(17a)***. Cyclopentylamine (10 uL, 0.11 mmol.) and triethylamine (20 uL, 0.14 mmol.) were added sequentially to a solution of N-(5-(2,3-dihydrobenzo[b][1,4]dioxine-6-carbonyl)-2-(methylsulfonyl)thiazol-4-yl)acetamide (20 mg, 0.05 mmol.) in anhydrous dioxane (2 mL). The reaction solution was stirred at room temperature overnight and then concentrated on a rotary evaporator. The residue was purified by column chromatography on silica gel using 0 to 100% ethyl acetate in hexanes to afford the desired product as an orange-yellow glassy solid (14 mg, 69%). ^1^H NMR (400 MHz, CDCl_3_) δ 11.83 (s, 1H), 7.33 (m, 2H), 6.92 (d, J = 8.24 Hz, 1H), 6.27 (bs,1H), 4.31 (m, 4H), 3.75 (m, 1H), 2.31 (s, 3H), 2.04 (m, 2H), 1.66 (m, 6H); MS (ESI): m/z 388.1 [(M+H)^+^].

Tert-butyl (2-(((1S,2S,4R)-bicyclo[2.2.1]heptan-2-yl)amino)-5-(2,3-dihydrobenzo[b][1,4]dioxine-6-carbonyl)thiazol-4-yl)carbamate (17b). *Step 1. Tert-butyl (5-(2,3-dihydrobenzo[b][1,4]dioxine-6-carbonyl)-2-(methylthio)thiazol-4-yl)carbamate*. Compound **10** (500 mg, 1.62 mmol.) was partially dissolved into dichloromethane (10 mL). A solution of di-tert-butyl dicarbonate (389 mg, 1.78 mmol.) in dichloromethane (10 mL) was added followed by 4-(dimethylamino)pyridine (20 mg, 0.16 mmol.). The resulting solution was stirred at room temperature overnight and then concentrated on a rotary evaporator. The resiue was purified by column chromatography on silica gel using a gradient of 0 to 10% ethyl acetate in dichloromethane to afford the desired product as a pale yellow foamy solid (320 mg, 48%).^1^H NMR (400 MHz, CDCl_3_) δ 10.59 (s, 1H), 7.35 (m, 2H), 6.93 (m, 1H), 4.31 (m, 4H), 2.73 (s, 3H), 1.54 (s, 9H); MS (ESI): m/z 431.0 [(M+Na)^+^].

*Step 2. Tert-butyl (5-(2,3-dihydrobenzo[b][1,4]dioxine-6-carbonyl)-2-(methylsulfonyl)thiazol-4-yl)carbamate*. Prepared using the same procedure as described for the synthesis of compound **14h** Step 2 using tert-butyl (5-(2,3-dihydrobenzo[b][1,4]dioxine-6-carbonyl)-2-(methylthio)thiazol-4-yl)carbamate (100 mg, 0.24 mmol.) and 3-chloroperbenzoic acid (70%, 154 mg, 0.61 mmol.). Product was obtained as a crystalline yellow solid (109 mg, 100%). ^1^H NMR (400 MHz, CDCl_3_) δ 9.96 (s, 1H), 7.40 (m, 2H), 6.97 (m, 1H), 4.32 (m, 4H), 3.41 (s, 3H), 1.54 (s, 9H); MS (ESI): m/z 463.0 [(M+Na)^+^].

Step 3. Tert-butyl (2-(((exo)-bicyclo[2.2.1]heptan-2-yl)amino)-5-(2,3-dihydrobenzo[b][1,4]dioxine-6- carbonyl)thiazol-4-yl)carbamate **(17b).** Prepared using the same procedure as described for the synthesis of compound **17a**, Step 3 using tert-butyl (5-(2,3-dihydrobenzo[b][1,4]dioxine-6-carbonyl)-2- (methylsulfonyl)thiazol-4-yl)carbamate (50 mg, 0.11 mmol.), exo-2-aminonorbornane (27 uL, 0.23 mmol.), and triethylamine (42 uL, 0.14 mmol.). Product was obtained as a glassy yellow solid (35 mg, 65%). ^1^H-NMR (400 MHz, CDCl_3_) δ 11.16 (s, 1H), 7.32 (m, 2H), 6.92 (d, J = 8.24 Hz, 1H), 6.12 (m, 1H), 4.31 (m, 4H), 3.18 (m, 1H), 2.34 (m, 2H), 1.88 (m, 1H), 1.58 (m, 1H), 1.51(s, 9H), 1.26 (m, 6H); ^13^C NMR (100 MHz, CDCl_3_) δ 190.1, 164.2, 153.9, 155.4, 150.2, 143.2, 138.1, 128.2, 122.7, 112.6, 112.2, 80.5, 65.3, 65.1, 57.5, 43.0, 40.2, 37.3, 36.1, 30.3, 29.4, 27.8; MS (ESI) *m/z* 472.2 [(M+H)^+^].

**N-(2-(((Exo)-bicyclo[2.2.1]heptan-2-yl)amino)-5-(2,3-dihydrobenzo[b][1,4]dioxine-6- carbonyl)thiazol-4-yl)butyramide (17c).** Butyryl chloride (6.4 uL, 0.062 mmol.) was added to a solution of compound **12ak** (14.3 mg, 0.039 mmol.) and pyridine (4.7 uL, 0.058 mmol.) in dichloromethane (500 uL). The reaction was stirred at room temperature overnight and then concentrated on a rotary evaporator. The residue was purified by column chromatography on silica gel using a gradient of 0 to 100% ethyl acetate in hexanes to afford the desired product as a glassy yellow solid (12.8 g, 75%). ^1^H-NMR (400 MHz, CDCl_3_) δ 11.87 (s, 1H), 7.32 (m, 2H), 6.92 (m, 1H), 6.27 (bs, 1H), 4.31 (m, 4H), 3.21 (m, 1H), 2.48 (t, J = 7.36 Hz, 2H), 2.34 (m, 2H), 1.89 (m, 1H), 1.80 (q, J = 7.48 Hz, 2H), 1.68 (m, 1H), 1.55 (m, 2H), 1.39 (m, 1H), 1.30 (m, 2H), 1.25 (t, J = 7.20 Hz, 3H), 1.15 (m, 1H); ^13^C NMR (100 MHz, CDCl_3_) δ 190.0, 172.9, 164.0, 143.2, 138.3, 155.5, 150.3, 128.2, 122.7, 112.7, 112.3, 65.3, 57.7, 43.3, 40.7, 40.4, 37.2, 36.0, 30.2, 29.5, 27.9, 20.3, 15.7; MS (ESI) *m/z* 442.1 [(M+H)^+^].

**N-(2-(((Exo)-bicyclo[2.2.1]heptan-2-yl)amino)-5-(2,3-dihydrobenzo[b][1,4]dioxine-6- carbonyl)thiazol-4-yl)acetamide (17d)**. Prepared using the same procedure as described for compound **17a**, Step 3 using N-(5-(2,3-dihydrobenzo[b][1,4]dioxine-6-carbonyl)-2-(methylsulfonyl)thiazol-4- yl)acetamide (20 mg, 0.05 mmol.), exo-2-aminonorbornane (12 uL, 0.10 mmol.), and triethylamine (20 uL, 0.14 mmol.). Product was obtained as a glassy yellow solid (17 mg, 79%). ^1^H-NMR (400 MHz, CDCl_3_) δ 11.83 (s, 1H), 7.32 (m, 2H), 6.93 (d, J = 8.36 Hz, 1H), 6.19 (bs, 1H), 4.31 (m, 4H), 3.21 (m, 1H), 2.35 (m, 2H), 2.31 (s, 3H), 1.89 (m, 1H), 1.57 (m, 2H), 1.40 (m, 1H), 1.30 (m, 2H), 1.17 (m, 2H); ^13^C NMR (100 MHz, CDCl_3_) δ 190.1, 169.6, 164.1, 143.3, 138.4, 155.7, 150.4, 128.1, 122.6, 112.4, 112.1, 65.5, 65.3, 57.8, 43.3, 40.3, 37.4, 36.2, 30.3, 27.9, 24.8; MS (ESI) *m/z* 414.1 [(M+H)^+^].

N-(2-(Cyclopentylamino)-5-(2,3-dihydrobenzo[b][1,4]dioxine-6-carbonyl)thiazol-4-yl)-5- ((3aS,4S,6aR)-2-oxohexahydro-1H-thieno[3,4-d]imidazol-4-yl)pentanamide (17e). *Step 1. 5- ((3aS,4S,6aR)-2-oxohexahydro-1H-thieno[3,4-d]imidazol-4-yl)pentanoyl chloride*. D-Biotin (42.5 mg, 0.17 mmol.) was dissolved in thionyl chloride (2 mL) and stirred at room temperature for one hour. The resulting orange-yellow liquid was concentrated on a rotary evaporator. The residual oil was treated with toluene and re-concentrated twice to remove any remaining thionyl chloride. The product was used immediately in the following reaction without further purification or analysis.

*Step 2. N-(2-(cyclopentylamino)-5-(2,3-dihydrobenzo[b][1,4]dioxine-6-carbonyl)thiazol-4-yl)-5- ((3aS,4S,6aR)-2-oxohexahydro-1H-thieno[3,4-d]imidazol-4-yl)pentanamide **(17e)***. A solution of 5- ((3aS,4S,6aR)-2-oxohexahydro-1H-thieno[3,4-d]imidazol-4-yl)pentanoyl chloride (theoretically 0.17 mmol.) in anhydrous tetrahydrofuran (5 mL) was added dropwise to a second solution of compound **12h** (50 mg, 0.14 mmol.) and triethylamine (40.4 uL, 0.29 mmol.) in anhydrous tetrahydrofuran (3 mL). The resulting solution was stirred at room temperature overnight and then concentrated on a rotary evaporator. The crude product was purified by chromatography on silica gel using a gradient of 0 to 10% methanol in dichloromethane to afford the desired product as a pale yellow solid (13.6 mg, 16%). ^1^H NMR (400 MHz, CD_3_OD) δ 7.28 (m, 2H), 6.91 (d, J = 8.28 Hz, 1H), 4.49 (m, 1H), 4.31 (m, 5H), 3.23 (m, 1H), 2.93 (dd, J = 5.00 Hz, J = 12.76 Hz, 1H), 2.71 (m, 2H), 2.06 (m, 2H), 1.65 (m, 12H), 1.31 (m, 2H); ^13^C NMR (100 MHz, CDCl_3_) δ 190.1, 172.8, 165.9, 164.2, 143.1, 138.4, 155.5, 150.0, 128.1, 122.6, 112.5, 112.2, 65.4, 65.2, 64.4, 63.2, 56.5, 56.2, 42.0, 39.1, 25.6, 32.9,28.8, 25.0, 24.4; MS (ESI): m/z 572.2 [(M+H)^+^].

**2-(((Exo)-bicyclo[2.2.1]heptan-2-yl)amino)thiazol-5-yl)(3,4-dimethoxyphenyl)methanone (17f).** *Step 1. (3,4-Dimethoxyphenyl)(2-(methylthio)thiazol-5-yl)methanone*. A solution of 2- (methylthio)thiazole (392 uL, 3.81 mmol.) in anhydrous tetrahydrofuran (7.5 mL) was cooled to −55°C under nitrogen. N-Butyl lithium (1.6 M in hexanes, 2.38 mL, 3.81 mmol.) was added dropwise, keeping the reaction solution temperature below −40°C. A solution of 3,4-dimethoxybenzonitrile (311 mg, 1.91 mmol) in anhydrous tetrahydrofuran (2.5 mL) was then added dropwise while keeping the reaction temperature between −35°C and −20°C. Stirring was continued while maintaining this temperature range for 50 minutes. The reaction was quenched by adding 1M aqueous acetic acid (11.4 mL, 11.43 mmol.) at −20°C. The reaction mixture was allowed to come up to room temperature while stirring overnight and then diluted with ethyl acetate and water. The aqueous phase was extracted with two additional portions of ethyl acetate. The combined organic layers were washed with 50% aqueous potassium carbonate solution twice and brine, dried over anhydrous sodium sulfate and concentrated on a rotary evaporator. The crude product was purified by chromatography on silica gel using a gradient of 0 to 80% of ethyl acetate in hexanes to afford the titled compound as an orange-yellow solid (212 mg, 19%).^1^H NMR (400 MHz, CDCl_3_) δ 8.08 (s, 1H), 7.53 (dd, J = 8.32 Hz, J = 2.0 Hz, 1H), 7.43 (d, J = 2.0 Hz, 1H), 6.94 (d, J = 8.36 Hz, 1H), 3.98 (s, 3H), 3.95 (s, 3H), 2.76 (s, 3H); ; MS (ESI): *m/z* 296.3 [(M+H)^+^].

*Step 2. 2-(((Exo)-bicyclo[2.2.1]heptan-2-yl)amino)thiazol-5-yl)(3,4-dimethoxyphenyl)methanone **(17f).*** (3,4-Dimethoxyphenyl)(2-(methylthio)thiazol-5-yl)methanone (50 mg, 0.1693 mmol.), exo-2-aminonorbornane (401 uL, 3.386 mmol.) and ethanol (2 mL) were combined in a glass pressure tube. This mixture was heated at 100°C for 23 hours. The reaction was then cooled to room temperature and concentrated on a rotary evaporator. The residue was purified by chromatography on silica gel using a gradient solvent system of 0 to 100% of ethyl acetate in hexanes to afford the titled compound as a pale yellow solid (20 mg, 33%).^1^H NMR (400 MHz, CDCl_3_) δ 7.72 (s, 1H), 7.46 (dd, J = 8.28 Hz, J = 1.96 Hz, 1H), 7.37 (d, J = 1.96 Hz, 1H), 6.92 (d, J = 8.32 Hz, 1H), 6.51 (bs, 1H), 3.96 (s, 3H), 3.94 (s, 3H), 3.34 (bs, 1H), 2.44 (bd, J = 4.16 Hz, 1H), 2.37 (bs, 1H), 1.94 (m, 1H), 1.58 (m, 2H), 1.46 (m, 1H), 1.36 (m, 1H), 1.28 (m, 2H), 1.20 (m. 1H); ^13^C NMR (100 MHz, CDCl_3_) δ 189.8, 163.1, 155.2, 150.5, 137.4, 128.9, 123.4, 122.3, 112.1, 111.7, 57.9, 57.6, 43.1, 40.0, 37.2, 36.1, 30.2, 27.7; MS (ESI): *m/z* 359.1 [(M+H)^+^].

**(2-(((Exo)-bicyclo[2.2.1]heptan-2-yl)amino)thiazol-5-yl)(2,3-dihydrobenzo[b][1,4]dioxin-6- yl)methanone (17g)**. *Step 1. (2,3-Dihydrobenzo[b][1,4]dioxin-6-yl)(2-(methylthio)thiazol-5- yl)methanone*. Prepared using the same procedure as described for the synthesis of compound **17f**, Step 1 using 2-(methylthio)thiazole (392 uL, 3.81 mmol.), N-Butyl lithium (1.6 M in hexanes, 2.38 mL, 3.81 mmol.) and 2,3-dihydro-1,4-benzodioxine-6-carbonitrile (307 mg, 1.91 mmol.). The desired product was isolated as a pale yellow solid (221 mg, 40%). ^1^H-NMR (400 MHz, CDCl_3_) δ 8.07 (s, 1H), 7.42 (m, 2H), 6.96 (d, J = 8.24 Hz, 1H), 4.32 (m, 4H), 2.75 (s, 3H); MS (ESI) *m/z* 294.1 [(M+H)^+^].

*Step 2. (2-(((Exo)-bicyclo[2.2.1]heptan-2-yl)amino)thiazol-5-yl)(2,3-dihydrobenzo[b][1,4]dioxin-6- yl)methanone **(17g)***. Prepared using the same procedure as described for the synthesis of compound **17f**, Step 2 using (2,3-dihydrobenzo[b][1,4]dioxin-6-yl)(2-(methylthio)thiazol-5-yl)methanone (50 mg, 0.17 mmol.) and exo-2-aminonorbornane (404 uL, 3.41 mmol.). The desired product was isolated as a pale yellow solid (21 mg, 35%). ^1^H-NMR (400 MHz, CDCl_3_) δ 7.70 (s, 1H), 7.34 (m, 2H), 6.93 (d, J = 8.28 Hz, 1H), 6.49 (bs, 1H), 4.31 (m, 4H), 3.32 (bs, 1H), 2.43 (bd, J = 4.12 Hz,1H), 2.37 (m, 1H), 1.93 (m, 1H), 1.56 (m, 2H), 1.44 (m, 1H), 1.37 (m, 1H), 1.24 (m, 3H); ^13^C NMR (100 MHz, CDCl_3_) δ 189.8, 163.0, 155.3, 150.6, 137.7, 128.7, 123.4, 122.0, 112.4, 111.0, 65.5, 65.3, 57.7, 43.2, 40.3, 37.3, 36.4, 30.3, 27.9; MS (ESI) *m/z* 357.1 [(M+H)^+^].

### Assessment of maximum kinetic aqueous solubility

Solubility assays were performed using Millipore MultiScreen^®^HTS-PFC Filter Plates designed for solubility assays (EMD Millipore, Billerica, MA). The 96-well plates consist of two chambers separated by a filter. Liquid handling was performed using JANUS^®^ Verispan and MTD workstations (Perkin Elmer, Waltham, MA). 4 µL of drug solutions (10 mM in DMSO) are added to 196 µL of phosphate-buffered saline (Millipore-defined universal buffer—45 mM potassium phosphate, 45 mM sodium acetate, 45 mM ethanolamine, pH = 7.4) in the top chamber to give a final DMSO concentration of 2% and a theoretical drug concentration of 200 µM. Plates are gently shaken for 90 min and then subjected to vacuum. Insoluble drug is captured on the filter. 160 µL of the filtrate is transferred to 96- well Griener UV Star^®^ analysis plates (Sigma–Aldrich, St. Louis, MO) containing 40 µL of acetonitrile. The drug concentration in the filtrate is measured by UV absorbance on a Spectromax^®^ Plus microplate reader (Molecular Devices, Sunnyvale, CA) using Softmax Pro software v. 5.4.5. Absorbances at 5 wavelengths (280, 300, 320, 340, and 360 nM) were summed to generate the UV signal. Assays were performed in triplicate. Standard curves were generated by adding 4 µL of 50× of five concentrations of test compounds in DMSO to 40 µL of acetonitrile in UV Star plates followed by 156 µL of the appropriate solubility medium. Analysis and statistics were performed using GraphPad^®^ Prism v. 5.04. Data are reported as the maximum concentration observed in the filtrate.

## Data and software availability

The atomic coordinates and structure factors have been deposited to Protein Data Bank with accession number 6W9E.

The comparative modeling pipeline data generated in this study are available upon request to the corresponding author.

## Supplemental Information

Supporting Information. The SMILES strings for the 77 MC analogs reported in this study (**Table S1**).

## List of abbreviations

3D: Three Dimensional
Å: Angstrom
Abl: Abelson Murine Leukemia
AcOH: Acetic Acid
ATP: Adeonsine Triphosphate
BOC: tert-Butoxycarbonyl
Cd: Compound
CDCl_3_: Deuterated Chloroform
CDK: Cyclin Dependent Kinase
CD_3_OD: Deuterated Methanol
DMAP: 4-(dimethylaminomethyl)pyridine
DMF: Dimethylformamide
DMSO: Dimethylsulfoxide
DMSO-d6: Deuterated Dimethylsulfoxide
ESI: Electrospray Ionization
FBS: Fetal Bovine Serum
GFP: Green Fluorescent Protein
HCl: Hydrochloric Acid
HDAC: Histone Deacetylase
HOBT: 1-Hydroxybenztriazole
K_2_CO_3_: Potassium Carbonate
KOtBu: Potassium tert-Butoxide
LC/MS: Liquid Chromatography/Mass Spectrometry
MCPBA: m-Chloroperbenzoic Acid
n-BuLi: n-Butyllithium
mg: milligram
mmol.: Millimoles
MHz: Megahertz
uL: Microliters
uM: Micromolar
nM: Nanomolar
NCI: National Cancer Institute
N. D.: Not Determined
NIH: National Institutes of Health
NMR: Nuclear Magnetic Resonance
nPr: n-Propyl
PDB: Protein Data Bank
PBS: Phosphate-Buffered Saline
QTOF: Quadrapole Time-Of-Flight
REU: Rosetta Energy Units
RMSD: Root Mean Square Deviation
RT: Room Temperature
SAR: Structure-Activity Relationship
tBu: tert-Butyl
TCEP: Tris(2-carboxyethyl)phosphine hydrochloride
THF: Tetrahydrofuran
XSEDE: Extreme Science and Engineering Discovery Environment

## Acknowledgements

We thank Vivek Modi and Roland Dunbrack for providing their list of active kinase structures ahead of publication. Computational resources were provided through Extreme Science and Engineering Discovery Environment (XSEDE) allocation MCB130049, which is supported by National Science Foundation grant number ACI-1053575. This work was supported by grant CHE-1836950 from the National Science Foundation, grant R01GM123336 from the National Institute of General Medical Sciences of the National Institutes of Health, and the NIH/NCI Cancer Center Support Grant P30CA006927.

---

**Table S1:**
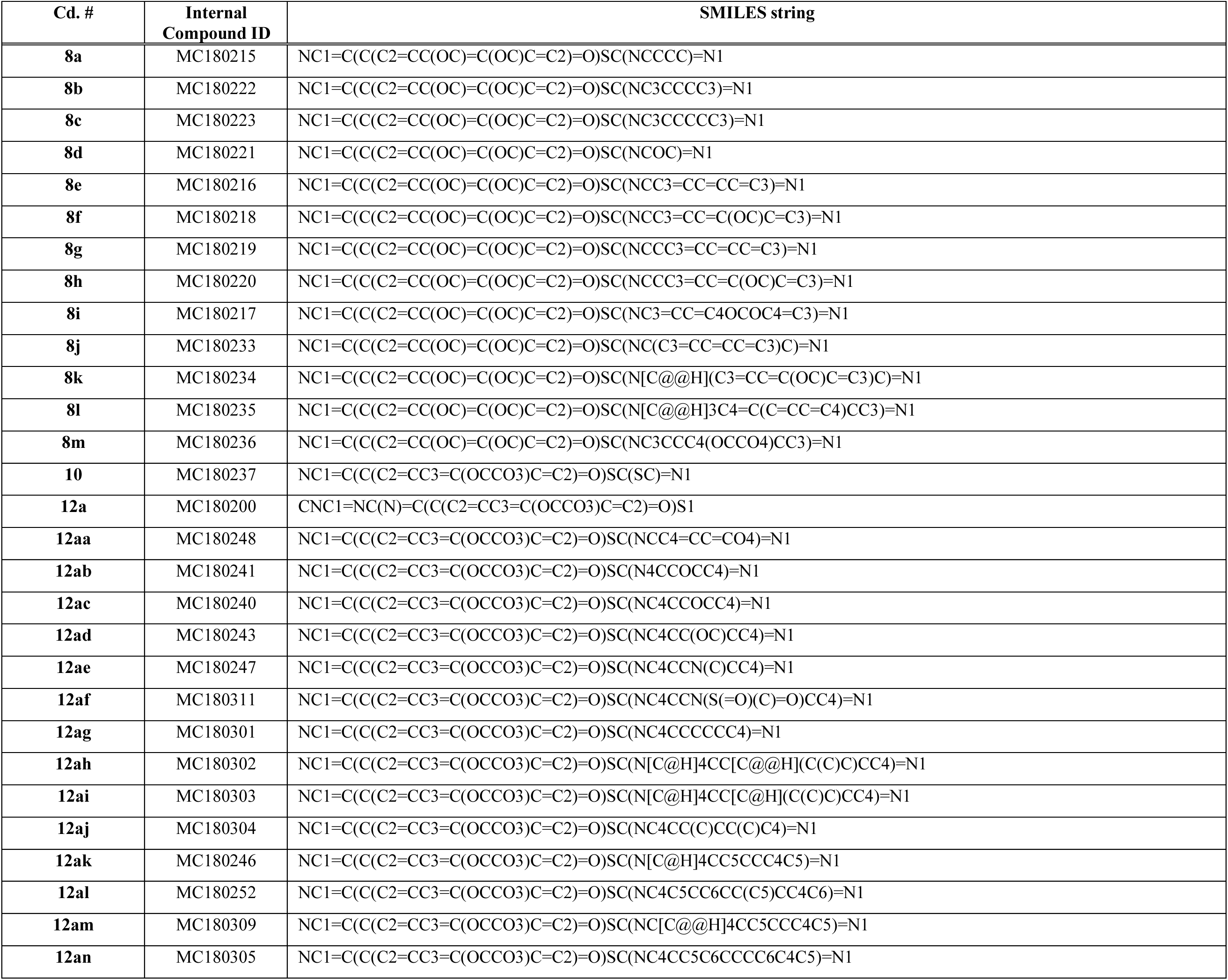
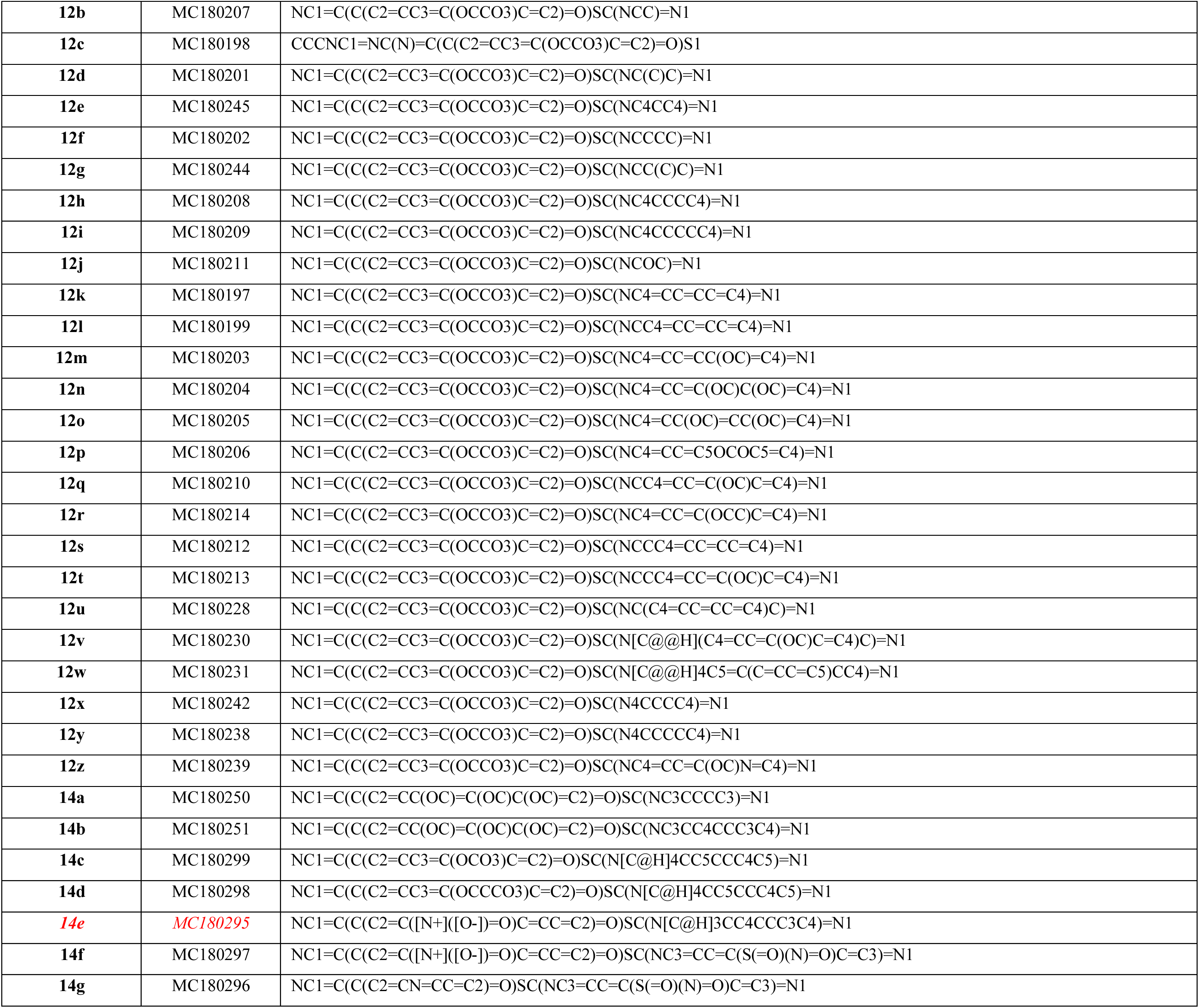
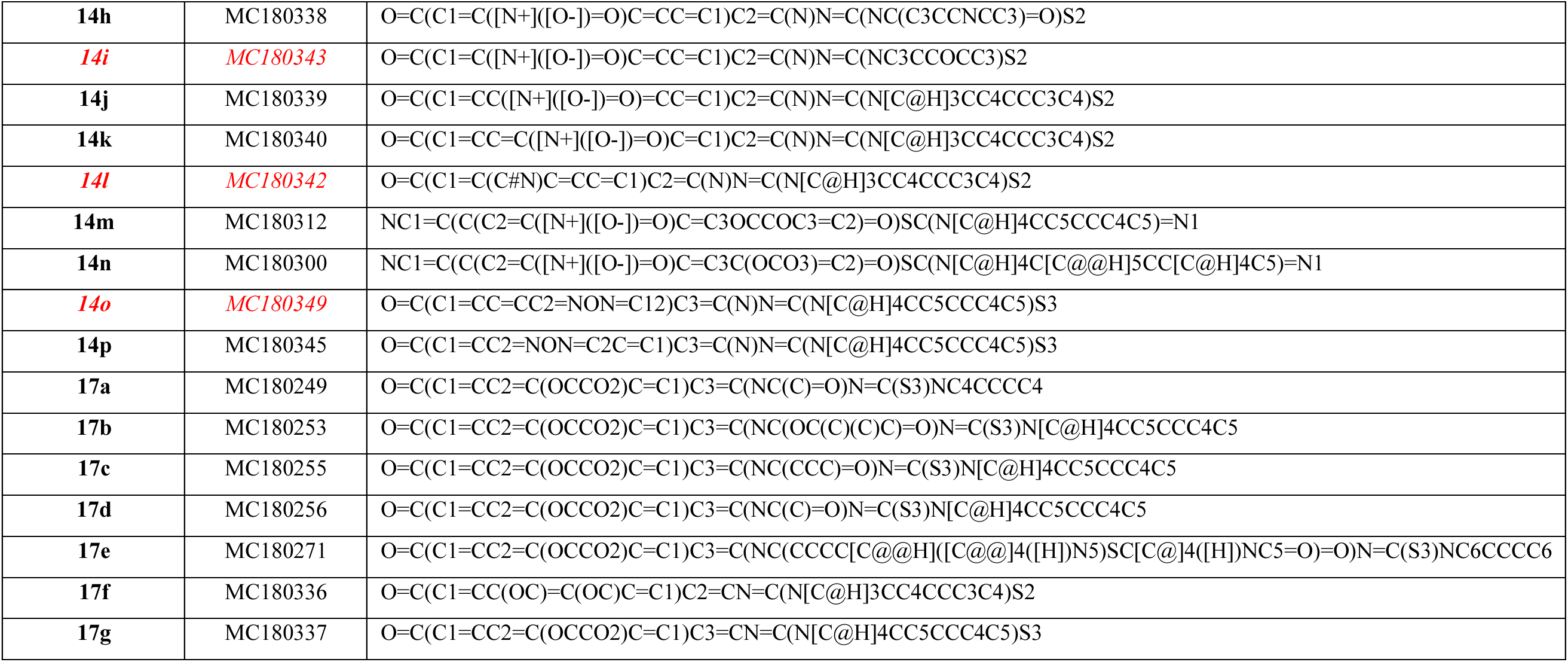
SMILES strings for the 77 compounds described in this study. Compounds highlighted in Figure 5 are shown in *red italics*.

